# AI, citizen science, and the 2024 eclipse emphasize the importance of light for bird behavior

**DOI:** 10.1101/2025.03.14.643188

**Authors:** Liz A. Aguilar, Isaac Miller-Crews, Jeremy M. Dobris, Jo Anne Tracy, Paul Macklin, Shantanu Dixit, Ryan A. Jacobson, Rachel L. Evans, Evan L. McGuire, Daniel P. Beverly, Dustin G. Reichard, Kimberly A. Rosvall

## Abstract

On April 8th 2024, a total solar eclipse disrupted light-dark cycles for North American birds during the lead-up to spring reproduction. Compiling over 10,000 community observations and AI analyses of nearly 100,000 vocalizations, we found that bird behavior was significantly affected by these few minutes of unexpected afternoon darkness. More than half of wild bird species changed their biological rhythms, with many producing a dawn chorus in the aftermath of the eclipse. This natural experiment demonstrates the power of technology-enabled and public science projects to understand our natural world. Further, it underscores the power of light in structuring animal behavior: even when ‘night’ lasts for just four minutes, robust behavioral changes ensue.

## Main Text

Humans have been captivated by total solar eclipses for millennia, and the 2024 Great American Eclipse on April 8 was no exception. April in North America is an especially interesting time for birds, many of which are gearing up for their once-in-a-lifetime chance at reproduction. As tens of millions of people looked to the sky for their own once-in-a-lifetime experience, many wondered how wild birds would respond. After all, birds use light to time important behaviors on a seasonal and daily basis (*1, 2*). Since a total solar eclipse occurs in the same location only once every three or four centuries (*3*), for generations, most free-living birds, like most people, have never seen day turn quickly to night in the middle of day, only to return again minutes later. Leveraging this natural experiment on the role of light in driving bird behavior, we used autonomous recording units, machine learning analyses, and community science observations collected from Mexico to Canada, to quantify how wild birds responded to this unexpected and sudden change in light.

Prior research lends some insight into this question, mainly via localized, anecdotal, or species-specific accounts during a past total solar eclipse. For example, orb-weaving spiders initiated their nightly take-down of their web (*4*), but other arthropods exhibited little to no detectable response (*5*). During a 1932 eclipse, almost 500 reports across New England documented a range of responses, including ‘subdued’ domestic poultry, cattle heading towards their barn, and frogs beginning their dusk chorus (*6*). In an observational study of zoo animals during the August 2017 eclipse, diurnal birds returned to their evening roosts, whereas some nocturnal species exhibited more activity as totality approached (*7*). Also from 2017, audio recordings scored by a trained human showed that this 2.5-minute eclipse changed the rate of vocalization for four bird species in Nebraska, and a computer-assisted soundscape analysis showed an increase in acoustic species richness during and especially just after totality (*8*). Additional big data approaches are starting to use eclipses as a natural experiment to probe the power of light in new ways. For instance, Nilsson and colleagues used doppler radar to monitor biotic airspace at eight locations experiencing totality during the 2017 eclipse, noting decreased activity leading up to totality and more idiosyncratic bursts of activity during totality (*9*). However, the nature of radar does not show exactly which behaviors or species were affected, and there are still no comprehensive analyses of how birds respond to a total solar eclipse.

Community science projects have the potential to resolve these uncertainties because they foster observations at a scale that is not otherwise possible. Such publicly collected data or ‘citizen science’ projects are accelerating discoveries in many fields, from public health (*10, 11*) to environmental science (*12-14*), including ongoing projects on wild birds (*15*). For the April 8, 2024 solar eclipse, totality lasted for only two-to-four minutes, with the moon’s shadow racing roughly 2500 km from Mazatlan to Newfoundland in a couple of hours (**Fig. 1**). To capitalize on this small window of totality and the intense human interest surrounding the eclipse, we created a free smartphone app called *SolarBird*, designed for users with zero birding expertise. We publicized the app through local/national media, schools, and birding groups, ultimately gaining almost 11,000 observations. Users were asked to find a bird and watch it or listen to it for just 30 seconds, clicking up to 10 boxes to record the bird’s behavior (e.g. singing/calling, flying, eating; **Fig. S1**). We analyzed user-recorded bird behavior and time stamps of observations, and the app calculated the sun’s obscuration via GPS coordinates and time stamps (see Materials and Methods).

**Fig. 1.**
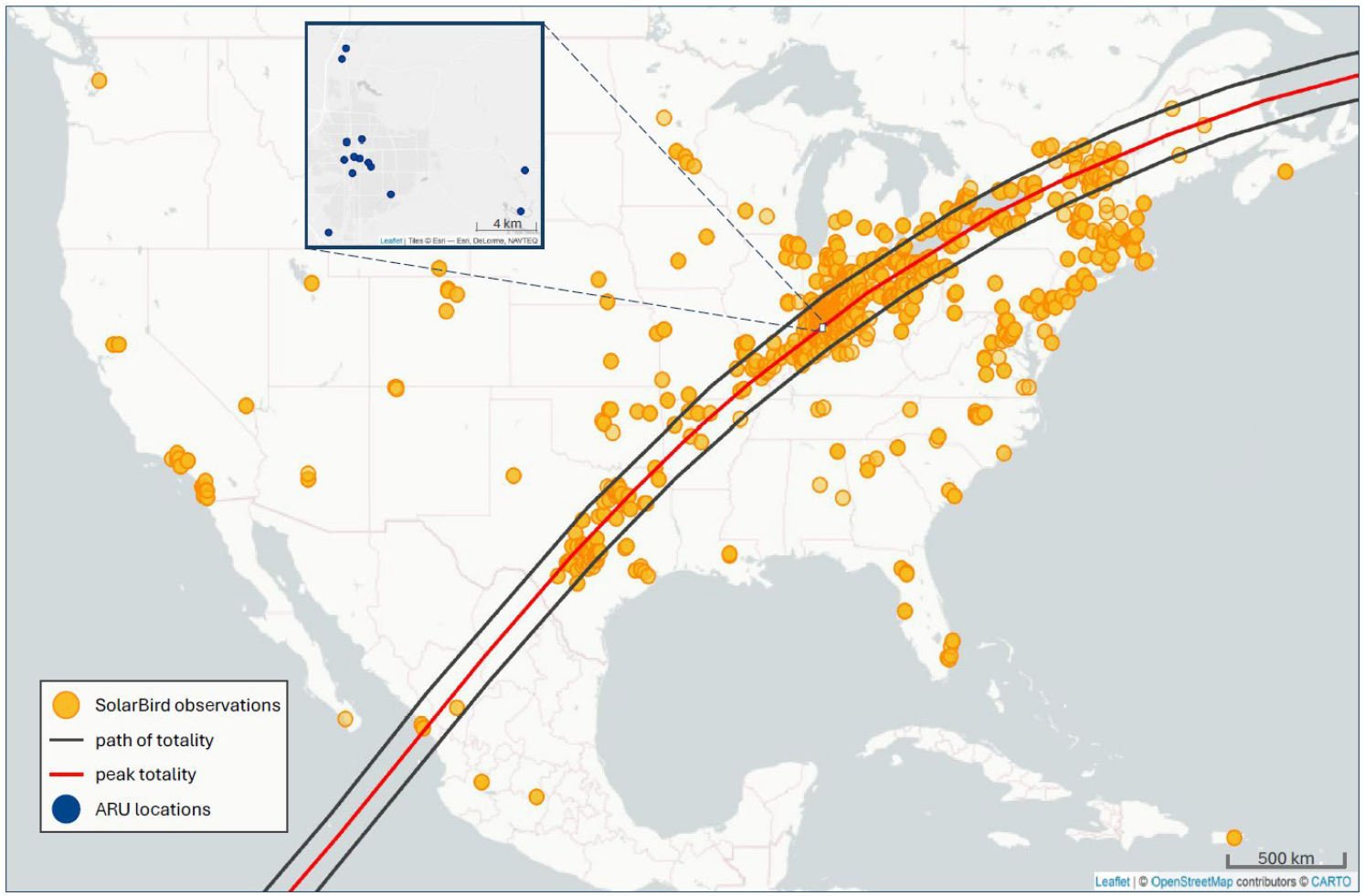
Location of data collection. Orange circles show unique observations from the *SolarBird* app submitted during the April 8, 2024 total solar eclipse. Blue circles (inset from the white box around Bloomington, Indiana, USA) denote locations for Autonomous Recording Units (ARU).

To contextualize observations from the app, we separately audio-recorded avian vocalizations before, during, and after totality. We were especially interested in sound because many birds rely on vocalizations to establish territories and attract mates (*16, 17*), and the timing of the April 8, 2024 solar eclipse overlapped with the annual peak of these behaviors during the lead-up to breeding in the lower Midwest of the United States. Avian vocal output often peaks daily around dawn and dusk (*16*), when light levels change most dramatically, particularly near the spring equinox in temperate regions (i.e., the time and location of this eclipse). Using autonomous recording units, we quantified all birds vocalizing within earshot (*18*) at 14 different locations in southern Indiana (**Fig. 1, inset**). Autonomous recording units have facilitated key insights on vocal behavior and biodiversity in the past (*19*), but recent advances in machine learning have accelerated this process (*20*), allowing scientists to identify avian vocalizations (by species) with limited human oversight (*21*). In particular, the deep artificial neural network BirdNET can now identify over 6,000 species (*21, 22*). Though we are still in the early stages of this technology (*23*), it is already measuring and mapping elements of avian diversity with fewer person-hours than traditional point counts (*24-27*) and therefore accelerating the pace of scientific discovery at scales that were previously infeasible.

We predicted that birds would exhibit dusk-like behavior during totality, mainly ceasing vocalizations and movement, followed by re-initiation of morning behaviors, such as the dawn chorus in which birds produce a burst of song just before, during, or after sunrise (*16*). We evaluated this prediction by leveraging artificial intelligence and community science observations, coupled with standardized focal observations that together provide a new way to experience an event that inspired so many.

## Results and Discussion

### Observations by the public show marked effects on bird behavior

We filtered the total number of observations from *SolarBird* (see Materials and Methods) and focused on 6,951 observations from 1,174 distinct app users who submitted observations before, during, and after totality on April 8 (**Data S1**). Vocalizations (songs or calls) were the most common observation, with users reporting significantly higher rates during totality than in the periods before or after (**Fig. 2a, Table S1**). This pattern is consistent with the burst of vocal behavior that some species exhibit around sunset. In contrast, observations of birds flying, remaining stationary, or exhibiting other behaviors were all documented at significantly lower rates during totality (**Fig. 2bcd, Table S1**). During totality, it is possible that these documented increases in vocalization and decreases in other behaviors may stem from observer bias; after all, light levels during totality were quite low (11.3 ± 1.1 lux, mean ± SD, **Fig. S2**), and observers may have reported what they heard while focusing on the sun’s corona. However, vocalizations and flying were both more commonly reported after totality than before totality (**Fig. 2ab, Table S1**). These differences before vs. after the period of darkness cannot be explained by differences in visibility, and instead they underscore that both behaviors increased in the aftermath of totality, consistent with a burst of activity after sunrise.

**Fig. 2.**
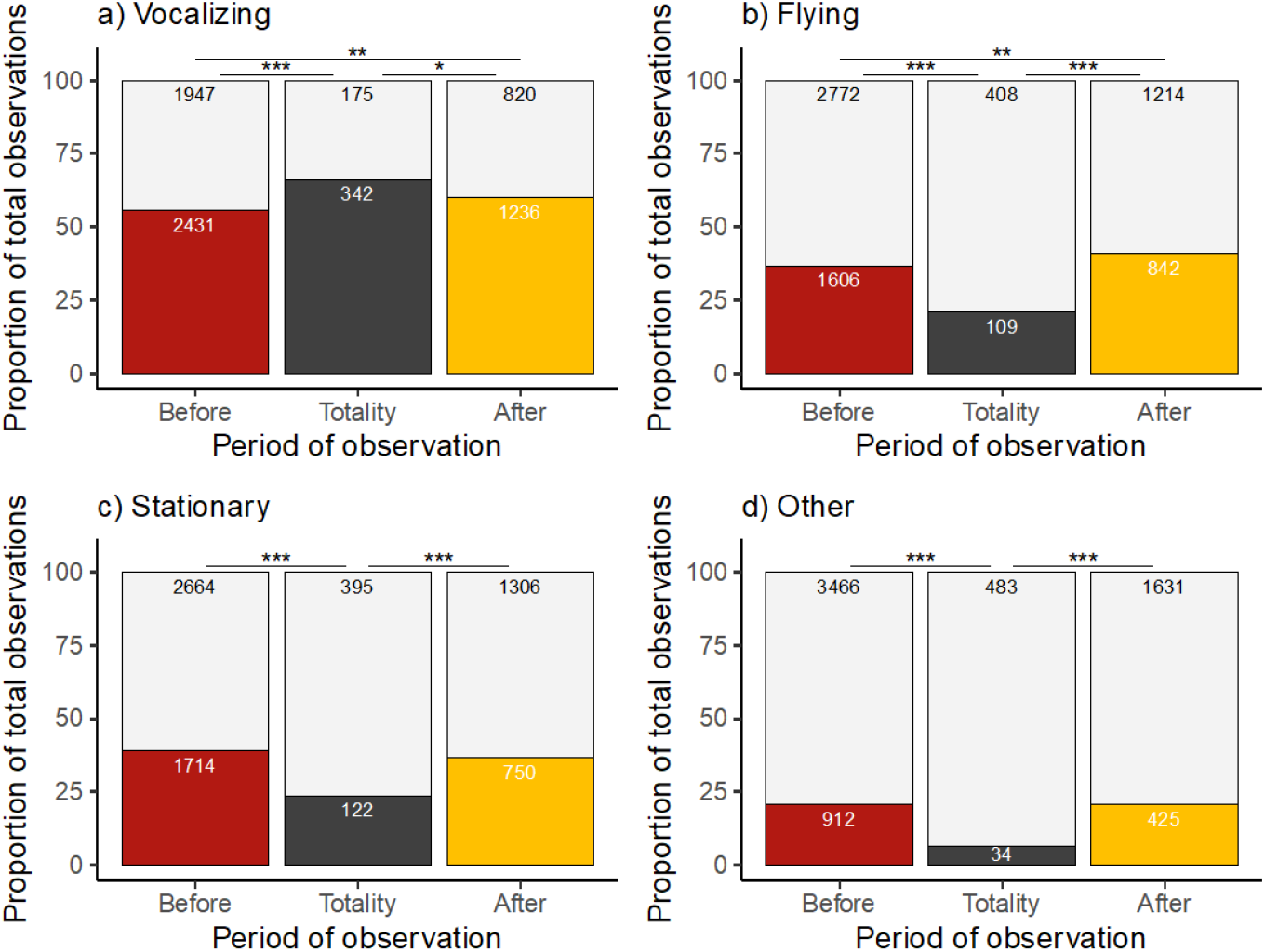
Bird behaviors documented by *SolarBird* app users. Colored bars denote observations submitted on April 8 before totality (“Before”, red), during totality (“Totality”, black), and after totality (“After”, yellow). Reports of vocalizations and flying (a, b) were significantly different between all three periods, while reports of stationary and other (c, d) were significantly lower only during totality. * p < 0.05, ** p < 0.01, *** p < 0.001. Exact p-values reported in Table S1.

### AI shows that some species are especially sensitive to light

Our own recordings also documented marked changes in bird vocalizations surrounding the eclipse. Across 14 locations near Bloomington, Indiana (39.16°N, 86.53°W), we detected 52 species on the afternoon of April 8, 2024 (threshold for inclusion: high confidence and >10 unique detections across at least 5 recorders; see Materials and Methods, **Data S2**). Totality began at approximately 15:05 Eastern Time in these 14 locations (mean starting time: 15:04:55 Eastern Time; see Materials and Methods) and lasted for about four minutes, and so we focused on the 60 minutes before and after totality. After processing these 124 minutes with BirdNET (see Materials and Methods), we detected 15,606 vocalizations from these 52 species (**Fig. 3, Data S3**). This total was 20-50% more than what we detected during the same 124 afternoon minutes on the day before (10,322) or after (13,008). For a subset of species, we validated these AI detections with an expert birder (ELM), who listened to excerpts of the audio files and scored them by ear using the event-recording software JWatcher v1.0 (*28*). Human-scored counts and AI-derived counts from these same time periods were significantly correlated (GLM: *t*(18) = 3.588, *p* = 0.002, *d*^*2*^ = 0.419; **Fig. S5b**). BirdNET detected fewer vocalizations than the trained and consistent human (**Fig. S5)**, but BirdNET was nevertheless a proportional reflection of vocalizations scored by a human.

**Fig. 3.**
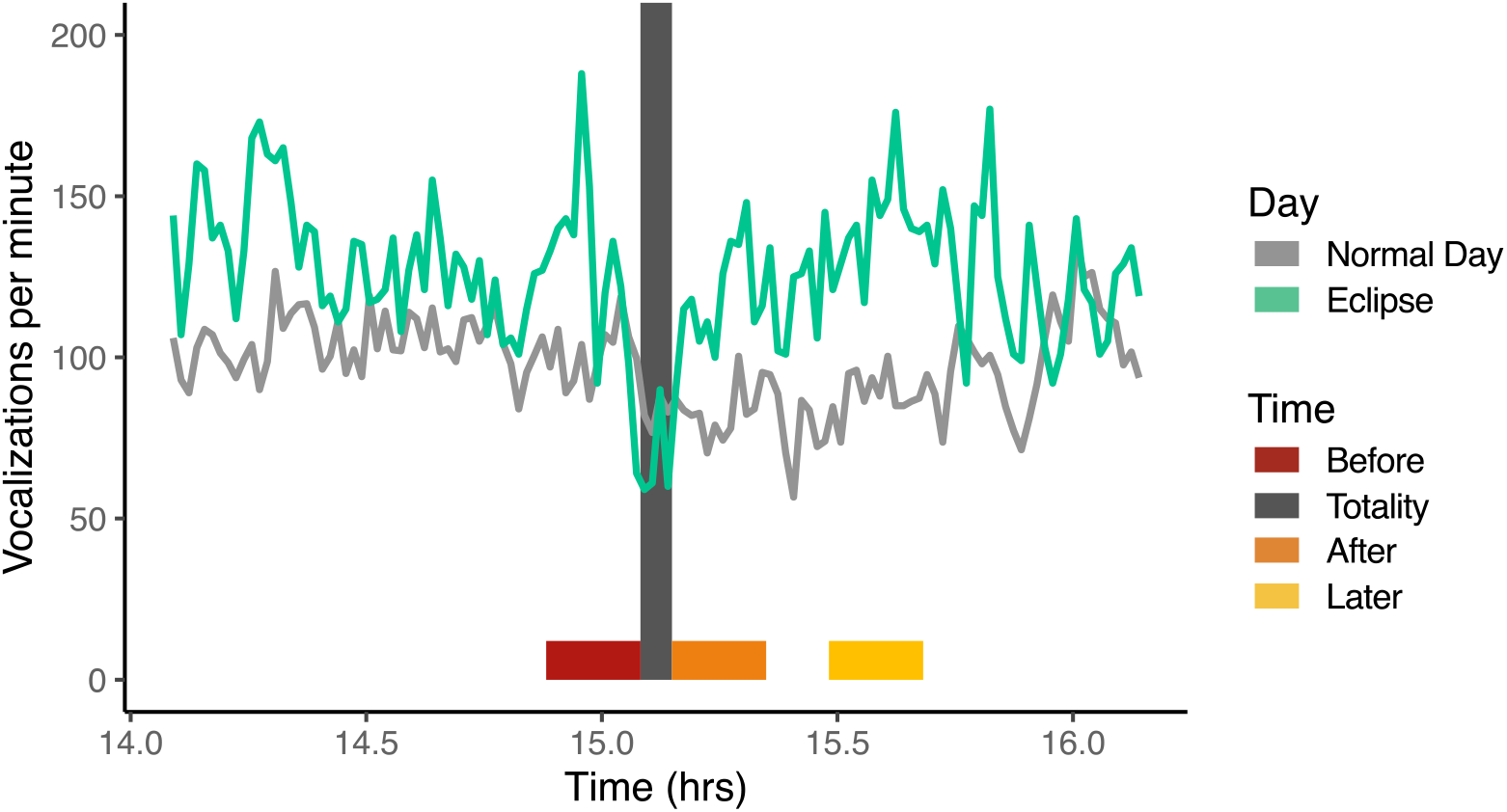
Vocalizations per min, in the hour ± totality. Total vocalizations summed across 52 bird species detected by BirdNET, on the afternoon of the eclipse (teal) and a normal day (light gray). Normal day data are the average number of vocalization drawn from three afternoons surrounding the eclipse day (see Materials and Methods); these data were the source for our formal analyses (see Fig. 4), which focused on species-level effects and used four scoring windows: 12 min just before totality (“Before”, red); 4 min during totality (“Totality”, dark gray), 12 min just after totality (“After”, orange); and 12 min slightly later when a delayed dawn chorus might be expected (“Later”, yellow).

Responses to the eclipse were not uniform among species. Using a Bayesian approach trained on the afternoons of April 6, 7, and 9 (**Fig. S3, Data S3**), we found that 29 of 52 focal species were affected in at least one period of time (**Fig. 4**). In the 12 minutes just before the eclipse (hereafter “before”), when light levels were especially different from a normal day (see Materials and Methods), we documented significant effects on 11 species, most of which (10 of 11) vocalized at higher rates than usual for that time of day.

**Fig. 4.**
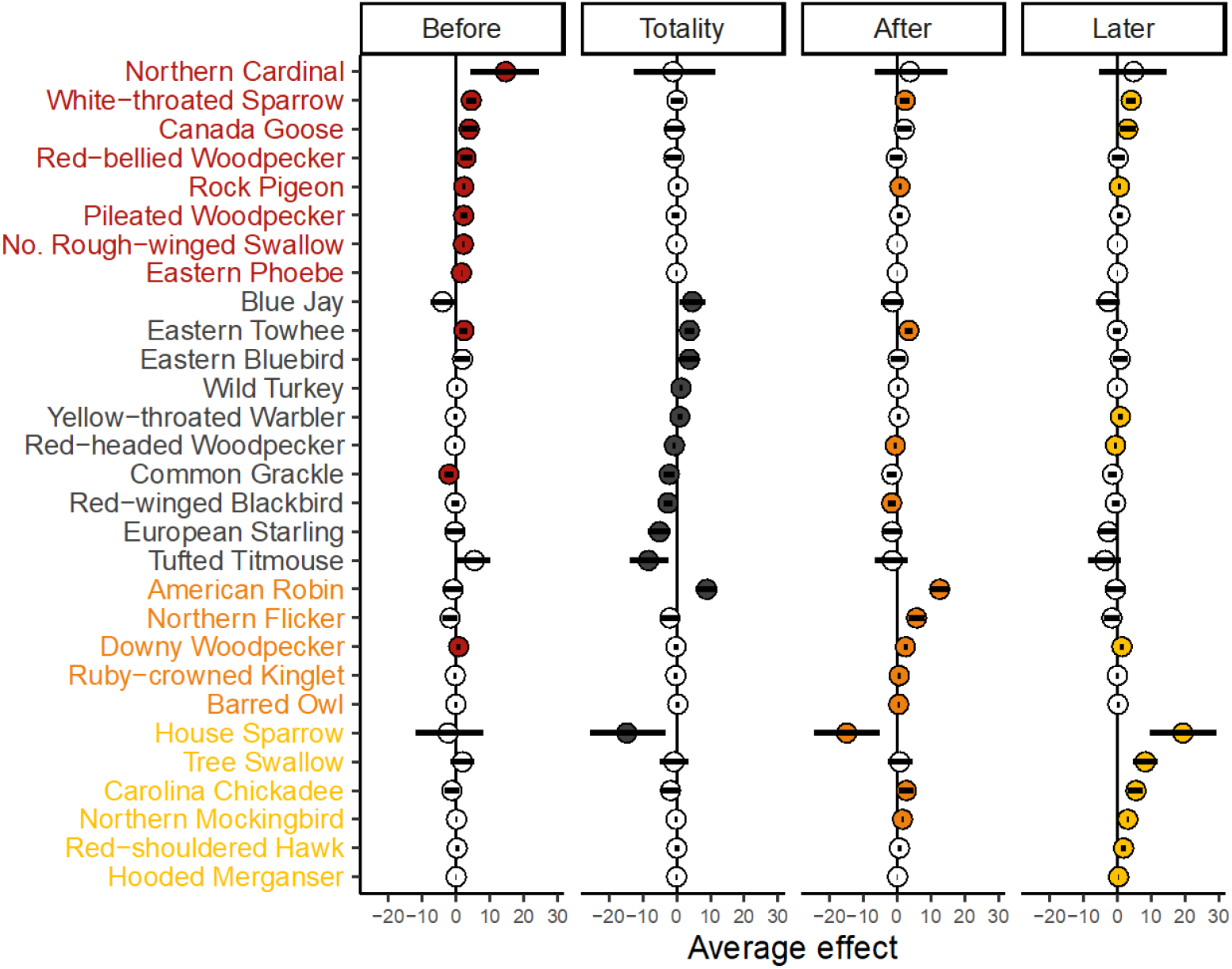
Twenty-nine species responded to eclipse. Species are sorted and colored based on the period in which they were most affected by the eclipse, including 12 min just before totality (“Before”, red), 4 min during totality (“Totality”, black), 12 min just after totality (“After”, orange), and 12 min slightly later when a delayed dawn chorus might be expected (“Later”, yellow). See Fig. 3 for exact sampling times and Materials and Methods for further justification of sampling periods. Each point represents an effect size (scaled per minute) with 95% CI, colored if significant during that particular time period. All 52 species, including those that were not significantly affected by the eclipse, are shown in Fig. S4.

During totality, 12 species vocalized at unusual rates, with half decreasing and the other half increasing, relative to a normal afternoon. American robins (*Turdus migratorius*) displayed the strongest immediate increase during the eclipse (nearly 5x higher than a typical afternoon). Considering that this species was the most abundant in our recordings and is also ubiquitous across US neighborhoods, parks, and cities, it is possible that the robin’s vocal burst may at least partially account for the significant increase in vocalizations reported by app users too.

The largest number of species (n=19) significantly adjusted their vocalization rate after totality ended, either in the 12 minutes immediately after totality (“after”) or the 12 minutes centered around 30 minutes after totality when a delayed dawn chorus might be expected (“later”). Nearly all post-eclipse changes were significant increases relative to a non-eclipse afternoon, consistent with a pseudo-dawn chorus. Notably, barred owls (*Strix varia*) produced 4x as many calls as a normal afternoon, and American robins, which are notorious for their boisterous predawn chorus, vocalized at 6x their normal rate just after totality.

### Dawn chorus behavior is repeatable and predicts eclipse behavior

Next, we asked whether dawn and dusk behavior predicted the type of response that birds had immediately before or during totality (akin to dusk) as well as immediately after totality or ∼30 minutes later (akin to dawn). Using the two most recent dusks (April 6 and 7) and dawns (April 7 and 8), we used BirdNET to score 2-hour periods centered on the transition between civil twilight and sunrise or sunset. Using all the same parameters and thresholds as the main eclipse analysis, this yielded 47,932 more vocalizations (**Data S2, Data S3**), which we found to be consistent in their general onset and offset times per species (**Fig. S6**). For instance, American robins and barred owls vocalize most when it is still dark out, tapering by civil twilight in the morning and picking up again near civil twilight in the evening (**Fig. S3**). Eastern towhees and European starlings exhibit a burst of vocalization in the morning around civil twilight and sunrise, respectively, but in the evening around sunset, only towhees increase their vocalization (**Fig. S3**). Still other species vocalize continuously or off-and-on throughout a typical day, with no crepuscular peak (e.g. house sparrow, northern cardinal; **Fig. S3**). These findings reinforce common knowledge among birders, that each species has a relatively stereotyped vocal response to naturally occurring changes in light. They also provide further evidence that AI can be applied to community-level questions that previously required substantial person-hours (*28, 29*) or were limited to just a few species (*30, 31*).

We operationalized the dawn (or dusk) chorus as ‘present’ if the species’ per minute morning (or evening) peak showed a 100% increase relative to the afternoon (**Fig. S7, Fig. S8**) and used binomial regressions to assess whether dawn or dusk behavior predicted eclipse behavior (see Materials and Methods). Considering that species with a robust chorus naturally display the greatest behavioral shift associated with everyday changes in light, we expected them to be overrepresented in the set of species affected by the eclipse. This prediction was not supported for dusk behavior, which was not predictive of eclipse behavior in the 12 minutes before (GLM: *z*(51)=1.158, *p* = 0.25) or four minutes during totality (GLM: *z*(51)=0.951, *p* = 0.34). However, species with a dawn chorus were more likely than expected by chance to exhibit a significant change in vocalization in the 12 minutes after totality (GLM: *z*(51)=2.023, *p* = 0.04) but not in the 12 minutes centered around 30 minutes after totality (GLM: *z*(51)=0.458, *p* = 0.65). In essence, even though the false night only lasted for four minutes, species that naturally produce a dawn chorus immediately began a new dawn chorus when the false night ended.

### Implications

Changes in light surrounding dawn and dusk are thought to be the primary cue structuring circadian changes in behavior for most organisms (*32*). Here, we show that, when the light-dark cycle was altered by a total solar eclipse, entrained biological rhythms were disrupted for over half of the observed wild bird species. This natural experiment – which essentially switched off the sun for four minutes in the middle of an otherwise normal spring afternoon – is unique for several reasons. For one, light experiments often start from a constant light or constant dark control (*32, 33*) in laboratory environments, the latter of which may not recapitulate natural behaviors and physiology (*34*). The total solar eclipse, however, offers a large-scale manipulation that is both naturalistic and decoupled from time of day. Our results, from both AI- and app-derived data, underscore that the return of light is an especially salient cue, even when that night-to-day switch occurs rapidly at a completely unexpected time (half a day too early) and after an exceptionally short ‘night’ (0.5% the length of the previous night). Eclipse-associated changes in light are also decoupled from some other major environmental shifts that accompany ordinary mid-day changes in light (e.g. heavy rainclouds), though atmospheric pressure, temperature, and even plant photochemistry do change surrounding a total solar eclipse (*35*) and may have played a role here. The fact that total vocalizations were generally high in the hour preceding totality (**Fig. 3**) suggests that birds are more sensitive to changes in light or other atmospheric shifts than are humans, most of whom do not detect the steady decrease in solar illuminance until closer to totality.

The reverse experiment – artificial light at night, or ALAN – disrupts everything from stress physiology to mating behavior to biodiversity, mediated via sometimes subtle changes in biological rhythms (*36-38*) and ensuing effects on melatonin (*33, 39*) or other elements of neurophysiology (*40*). With increasing urbanization and ALAN levels of just a few lux affecting ecosystems across the globe (*33*), there is an urgent need to understand how and why some organisms are more affected than others. Our results provide experimental evidence of marked interspecific differences in the timing and degree of light sensitivity (**Fig. 4**), including 23 species whose vocalization rate was not significantly affected by the eclipse (**Fig. S4**). Though it is possible some species may have switched the nature of their vocalizations (e.g. from mating calls to alarm calls), their otherwise behavioral insensitivity implies sensory mechanisms that maintain biological rhythms despite unusual changes in light. Have they beneficially adjusted to depend on other cues? Or might ALAN maladaptively select for organisms to ignore unusual changes in light to some degree? Though we do not yet know if the mechanisms controlling unusual darkness during an eclipse day are the same as those responding to unusual light during an urbanized night, we urge further research on these species as we seek to mitigate ALAN impacts on wildlife.

By combining AI with community science, we were able to collect and analyze large datasets at both local and continental scales. These powerful approaches are frequently applied to studies of biodiversity (*41, 42*) and more recently to behavior (*31, 43-46*). Our study highlights their underused ability to sample behavioral responses to natural or human-induced experiments, particularly those that occur irregularly over space or time (e.g. wildfire, heatwave, or pollution event). Together, the two approaches enrich one another. For example, the app results show that changes in vocalizations were accompanied by changes in movement (**Fig. 2**), and elements of our localized data apply at a continental scale. In turn, AI provides fine-scale, species-level results (**Fig. 4**) that expand and clarify the public’s observations in the app (**Fig. 2**). Although AI is an imperfect observer requiring human validation (*22*), it extracted data from dozens of species across 196 hours of acoustic recordings, in minutes. This technique allowed us to sample far more species and more time than otherwise possible, opening our eyes to the remarkable effects of light on bird communities. Moving ahead through the Anthropocene, embracing technologies like those employed here promises to lower the bar for community engagement – even for casual observers. Along the way, we can deepen society’s first-hand experience with science and our understanding of the natural world.

## Acknowledgments

We are grateful to Mary Clapp and Elizabeth Derryberry for advice during early phases of this work, to Todd Freeberg for lending additional recorders, to Caty Pilachowski for the initial vision of *SolarBird* and early support, to Teresa Mackin and IU Press for support with app distribution, and to all app users for sharing their eclipse experience with us (listed by name, with permission, in Supplementary Text).

## Funding

NASA Indiana Space Grant Consortium F90002188.02.416 (LAA, KAR) Ohio Wesleyan University (DGR)

National Institutes of Health ‘Common Themes in Reproductive Diversity Training Fellowship 2T32HD049336-16A1 (to IMC)

National Science Foundation IOS-CAREER 1942192 (KAR), with REU support (JD)

National Science Foundation Graduate Research Fellowship (LAA)

United States Department of Agriculture—National Institute of Food and Agriculture 2023-67012-40083 (DPB)

## Author contributions

Conceptualization and funding acquisition: LAA, JAT, IMC, DGR, KAR

App development: LAA, PM, SD, RAJ, JAT

App validation and user interface design: SD

AI validation and analyses: JMD, IMC, KAR

Implementation in the field: RLE, ELM, DGR, DPB, KAR

Formal analysis and visualization: IMC, LAA, JMD, SD, KAR

Writing – original: KAR, DGR, LAA

Writing – review & editing: all

## Data and materials availability

All data are available in the main text and/or supplementary materials, with scripts available at github.com/imillercrews/Eclipse_bird and source code for *SolarBird app* development available at https://github.com/MathCancer/Solarbird.

## Supplementary Materials

Materials and Methods

Supplementary Text

Figs. S1 to S9

Tables S1 to S2

References (47–50)

## References and Notes

1. K. J. Gaston, J. P. Duffy, S. Gaston, J. Bennie, T. W. Davies, Human alteration of natural light cycles: causes and ecological consequences. Oecologia 176, 917–931 (2014).

2. T. D. Williams, Physiological Adaptations for Breeding in Birds. (Princeton University Press, Princeton, NJ USA, 2012).

3. NASA.

4. G. W. Uetz et al., Behavior of Colonial Orb-weaving Spiders during a Solar Eclipse. Ethology 96, 24–32 (1994).

5. M. L. Ferro, Beetles collected in light-traps during the North American total solar eclipse of 21 August 2017 with a review of arthropod behavior during solar eclipses. Entomol News 129, 348–359 (2020).

6. W. M. Wheeler, C. V. MacCoy, L. Griscom, G. M. Allen, H. J. Coolidge, Observations on the behavior of animals during the total solar eclipse of August 31 1932. PNAS 70, 33-70 (1935).

7. A. Hartstone-Rose et al., Total Eclipse of the Zoo: Animal Behavior during a Total Solar Eclipse. Animals 10, 587 (2020).

8. E. M. B. Buckley et al., Assessing biological and environmental effects of a total solar eclipse with passive multimodal technologies. Ecological Indicators 95, 353–369 (2018).

9. C. Nilsson, K. G. Horton, A. M. Dokter, B. M. Van Doren, A. Farnsworth, Aeroecology of a solar eclipse. Biology Letters 14, (2018).

10. J. R. B. Palmer et al., Citizen science provides a reliable and scalable tool to track disease-carrying mosquitoes. Nature Communications 8, (2017).

11. G. N. Bratman et al., Nature and mental health: An ecosystem service perspective. Science Advances 5, (2019).

12. D. Fraisl et al., Citizen science in environmental and ecological sciences. Nature Reviews Methods Primers 2, (2022).

13. J. K. Sheard et al., Emerging technologies in citizen science and potential for insect monitoring. Phil Trans Royal Soc B 379, (2024).

14. D. C. McKinley et al., Citizen science can improve conservation science, natural resource management, and environmental protection. Biol Conserv 208, 15–28 (2017).

15. B. L. Sullivan et al., eBird: A citizen-based bird observation network in the biological sciences. Biol Conserv 142, 2282–2292 (2009).

16. C. A. Staicer, D. A. Spector, A. G. Horn, The dawn chorus and other diel patterns in acoustic signaling. Ecology and evolution of acoustic communication in birds, 426–453 (1996).

17. C. K. Catchpole, P. J. B. Slater, Bird song: biological themes and variations. (University Press, Cambridge, 1995).

18. D. J. Mennill, Field tests of small autonomous recording units: an evaluation of in-person versus automated point counts and a comparison of recording quality. Bioacoustics 33, 157–177 (2024).

19. S. Ross et al., Passive acoustic monitoring provides a fresh perspective on fundamental ecological questions. Funct Ecol 37, 959–975 (2023).

20. D. Tuia et al., Perspectives in machine learning for wildlife conservation. Nature Communications 13, (2022).

21. S. Kahl, C. M. Wood, M. Eibl, H. Klinck, BirdNET: A deep learning solution for avian diversity monitoring. Ecological Informatics 61, (2021).

22. C. M. Wood, S. Kahl, Guidelines for appropriate use of BirdNET scores and other detector outputs. Journal of Ornithology 165, 777–782 (2024).

23. C. Pérez-Granados, BirdNET: applications, performance, pitfalls and future opportunities. Ibis 165, 1068–1075 (2023).

24. S. S. Sethi et al., Large-scale avian vocalization detection delivers reliable global biodiversity insights. Proceedings of the National Academy of Sciences 121, e2315933121 (2024).

25. Y. Lavner, R. Melamed, M. Bashan, Y. Vortman, The bioacoustic soundscape of a pandemic: Continuous annual monitoring using a deep learning system in Agmon Hula Lake Park. Ecological Informatics 80, (2024).

26. B. Hack, C. A. Cansler, M. Z. Peery, C. M. Wood, Fine-scale forest structure, not management regime, drives occupancy of a declining songbird, the Olive-sided Flycatcher, in the core of its range. Ornithological Applications, (2023).

27. L. Bielski, C. A. Cansler, K. McGinn, M. Z. Peery, C. M. Wood, Can the Hermit Warbler (Setophaga occidentalis) serve as an old-forest indicator species in the Sierra Nevada? J Field Ornithol 95, (2024).

28. R. D. Alquezar, R. H. Macedo, J. Sierro, D. Gil, Lack of consistent responses to aircraft noise in dawn song timing of bird populations near tropical airports. Behav Ecol Sociobiol 74, 88 (2020).

29. A. Bruni, D. J. Mennill, J. R. Foote, Dawn chorus start time variation in a temperate bird community: relationships with seasonality, weather, and ambient light. Journal of Ornithology 155, 877–890 (2014).

30. D. Amorós-Ausina, K.-L. Schuchmann, M. I. Marques, C. Pérez-Granados, Living Together, Singing Together: Revealing Similar Patterns of Vocal Activity in Two Tropical Songbirds Applying BirdNET. Sensors 24, 5780 (2024).

31. C. M. Wood, S. Kahl, S. Barnes, R. Van Horne, C. Brown, Passive acoustic surveys and the BirdNET algorithm reveal detailed spatiotemporal variation in the vocal activity of two anurans. Bioacoustics 32, 532–543 (2023).

32. R. G. Foster, S. Hughes, S. N. Peirson, Circadian Photoentrainment in Mice and Humans. Biology 9, 180 (2020).

33. M. Grubisic et al., Light Pollution, Circadian Photoreception, and Melatonin in Vertebrates. Sustainability 11, 6400 (2019).

34. R. M. Calisi, G. E. Bentley, Lab and field experiments: Are they the same animal? Horm Behav 56, 1–10 (2009).

35. D. P. Beverly et al., Hydraulic and photosynthetic responses of big sagebrush to the 2017 total solar eclipse. Scientific Reports 9, 8839 (2019).

36. K. J. Gaston, T. W. Davies, S. L. Nedelec, L. A. Holt, in Annual Review of Ecology, Evolution, and Systematics, Vol 48, D.J. Futuyma, Ed. (2017), vol. 48, pp. 49-+.

37. U. Tuomainen, U. Candolin, Behavioural responses to human-induced environmental change. Biological Reviews 86, 640–657 (2011).

38. K. J. Gaston, A. Sánchez de Miguel, Environmental Impacts of Artificial Light at Night. Annual Review of Environment and Resources 47, 373–398 (2022).

39. G. C. Brainard, B. A. Richardson, L. J. Petterborg, R. J. Reiter, The effect of different light intensities on pineal melatonin content. Brain Res 233, 75–81 (1982).

40. O. J. Marston et al., Circadian and dark-pulse activation of orexin/hypocretin neurons. Molecular Brain 1, 19 (2008).

41. C. M. Wood et al., A scalable and transferable approach to combining emerging conservation technologies to identify biodiversity change after large disturbances. J Appl Ecol 61, 797–808 (2024).

42. M. Chandler et al., Contribution of citizen science towards international biodiversity monitoring. Biol Conserv 213, 280–294 (2017).

43. D. Sossover, K. Burrows, S. Kahl, C. M. Wood, Using the BirdNET algorithm to identify wolves, coyotes, and potentially their interactions in a large audio dataset. Mammal Research 69, 159–165 (2024).

44. G. M. Leighton, J. P. Drury, J. Small, E. T. Miller, Unfamiliarity generates costly aggression in interspecific avian dominance hierarchies. Nature Communications 15, 335 (2024).

45. E. T. Miller et al., Fighting over food unites the birds of North America in a continental dominance hierarchy. Behav Ecol 28, 1454–1463 (2017).

46. R. Kaartinen, B. Hardwick, T. Roslin, Using citizen scientists to measure an ecosystem service nationwide. Ecology 94, 2645–2652 (2013).

47. D. H. McDougal, P. D. Gamlin, in Comprehensive Physiology. pp. 439–473.

48. K. H. Brodersen, F. Gallusser, J. Koehler, N. Remy, S. L. Scott, Inferring Causal Impact Using Bayesian Structural Time-Series Models. Ann Appl Stat 9, 247–274 (2015).

49. T. Giorgino, Computing and Visualizing Dynamic Time Warping Alignments in R: The dtw Package. Journal of Statistical Software 31, 1–24 (2009).

50. R. Killick, I. A. Eckley, changepoint: An R Package for Changepoint Analysis. Journal of Statistical Software 58, 1–19 (2014).

