## Supplemental Material for "AI, citizen science, and the 2024 eclipse emphasize the importance of light for bird behavior"

**The PDF file includes:**

Materials and Methods
Supplementary Text
Figs. S1 to S9
Tables S1 to S2
References

### Materials and Methods

*SolarBird* app overview and initial processing of observations

*SolarBird* was designed for simplified collection of observations from Apple and Android phones across the path of the eclipse; see below for more details on the app design and implementation. We obtained a total of 10,859 observations from 1,738 users, each of whom were given a unique ID based on their phone. We recorded GPS coordinates, and submissions from outside North America were blocked. Before analysis, we implemented the following data filters: First, we removed observations submitted as practice (a feature in the app), those with missing metadata (e.g., location, time), and those recorded outside the path of totality. Next, we focused only on observations from eclipse day, categorizing them into before, during, or after totality, based on when the observation began in relation to the beginning of totality at the user's GPS position. Totality occurred when the sun was completely obscured by the moon (100%), and all other values of obscuration (0-99%) were treated as before or after totality. On average, observations were submitted 43 minutes before totality (range: 1 min to 9 hrs) and 20 minutes after totality (range: 1 min to 8 hrs). During totality, observations were submitted on average ± 1 minute from peak totality, with totality lasting an average of 3 minutes 42 seconds at the user's location. The app calculated the duration of the observation, and we kept only those within the range of 5-60 seconds; this filter eliminated entries that were either too short to be reliable or too long to match to a specific time entry (before, during, or after totality). This left 6,951 observations from 1,174 unique users, used in our analyses (**Data S1**). To better categorize related behaviors from among the 10 possible boxes, we combined 'perched alone,' 'perched with others,' and 'sitting' into a single 'stationary' category. We also combined user observations of 'hearing' and 'singing' into a single 'vocalizing' category. Due to low counts, we combined 'eating,' 'swimming,' 'walking,' and 'other' into 'other'. To compare the proportion of behaviors observed before, during, and after totality, we used binomial general linear models with post-hoc Tukey tests with FDR correction (**Fig. 2**; **Table S1**).

Autonomous recording units and locations

We used Song Meter SM4 Acoustic Recorders (Wildlife Acoustics, Maynard, MA, USA), with current firmware and synchronized to the same clock using a GPS connection prior to placement in the field. We recorded uncompressed audio (.WAV) at a sample rate of 24 kHz in stereo with a gain setting of 16 dB for each channel. To conserve battery life and data storage, we limited recordings to 75 minutes before and after both sunrise and sunset as well as a three-hour period surrounding the eclipse totality (13:30-16:30) each day. Systematic recording began at dawn on April 2 and ended at dusk on April 9. We placed recorders in open habitat (i.e. not forests or woodlands), though sites ranged from completely rural (wetlands, pasture) to suburban to more urban sites around Bloomington, Indiana (population ~150,000).

Our eclipse recordings focused on data collected during totality, plus one hour before and one hour after. Because sites were up to 14.4 km apart from one another (**Fig. 1**), the exact time of totality varied slightly among sites, starting between 15:04:47 to 15:05:04 local time and lasting for 4m00s to 4m03s minutes. For simplicity and to allow for subtle variation that could stem from nearby structures, vegetation, or topography, we treated all sites as having a 04m00s period of totality, beginning at the midpoint of the start of totality (15:04:55).

To quantify typical behavior surrounding dawn, dusk, and 3PM local time, we used recordings from April 6, 7, and 9. We did not extend beyond these few days because of the rapid temporal changes that occur during the ramp up to spring breeding. In addition, April 6-9 were all cool spring days (mostly sunny to mostly cloudy), without inclement weather that could interfere with either behavior or our ability to detect it.

Quantifying behavior in BirdNET

BirdNET processes audio by dividing files into 3-second segments. By default, the segments do not overlap, and each segment begins when the previous one ends. For each segment, the software produces a list of the species identified and counts the number of times identified. By using this default overlap of “0” seconds, we minimize issues of double-counting songs that cross between two segments, though we should also miss vocalizations that are bisected by the 3-second time bins. We uploaded our audio files without prior filtering because we found during parameter testing that unfiltered files yield more detections, so we used the default frequency range of 0-15kHz. We initially used BirdNET to generate a custom list of focal species based on the location of the recorders and the date of the eclipse, but filtered this possible list from 142 species to only 52 species that were detected at least 10 times across at least 5 recorders with moderate to high confidence (>0.5) on the afternoon of the eclipse. BirdNET’s confidence scores are unitless measures that range from 0-1. They are not synonymous with a confidence interval, nor are they transferable across species, but they do provide a measure of the neural network’s confidence in its accuracy (*22*), which can be shaped by background noise or the bird’s distance from the recorder (*23*). After removing species that never achieved confidence >0.5, we otherwise used all detections with a confidence score of 0.1 or above because this best fit the human-scored analyses.

*Validation with human*: An expert birder (ELM) listened to excerpts of the audio files, scoring them by ear using the event-recording software JWatcher v1.0 (*28*), to compare to the AI results. Based on the habitat and date, but blind to the AI data, we selected (*a priori*) 36 species (e.g. American robin, song sparrow, downy woodpecker) or species groups (e.g. owls, other sparrows, other woodpeckers) and recorded each vocalization with a unique keystroke (**Table S2**). Results were summed at 3-second intervals to mirror BirdNET’s algorithms. We scored four different 4-minute time windows relative to the start of totality: (a) 0-3 minutes (totality), (b) +4 to +7 minutes (immediately after), (c) +28 to +31 minutes (~30 min after), and (d) 24 hours before, the latter of which represents a control measure of vocal output on a typical (non-eclipse) afternoon in early April. Minutes 0-7 were scored three times each to validate the human-derived score, which was highly repeatable (ICC = 0.98 [0.97-0.99]; **Fig. S5a**). To compare the human-scored data to AI-scored data, we used the total number of bird vocalizations for each species across the four minutes of totality and the four minutes right afterwards, focused on the 20 species that received non-zero counts by the human. Initially, we used a poisson GLM of total song calls per species for AI~Human with 19 degrees of freedom. With this model, over dispersion was high and American Robin was identified as an outlier with a cooks distance of 28. To deal with over dispersion, we changed to a quasi-poisson model, with American Robin removed, and found a significant correlation between human and AI bird vocalization counts (GLM: $t(18) = 3.588$, $p = 0.002$, $d^2 = 0.419$; **Fig. S5b**). Based on this pseudo-$R^2$, 41.9% of the variability in AI counts is explained by its relationship with the human counts. Likewise, based on the effect size of 1.01, a one unit increase in human count corresponds to a 1% increase in AI count.

*Sampling Periods and Analysis*: To focus our main eclipse analyses on the most atypical light conditions, we filtered afternoon data to focus on four main times (**Fig. 3**): (a) the four minutes of totality, (b) the 12 minutes immediately after totality, (c) the 12 minutes immediately before totality, and (d) 12 later minutes centered on the 30 minutes after totality ended. This lattermost time was selected due to the expected delay between the onset of light and the peak of "dawn" vocalizations. Times (b) and (c) were selected because (i) light intensity fell well below normal levels around this time (lux <9k) and (ii) light intensity did not dip below 9k during cloud cover earlier in the day. Typical values for the afternoon of April 8 range from 41k to 96k in the last five years: https://igws.indiana.edu/eclipse#eclipseData. Our measurements on April 8, 2024 showed a peak of almost 100k lux just before partiality began at 1:49PM local time, followed by a steady decline until totality (**Fig. S2**). We do not know exactly how birds perceived this steady change in light, but vertebrate eyes can compensate for subtle changes in visible light, at least until it is dark enough that they cannot further adjust (*47*).

To assess the impact of the eclipse on bird vocalizations, we implemented a Bayesian casual inference approach for time-series data using the R package 'CausalImpact' (*48*) (**Fig. 4, Fig. S4)**. We generated a single pretraining window for all comparisons, the 12-minutes centered around the time totality occurred, using non-eclipse afternoons on April 6, 7, and 9. Vocalizations produced during the four minutes of totality and each of the three 12-minute time windows surrounding the eclipse were compared to the number of vocalizations expected based on the pretraining window. We then FDR corrected within a comparison across all 52 species.

*Dawn and dusk behavior*: To score each species for the presence or absence of a dawn or dusk chorus, we first needed to quantify common knowledge among birders, that each species has a stereotyped timing of vocalization, in association with normal daily changes in light. To do this, we used dynamic time warping (DTW), which is a flexible method for comparing the shape of curves in time-course data. Beginning with vocalization per minute data across each dawn, dusk, or afternoon (**Fig. S3**), we calculated distance between recordings of the same time period across days (i.e. dawn, afternoon, and dusk). We used a Sakoe-Chiba band constraint to allow for some invariance because we expected the pace of vocalization to imperfectly match minute-to-minute, but we wanted to maintain variance if vocalizations were outside of that window size. To set the Sakoe-Chiba band window size, we optimized the value for every pairwise comparison in each species separately. For each pairwise comparison per species (e.g. Dawn of the 6th vs Dawn of the 7th; "Dawn6_7"), we calculated the DTW distance with Sakoe-Chiba window sizes ranging from 1 to 60 minutes using dtwDist from the R package 'dtw' (*49*). We then used a change point analysis to get the first change point using a nonparametric cost function with cpt.np from the R package 'changepoint.np' (*50*). This change point value was then used to calculate the Sakoe-Chiba band window size for that specific pairwise comparison for that species. Though there is no agreed-upon threshold for "consistency" versus "inconsistency," we note that dawns and dusks are highly consistent, with afternoons less so (**Fig. S6).**

Next, we calculated per minute rates of vocalizing across seven key periods of the day, on non-eclipse days, visualized in **Fig. S7** and listed here:

(1) the 45 minutes prior to civil twilight in the morning,
(2) the 30 minutes between civil twilight and sunrise in the morning,
(3) the 45 minutes after sunrise in the morning,

(4) the 124 minutes centered on the would-be time of totality (15:04:55),
(5) the 45 minutes prior to sunset,
(6) the 30 minutes between sunset and civil twilight,
(7) the 45 minutes after civil twilight.

We compared the peak morning rate (from periods 1-3) and peak evening rate (from periods 5-7) to that same species' afternoon rate (period 4). If the rate of increase surrounding dawn or dusk was ≥ 100% (or, a doubling, see **Fig. S8**), we scored that dawn or dusk chorus, respectively, as "present." There is no single best definition of a dawn (or dusk) chorus, other than a burst of vocalization that occurs before, during, or just after sunrise (or sunset). This 100% increase is conservative, focusing our predictions on species with the most robust chorus. To relate the presence/absence of a chorus to the effect of the eclipse, we used binomial regressions, with presence/absence of dawn (or dusk) as the independent variable (0/1) and the eclipse effect in each sampling period as the dependent variable (before, during, after, later; scored as 1 if significant and 0 if not significant). Since most species with a significant effect of the eclipse on vocalizations showed only increases in vocalizing (23/29; 79.3%), and very few species had only decreases (5/29; 17.2%) or a combination of the two (1/29; 3.4%), we did not feel we had enough statistical power for a multinomial logistic regression comparing the directionality of significance. Instead, we focused on whether a species' vocalizations were impacted by the eclipse using a binomial approach.

### Supplementary Text

Additional Detail on the *SolarBird* app

*SolarBird* was written with the Flutter framework (Version 3.16.8; https://flutter.dev/), a cross-platform app development toolkit that allows a unified code base to be used to create apps for iOS (iPhone) and Android. Flutter-based apps store user responses (submitted observations, stored with tailored data elements, namely: udid, observationTime, timeDiff, eclipseType, maxEclipse, timeSubmit, totalBegins, obscuration, flying, hear, perchedAlone, perchedOthers, sing, sitting, swimming, walking, other, eating) in a secured Google Firebase database (https://firebase.google.com/), a real-time NoSQL database.

After extensive internal development and testing, *SolarBird* was released for iPhone / iOS in the Apple App Store and for Android devices in the Google Play Store. After the eclipse, we downloaded all user data from the Firebase database for analysis, and we purged the Firebase database to confirm that no user data were retained in the cloud.

*SolarBird* included features aimed to prepare our community researchers to collect quality data and engage with additional aspects of the eclipse as well as the scientific process. The homepage displayed real-time progression of the eclipse based on the user's GPS coordinates that was supplemented by push notifications as totality approached. In a 'Guide' section, we provided step-by-step instructions for observing a bird, defined focal behaviors in an ethogram, and enabled optional practice submissions so participants could familiarize themselves with data collection prior to the eclipse. There were also reminders for safely viewing the eclipse and not interacting with wild animals. In the 'Info' section of *SolarBird*, we outlined project goals to ensure participants understood the purpose of their contributions and how their efforts as citizen scientists would have critical impacts on these research questions. This section also included general eclipse information that defined terms displayed on the homepage.

Our app design philosophy aimed to balance the goals of gathering scientifically meaningful observations without substantially impacting on their eclipse viewing experiences. The user interface and user interactions described below were chosen to reduce the physical clicks required to submit an observation (to increase likelihood of participation and reduced impact on eclipse viewing) and to reduce the required specificity and complexity in observations (to increase the likelihood of gathering robust measurements from non-expert participants). In the interest of ease-of-use, we did not require the user to identify the species they observed, but rather to click four possible options for its size. Each option also included examples to serve as accessible scales for the layperson: Tiny (house sparrow, tree swallow, eastern bluebird); Small (northern cardinal, blue jay, American robin); Medium (American crow, mallard, barred owl); and Large (turkey vulture, goose, eagle). Since the user may not have visually seen their focal bird and/or were focused on its vocalizations, this step of the observation submission was optional. Out of the 6,951 observations used in our analyses, most users reported watching a small bird (**Fig. S9**), about the size of a northern cardinal, blue jay, or American robin.

*SolarBird participants*: Per our privacy notice (below), app users could optionally share their name and opt into being listed in a publication. We are extremely grateful to:

Abbey Rhein, Abby Publow, Abby Westropp, Abigail Kallay, Abigail Shake, Abigail Weber, Aby Wischmeier, Adam Akin, Adam Raven, Adam Thada, Aden Lawyer, Aidan Burr, Aimee Osborne, Alaina

Hensley, Alan Blanchard, Alec Dorn, Alex Bzdafka, Alex Nahrwold, Alex Sidare, Alexa Gioia, Alexander Comiso, Alexander Gansmann, Alexander Mindock, Alexander Stein, Alexandra Verma, Alexis Klutinoty, Ali Weising Pike, Alisa Devenport, Alisha Stapp, Alison Anderson, Alle Kloss, Alli Gauthier, Allison Lippard, Allison Litmer, Allison Pierce, Aly Van Zyl, Alyssa Beccue, Alyssa Foss, Amalie Orme, Amanda B., Amanda East, Amanda Thornburg, Amanda Vavala, Amarilys Torres, Amber Schiefelbein, Amelia Lobo, Amrith Annamreddy, Amy Blase, Amy Cornell, Amy Downing, Amy Girten, Amy Jessmer, Amy Jones Richardson, Amy Lazoff, Amy Marks, Amy Ollendorf, Amy Piekosz, Amy White, Amy Wolfram, Ana Bacon, Ana Luisa Santo, Andis Berzins, Andrea Lynch, Andrea Mair, Andrew Beauchamp, Andrew Decker, Andrew Hutchison, Andrew Kaizer, Andrew Vandermeer, Andy Belt, Angel Jackson, Angela Riley, Angela Tammen, Angelyn Kensinger, Angie Damm, Angie Parilac, Angie Shelton, Anisah Mihaljevic, Anita Ragan, Ann MacDonald, Ann Maner, Ann Snider, Anna Bruettig, Anna Byers, Anna Chinni, Annalee and SiennaMay , Annalee Friedman, Annamae Harmon, Anne Anderson, Anne Brahaum, Anne Haines, Anne Larr, Annette McClellan, Annie Aguirre, Annie Turnee, Anthony Ramirez, April Bednarski, Araceli Akers, Ariana Huggins, Arnold Israel, Arthur Hertz, Ash Bermam, Ashlee Engelhardt, Ashley Gant, Ashley Johnson, Ashley Parsons, Aubrey Dryden, Audra Loney, Audrey Brumback, Austen Ehrie, Austin Broadwater, Ava Ciaccia, Ava Harris, Bailee Parsons, Bailey Hollweg, Bandy Russell, Barb Brown, Barbara Bennett, Barbara Hawkins, Barbara Ray, Barry Pratt, Beau Nance, Becca Horger, Beck Dalton, Bee Stoll, Belinda Schneider, Ben Duggan, Ben Schrader, Ben Smith, Benjamin Gombash, Benjamin Irvin, Bennett Hanson, Bennett Thompson, Beth Lewer, Beth Miller, Beth Walker, Betty Watson, Beverly Kysar, Bill Davis, Bill Watson, Bob Davids, Bob T, Bob White, Brandi Perkins, Brandie Ruark, Breana Murad, Brenda Chew, Brenda Phillips, Brendan Thomas, Brett Keller, Brian Davis, Brian Ellison, Brian Iezzi, Brian Kelly, Brian Murphy, Brian Rosen, Brian Stokes, Brianna Firkowski, Bridget Cole, Bridget Merrion, Brielle DeCarolis, Brittany Thornburgh, Brittiney Reese, Broderick Parks, Brooke Kile, Brooke Stokdyk, Bryan Fleck, Bubba Mac, Burton Vculek, C Ford, C Peters, Caillie Monrad, Caleb Scholtens, Caleb Smith, Callie, Cameron Yeakle, Camila Carvajal, Camille Knight, Can Wu, Candace Kingma, Cara Righter, Carly Hawkins, Carmalee Treat, Carol Goodall, Carol Mardeusz, Carol Suich, Carole Kulpa, Caroline Gilley, Caroline Lafleur, Carrie Walters, Carsyn Hagans, Cary Albright, Casey Buckley, Casey Coomes, Casey Matczak, Casey Smith, Cassie Childress, Cassie Crawford, Catherine Daligga, Catherine Haag, Catherine Paquet, Cathryn Alderson, Cathy Butterfield, Cathy Kenny, Cathy Walz, Cece Wong, Cedric Harris, Chad Herrenbruck, Chandler Carr, Chanel Williams, Charci Causey, Charity Robertson, Charlotte Griffiths, Charlotte Reemts, Charlotte Toombs, Chelsea McDonnough, Cheryl Rizzo, Cheryl Wall, Chris Armstrong, Chris Enderle, Chris Goldrick, Chris Jordan, Chris Padgett, Chris Shelton, Chris Tebbens, Chris Usher, Christie Debelius, Christina Lawson, Christina Sargent, Christina Vendely, Christine Davis, Christine Lattin, Christine Schumacher, Christopher Rotunno, Christopher Shaw, Christy Hyman, Christy Lorenzoni, Christy Scheid, Cindy Grant, Claire Jones, Clarissa Zore, Clark Richter, Clementine Bodin, Connie Richardson, Constance Ober, Courtney McManus, Courtney Meyer, Craig Endres, Craig Stephenson, Cristin Hagans, Crystal Stein, Crystal Thompson, Cyndra Pilkington, Cynthia Collie-Lunsford, Cynthia Crawford, D Naluzny, D Steitz, Daisy Spalding, Dan Calarco, Dan Cristea, Dan Lambert, Dan Mahoney, Dan Reed, Dana Coplin, Dana Jarboe, Dana Newell, Dani Ansaldo, Daniel C, Danielle Carmichael, Danielle Schroeder, Darleah Winsted, Darren Johnson, Dave Prentice, Dave Worthington, David Castillon, David Kruse, David Orr, David Polly, David Rodgers, David White, David Wolowitz, Dawn Hewitt, Dawn Martin, Dawn Terry, Dean Weinkein, Deb Loll, Deb Reed, Debbe Staggs, Debbie Snider, Debby Herbenick, Deborah Blue, Deborah Cohen, Deborah Crowe, Deborah Dohne, Deborah Perry, Deborah Sanders, Deborah Stukenborg, Debra Atkins, Debra Potts, Delana Lewis, Delaney Demster, Denise Gardiner, Denise Girolamo, Dennis Johnson, Dennis Mishler, Deon Allen, Derek Worch, Devin Dempsey, Dewanna Lindo, Diana Kenney, Diane Campanello, Diane Howe, Diego Morales, Dimitar Nikolov, Dinah Cohen, Dionna McClure, Dong-gyu Kim, Donna Bernens-Kinkead, Donna Graber, Donna Spalding, Dora George, Doreen Deutsch, Dorothy Rich, Douglass Gaking, Dulce Ramos, Dylan Anderson, E. Smout, Edith Irvin, Eileen Doherty, El Mayle, Elaine Barr, Elisabeth Starr, Elissa Magsamen, Eliza Rose, Elizabeth A. Williams, Elizabeth Baldwin, Elizabeth George, Elizabeth Hobson, Elizabeth J Maxim, Elizabeth Nelson, Elizabeth Schmidt, Elizabeth Theobald, Elizabeth Traver, Elizabeth Tullett,

Ellen Hemann, Ellen M Bloom, Ellen Rine, Ellen Zemlin, Elliot Tillis, Ellis Duspiva, Emi Carr, Emilee Asbury, Emily Abernathy, Emily Bradway, Emily Brewer, Emily Clark, Emily Levy, Emily Phillips, Emily Zarse, Emma Lloyd, Emma Shaw, Eric Baldo, Eric Lamond, Eric Oehler, Eric Simonds, Eric Strother, Erica Biven, Erich Nolan, Erika Bartlett, Erika Sueker, Erin E. Jordan, Erin Elam, Erin Hardy, Erin Liparota, Erin Nieten, Erin Rich, Erin Sauer, Erin Wall, Ervin Fussner, Ethan Call, Eva Hauser, Eva Moylan, Evan Dalton, Evan Miller, Eve Cusack, Evelyn Kirkwood, Evyette Tremblay, Ezra Engels, Faith Kim, Faye Holland, Florence Caplow, Florence Frazier, Franchesca Nestor, Frederick Bloom, Gabriel Chlebowski, Gail G Hardy, Garth Riley, Garth Sabo, Gary Langell, Gary Oldham, Gayle Chalk, George Adams, George Emerson, Gerald Dunmire, Gerald Smith, Gerard Skibinski, Gillian Martin, Gillian Michael, Gina Jannazzo, Gina Lincoln, Gina Maduri, Glen Marques, Grace Kempf, Grace Landers, Grace Olsen, Gray Carlin, Greg Buehler, Greg Mankin, Greg Overtoom, Greg Rouse, Gregory, Gregory Villano, Guillaume Dury, Guillaume Salze, Hadley Biggs, Hal Grant, Haley Clements, Haley Crim, Haley Gmutza, Halle Kozak, Halo Wood, Hannah Branchick, Hannah Goverman, Hannah Kim, Hannah Knafl, Hannah Maingot, Hannah Marchaesi, Hannah Warrell, Happy Bird, Harmon Abrahamson, Harrison Werner, Hayden Kelner, Hayley Imel, Hazel Phillips, Heath Miller, Heather Barclay, Heather Bobich, Heather Hales, Heather Lamm, Heather Mantor, Heather Rowe Casey, Helen Dillon, Helen Livson, Helena Keller, Henri Schuette, Henry Ising, Henry Ross, Hilarie Klapman, Hillary Brass, Holly Law, Holly Simpson, Hope Rice, Hudson Kugele, Hunter Weedin, Iain Fleming, Ian Haliburton, Ingrid Moser, Iona Fleming, Irene Michaels, Isabel Orem, Isabelle Conner, Ivanka Simic Stanojevic, J Dale, J Koester, J Rob Taylor, J Wilkerson, Jack Berg, Jack Doherty, Jack Farzan, Jack Southall, Jack Wight, Jackie Bittner, Jackie Horn, Jaco Paco, Jacob Foster, Jacob Whitaker, Jacquelyn Randolet, Jacqui Bauer, Jaime Morales, Jama Crowe, James Andersonkirk, James Barnes, James Hubbard, Jamie Griswold, Jamie Primm, Jan McGowan, Jan Wassemiller, Jana Cooper, Jana Pereau, Jane Harvey, Janelle Coste-Swallow, Janet Akin, Janet Lind, Janet Loudermilk, Janet Truax, Janice Dykstra, Janice Evans, Jannalee Neff, Jasmine Beveridge, Jason Goshert, Jason Hill, Jason Messman, Jay Sparks, Jayne Berglund, Jean, Jean Iron, Jeanene Pratt, Jeannette Alberte, Jeannette Sutton, Jeff Dukes, Jeff Guide, Jeff Kovacs, Jeff Newby, Jelena Woehr, Jen Madaffer, Jen Walton, Jenna Cooper, Jennifer Ackerman, Jennifer Aguilar, Jennifer Benefiel, Jennifer Berger, Jennifer Christensen, Jennifer Evans, Jennifer Fisher, Jennifer Howell, Jennifer Kalas, Jennifer N. Phillips, Jennifer R. Guimont, Jennifer Shepherd, Jennifer Ulery, Jenny Andrews, Jenny Bauer, Jenny Habecker, Jerry Griego, Jess Laplante, Jess Miller-Camp, Jess Miskimens, Jess Schnur, Jessica DeWitt, Jessica Gorzo, Jessica McCory, Jessica Vandergraff, Jessie Armitage, Jessie Elsbury, Jill Courtney, Jill Keepes, Jill Kottlowski, Jillian Gorman, Jim Heiman, Jim Mckean, Jim Reilly, Jim Timmons, Jim Wooller, Jina Lawrence, Joan Frank, Joan Strassmann, Joann Boyer, Joanne Opp, Jodi Christian, Jody Maley, Joe Bitner, Joe Girgente, Joe Miles, Joe Schiefelbein, Joe Sheese, Joelle Halon, Joellen Lampman, Johanna Haas, John Evans, John Lustrea, John Salamone, John Tollefson, Jolene Headley, Jonathan Hines, Jonathan Shull, Joni James, Jordan Bennett, Joselyn Molinar, Joseph Attaway, Joseph Baird, Joseph Manley, Joseph Turpin, Joseph Zundell, Josephina H. Fornara, Josephine, Josephine R. Giannini, Josh Lodolo, Joshua Gild, Joshua Wilson, Josiah Gross, Josie Labath, Joyce Kroger, Judy Ashley, Judy Gilsdorf, Juli Jacobson, Julia Dickinson, Julia Vrtilek, Julia York, Juliana Looman, Julianna DiOrio, Julianna Escudero, Julie Bruner, Julie Roberts, Justin Barnette, Justin Havird, K Parker, K. Stewart, Kaci Lacy, Kady Carmack, Kaeko Liff, Kailey Rhein, Kaleigh Foster, Kara Andres, Kara Smith, Karee Buffin, Karen Breunig Kimbrel, Karen Hrycyna, Karen McDiarmid, Karen Scout, Karen Stancombe, Karen Stewart, Karen Sweeny, Karen Teach, Kari Brock, Kari M Campbell, Karla Simpson, Karyn Jensen, Karyn Vaughn, Kat R, Kate Hunt, Kate May, Kate Millar, Kate Squires, Kate Tallman, Katelyn Redelman, Katherine Bull, Katherine Devich, Katherine Fisher, Katherine Johnson, Katherine Pope, Katherine Sliter, Katheryn Howard, Kathleen Roos, Kathryn Amick, Kathryn Crane Doran, Kathryn Finney, Kathy Haddix, Kathy Headley, Kathy McClain, Kathy Vassari, Katie Arndt, Katie Edmonds, Katie Nolan, Katie Reichard, Kay Gross, Kay Litzer, Kay Zagst, Kayce Reed-Buechlein, Kayden Gregory, Kaylle Palma, Kc Douglass, Keath Rhymet, Keegan Deck, Keegan Stansberry, Kelley Scholfield, Kelley Wight, Kellie McBride, Kelly Barnes, Kelly Klingler, Kelly Ronald, Kelly Trimbla, Ken Baldauff, Ken Ostermiller, Kenaso Davis, Kendall Cochrane, Kenna Woods, Kenneth Cox, Kenneth Dunn, Kenneth Gisi, Kevin Cliburn, Kevin Makice, Kevin Young, Kiara Jeffries,

Kim Burrows, Kim Hughes, Kim Lascola, Kim McCain, Kim Polk, Kim Rabon, Kim Townsend, Kimberly Batker, Kimberly Guard, Kimberly Williams, Kloe Timmons, Kodi Marks, Kris Weidman, Krista Koeller, Kristen Desautell Henderson, Kristen Kobold, Kristi J. Brown, Kristina Knapp, Kristine Goossen, Kristoff Family, Kristy Harden, Kristyn DiGiovanni, Kyla Cox Deckard, Kyle Rush, Kyndra Snyder, L Blankenburg, L E Dodds, Lacey Causseaux, Larry Allen, Lashawntae Robinson, Latjor Wal, Laura Brown, Laura Kovacs, Laura Raper, Laura Reed, Laura Rojas, Laura Sargent, Laura Sheehan, Laura Useche, Laura Van Scoyoc, Lauren Bailey, Lauren Brunner, Lauren Hendrickson, Lauren Johnson, Lauren Manges, Lauren McDonald, Lauren McKinney, Lauren Stein, Lauren Witterick, Lauren Wolford, Laurie DeMott, Laurie Doss, Laurie Lee, Laurie Smith, Leah Mullins, Leane Hieber, Leanna Baldridge, Leanne Crowder, Leigh Ann Peercy, Leonard Burton, Lesa Nelson, Levi Dye, Liam Newlin, Liana Krissoff, Lidia Krylova, Lily Rieman, Linda Franklin, Linda Kellar, Linda Krystowski, Linda Penrod, Lindsay Warren, Lindsey Levine, Lindsey Potter, Lisa Champion, Lisa Derbyshire, Lisa Ellis, Lisa Knudsen, Lisa Moscoso, Lisa Shrestha, Liz Bennett, Liz Conway, Liz Finley, Liza Loza, Liza Pasch, Lizzie Shelton, Logan Rosas, Lois, Lori Lawson, Lori Maloy, Lori Schmidt, LouAnn Allen, Louise Ivers, Loyce Wilhelm, Lucas Beaver, Luci McKean, Ludmila Kaletin, Lydia Nixon, Lylanne Musselman, Lynette Hobbs, Lynleigh Wood, Lynn Antisdel, Lynn Braband, Lynn Heiman, M Wolejko, Mackenzie Maynard, Madeline Griswold, Madeline McQuiston, Madi Parker, Madison Steffel, Mads Moore, Malaak Alqaisi, Maleeha Rizvi, Mallory Pierani, Mandy Torto, Manika Chaulagain, Marci Young, Marcia Merithew, Maren Vitousek, Margaret Adkins, Margaret Kuhn, Margaret Sears, Margaret Wyant, Maria Tori, Marianne Magner, Marilyn Hollander, Marilyn Smith, Mark Guard, Mark Hodge, Mark Peck, Marketa Anderson, Markus Dickinson, Marlie Frye, Marsha Mills, Marta Barnes, Marta Diaz, Marta Jimenez, Martha Carlson Mazur, Martin Roncetti, Mary Baird, Mary Colgan, Mary E. K. Cate, Mary Hardin, Mary Hoover, Mary Jane Turnmire, Mary K, Mary Roberson, Mary Thayer, Mary Truglia, Mary Woodruff, Matt Brauer, Matt Caldie, Matt Campbell, Matt Sowders, Matthew B. White, Matthew Citron, Matthew Gerlaxh, Matthew Kelley, Matthew Kuehn, Matthew McKenna, Maya & Lucie Manners, Maya Prabhakar, Meagan Need, Meelyn Pandit, Meg Bloom, Meg McDonald, Megan Carlson, Megan Doughman, Megan Gaughan, Megan Kortlandt, Megan Ratts, Megan Thielges, Megan Van Wert, Mel McHaley, Melanie Carter, Melissa Carmichael, Melissa Fulton, Melissa Gunter, Melissa Jordan, Melissa Kocias, Melissa Meadows, Meredith Swiatek, Mia Edwards, Mia Lattanzi, Mic Bosche, Mica Gosnell, Michael Autin, Michael Burdsall, Michael Cooper, Michael Davis, Michael Gilmore, Michael Iafrato, Michael Mecham, Michael Niemack, Michael Santoro, Michael Smith, Michaela Kelly, Micheal Leisure, Michele Noel, Michelle Barnette, Michelle Del Rio, Michelle Demaree, Michelle Henkle, Michelle Roth, Michelle Spall, Miguel Guerrero, Mike Morgan, Mila Sonkin, Miles Newman, Milla Betley, Minka Wegner, Miriam Eck, Molene Greenwood, Molly Cable, Molly Deckard, Molly Fisher, Molly Guedel, Mona Mehas, Mona Miller Clayton, Monica Grubbs, Monica Jackson, Morgan Leever, Morgan Shaw, Moriah Sowders, Mylene Flores, Myra McAllister, Myra Raake, N Buis, Naman Pandey, Nancy Eppelheimer, Nancy Ferguson, Nancy Harmon, Nancy Reed, Nancy Strzelecki, Nancy Wermuth, Naomi Hood, Nat Evans, Natalie Auberry, Natalie Fiur, Natalie George, Nate Southwick, Nathan Douty, Nathan Klassen, Nathan Murnaghan, Nathaniel Clifford, Neala Lane, Nelda Nikirk, Nicholas Buehler, Nicholas E. Tishlet, Nicholas Wallace, Nick Hobbie, Nick Vetter, Nicole Dearman, Nicole Gerlach, Norm Goings, Novella Kalbaugh, Oberon Wonch, Oliver Huh, Olivia Gamsky, Olivia McDermott-Sipe, Olivia Mun, Olivia Shepherd, Omoarukhe Iruoje, Oscar Baez Montes, Oscar Morales, P Rock, Pam Bender, Pam Grtz, Pamela Rosales, Pamela Rude, Pammy Harper, Pat, Patricia Elliott, Patricia Rettig, Patricia Rose, Patricia Tredwell, Patrick Coy, Patrick Horton, Patty Franklin, Patty Wright, Paul Addington, Paul Kellogg, Paul Miller, Paul Schley, Paul Toth, Paula Goodwin, Peggy Boehnlein, Peggy Duffield, Peggy Kowalski, Peggy Moore, Peter Heile, Peter Sniffen, Petra Bragt, Phil Vickers, Phoebe Myers, Pierre Dionne, Piper & Campbell Downs, Pratush Brahma, Qnn Stork, Rachel Gugel, Rachel Harris, Rachel Kilbourne, Rachel Yoder, Rachel Zarghami, Raina McManus, Randal Pope, Randall Banks, Randy Thorburn, Ransford Walker, Rebecca Migl, Rebecca Nunley, Rebecca Winton, Rebekah Newman, Regan Hoover, Regis Terney, Rehabot Beraki, Ren Geist, Renee Albrecht, Renee Beacom, Renee Ullinskey, Rhonda Merrick, Richard Burtt, Richard MacCready, Richard Routten, Rick Dietz, Rick Liao, Rick Trevino, Rick Whitten, Riley Morris, Rita Stephens, Robby Kuehnlein, Robert Alderson, Robert Curry,

Robert Eddy, Robert Maddox, Robert Matt, Robert Rice, Robert Roos, Robert Willis, Robin Schweikart, Roger Burks, Roland-William McGurr, Ronald Kemper, Ronan Krissoff, Rosann Wattonville, Rose Taulia, Rosemary Head, Rosemary Stefani, Rowena Villarias, Roxanne Lorati, Rudy Monterrosa, Ruth Klein, Ryan Drames, Ryan Ewers, Ryan Glander, Ryan James, Ryan Koch, Ryan McKee, Ryan Skaar, Ryleigh Jones, Sage Madden, Sally Burt, Sally Ronald, Sam Cameron, Sam Cohen, Sam Hauptstueck, Sam Kiser, Sam Ritchie, Sam Tobin-Hochstadt, Sam Van Horne, Samantha Engstrom, Samantha Johnson-Helms, Samantha King, Samantha L. Heiman, Sandra Leys, Sandra Merritt, Sandy H, Sandy Ott, Sandy Youngstrom, Sara Brewer, Sara Lipshutz, Sara Neitzel, Sara Pearce, Sara Sperber, Sara Wyant, Sarah Barton, Sarah Claycomb, Sarah Cooper, Sarah Dowling, Sarah Endris, Sarah Ferritto, Sarah Keller, Sarah Lewandowski, Sarah Love, Sarah Lustrea, Sarah Marvell, Sarah Meyer, Sarah Pierson, Sarah Rhein, Sarah Sharp, Sarah Speiser, Sarah Tague, Sarah Weber, Sasha Totah, Savannah Batz, Sawyer Fouch, Schyler Marolf, Scott Hallberg, Scott Pletcher, Sean Bates, Sean Helbing, Sean Mahoney, Sean McCulloch, Seleena Baig, Serena Saltzman, Sh Kemp, Shae Kline, Shalom Drummond, Shane Lyons, Shannon, Shannon Bradley, Shannon Schroder, Sharon Barbick, Sharon Ebert, Sharon Monigold, Shawn Garrett, Shawn MacDonnell, Shea Molloy, Sheila Patton, Shelby GP, Shelby Phipps, Shelia Thomas, Shellee Ruby, Shellie Harshberger, Shelly Simmons, Shelly Swain, Sherrie Pangborn, Sherry Jordan, Sheryl MacDonald, Shirley Winslow, Shu Lai, Shulana Kpabar, Siddhartha Yaddanapudi, Solomon Nolan, Sophia Percival, Sophie Jarboe, Spencer Pevsner, Stacey Summitt-Mann, Stacie Mander, Stefanie Laputz, Stepfanie Aguillon, Stephanie Beaty, Stephanie Haskett, Stephanie McGuire, Stephanie Schene, Stephanie Temple, Stephen Ferguson, Stephen Mahoney, Stephen Sanders, Steve Lorenz, Steve Rissing, Sue Oumedian, Sunny Stachera, Sunny Thomas, Susan Atwell, Susan Grade, Susan Haislip Daleke, Susan Hayenga, Susan Helton-Groce, Susan Kramer-Gilbert, Susan Moitoza, Susan Reed, Susan S. Rosvall, Susie Shelton, Suzanne Atkinson, Suzanne Bahls, Suzanne Bergsma, Suzanne Godby Ingalsbe, Suzanne Hemann, Suzanne Thomson, Sydney Peterson, Takeo Ratnayake, Talha Ahmed, Tamara Hermesch, Tamra Thomas, Tanner Elliott, Tanya Weising-Pike, Tara Miller, Tara Phillips, Tarra Turnmire, Taylor Eagan, Tecla Winfrey, Telly & Dea Lotven, Teresa Dyer, Teresa Fancher, Teresa Morehead, Terri Furry, Terri Helms, Terri Pope, Terri Whiteman, Terry Adams, Terry Herthel, Tessa Fortnum, The Faris Family, The Kristoff Family, The Snekser-Wynne Family, Thea Atwood, Theo Cambert, Theodora Urquhart, Theresa Borel, Theresa Wilkins, Thom Shawna, Thomas Snetsinger, Thomas Tao, Tiffanie Ellis, Tiffany Edwards, Tim Barnes, Tim Holzer, Tina Greenberg, Tina Miller, Tina Nguyen, Tina Phonesaithip, Tom Angelica, Tony Brusate, Tonya Wendling, Tori Frezza, Tori Nelson, Tracy Finney, Trey McCullum, Triaha Burton, Troy Cockrum, Tyeisha Fordham, Tyler Benton, Valeri Mullins, Valerie Joyal, Vanessa Speight, Veronica Bobskill, Vicki McKay, Vickie Smith, Vincent Le, Violet Phillips, Virginia Lauterbach, Virginia Stoltz, Virginia Urban, Vivek Govind Kumar, Vivian Valena Oliva, VJ Reilly, Wally Rosvall, Wayne Brown, Wendy Byers, Wendy Cornwell, Wendy King, Wendy Mattson, Whitney Nicole Harruff, Whitney Wickesberg, Wiley Weinmann, Will Boland, Will Pierce, William Bell, William MacDonald, William Montgomery, William Pastorius IV, Wyatt Perkoski, Wylah Brahaum, Wynn Webster, Yixiao Liu, Yvonne Hurd, Zachary Bishop, Zoe Ackerman, Zoe Shive, Zofia Ahmad

*SolarBird app Privacy Notice*: At Indiana University (IU), we are committed to protecting the privacy and confidentiality of personal information entrusted to us. By accessing and using IU's services, you acknowledge and consent to the practices described in our global privacy statement here: https://privacy.iu.edu/privacy/global.html.

For additional information outlining how IU College/Luddy collects, uses, and safeguards personal information obtained specifically through our website (https://luddyeclipse2024.org/solarbird.html), please also review the information below. Continued use of our website indicates consent to the collection, use, and disclosure of this information as described in this notice. Visitors to other IU websites should review the privacy notices for the sites they visit, as other units at the university may collect and use visitor information in different ways. IU College/Luddy is not responsible for the content of other websites or for the privacy practices of websites outside the scope of this notice.

Changes: Because Internet technologies continue to evolve rapidly, IU College/Luddy may make appropriate changes to this notice in the future. Any such changes will be consistent with our commitment to respecting visitor privacy and will be clearly posted in a revised privacy notice.

Passive/Automatic Collection: In addition to any information outline in the global statement, our server and/or site collects the following: Your IP address, The date and time of visit, GPS coordinates. This technical information is retained in detail for up to 3650 days. Some technical information is retained in aggregate for up to 3650 days.

Active/Manual/Voluntary Collection: In addition to the technical information about your visit described above (or cookies, described below), we may ask you to provide information voluntarily, such as through forms or other manual input. We might ask for you to provide this information in order to make products and services available to you, to maintain and manage our relationship with you, including providing associated services or to better understand and serve your needs. This information is generally retained as long as you continue to maintain a relationship with us. Your providing this information is wholly voluntary. However, not providing the requested information (or subsequently asking that the data be removed) may affect our ability to deliver the products or service for which the information is needed. Providing the requested information indicates your consent to the collection, use, and disclosure of this information as described in this notice. Information we may actively collect could include: The email addresses of those who communicate with us via email; Name; Information volunteered by the visitor, such as preferences, survey information and/or site; registrations; Observed bird behavior.

Information Used For Contact: If you supply us with your postal/mailing address: You will only receive the information for which you provided us your address.

Information Sharing: We may share aggregate, non-personally identifiable information with other entities or organizations. Except as described in the IU Privacy statement, we will not share any information with any other entities or organizations for any reason. Except as provided in the Disclosure of Information section below, we do not attempt to use the technical information discussed in this section to identify individual visitors.

Cookies: For more information on how we use cookies, please review the IU Privacy statement. Our site does not use cookies to store information about your actions or choices on pages associated with our site.

Children: This site is not directed to children under 13 years of age, does not sell products or services intended for purchase by children, and does not knowingly collect or store any personal information, even in aggregate, about children under the age of 13. We encourage parents and teachers to be involved in children's Internet explorations. It is particularly important for parents to guide their children when they are asked to provide personal information online.

Use of Third Party Services: Our website does not utilize web analytics services beyond what is noted in the IU Privacy statement.

Security: Due to the rapidly evolving nature of information technologies, no transmission of information over the Internet can be guaranteed to be completely secure. While Indiana University is committed to protecting user privacy, IU cannot guarantee the security of any information users transmit to university sites, and users do so at their own risk.

- We have appropriate security measures in place in our physical facilities to protect against the loss, misuse, or alteration of information that we have collected from you at our site.
- Once we receive user information, we will use reasonable safeguards consistent with prevailing industry standards and commensurate with the sensitivity of the data being stored to maintain the security of that information on our systems.
- We will comply with all applicable federal, state and local laws regarding the privacy and security of user information.

Links to non-university sites: Indiana University is not responsible for the availability, content, or privacy practices of non-university sites. Non-university sites are not bound by this site privacy notice policy and may or may not have their own privacy policies.

Privacy Notice Changes: From time to time, we may use visitor information for new, unanticipated uses not previously disclosed in our privacy notice. We will post the policy changes to our Website to notify you of these changes and provide you with the ability to opt out of these new uses. If you are concerned about how your information is used, you should check back at our Website periodically. Visitors may prevent their information from being used for purposes other than those for which it was originally collected by sending us email at the listed address.

Supplemental Information: Email addresses are not required information; by supplying your email address, you opt into receiving occasional emails that disseminate the results of this research study. They are securely stored and will never be shared with third parties or used for any other purposes beyond disseminating the research results. Names are not required information; by supplying your name and providing valid bird behavior observations, you opt in to be acknowledged in a forthcoming scientific publication. They are securely stored and will never be shared with third parties or used for any other purposes beyond acknowledgement of your contributions. IP addresses are securely stored for internal use in data validation and quality control; they are not shared publicly in any form, including in scientific datasets or publications. GPS coordinates and observation times are securely stored for internal use in scientific data analysis (including but not limited to correlating observations with eclipse phase, temperature, cloud coverage, barometric pressure, and other nonidentifying variables of scientific interest). GPS coordinates will never be shared with third parties or used for other purposes. Bird observations are not personally identifying and will be shared as open data and in scientific publications.

Contact Information: If you have questions or concerns about this policy, please contact us. Indiana University, ATTN: Liz Aguilar, 107 S Indiana Ave. Bloomington, IN. 47405. USA.

If you feel as though this site's privacy practices differ from the information stated, you may contact us at the listed address or phone number. If you feel that this site is not following its stated policy and communicating with the owner of this site does not resolve the matter, or if you have general questions or concerns about privacy or information technology policy at Indiana University, please contact the chief privacy officer through the University Information Policy Office, 812-855-UIPO,.

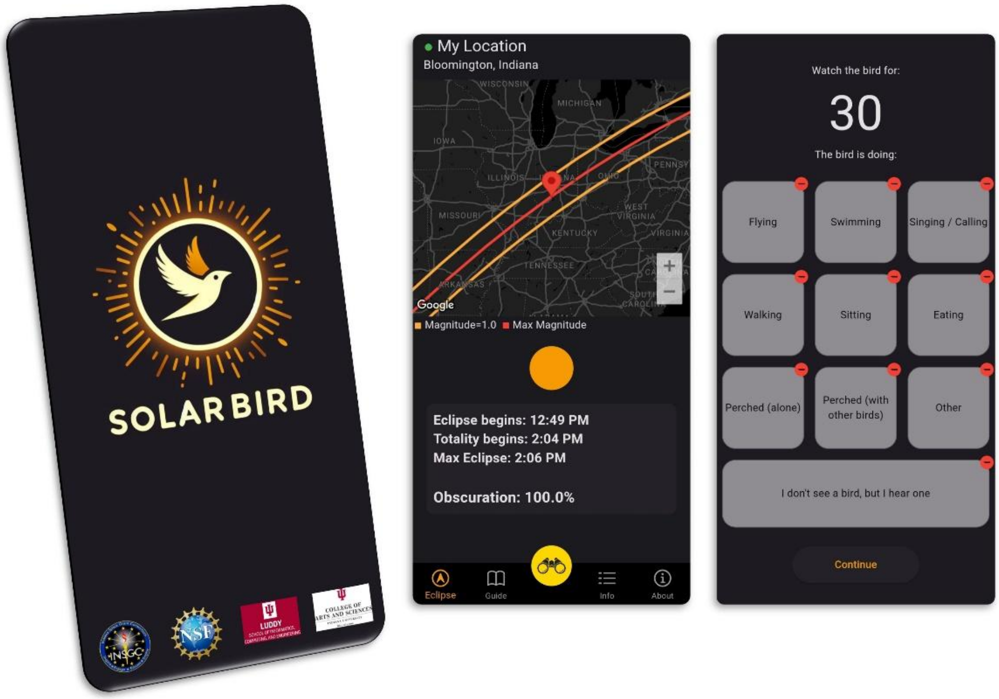


**Fig. S1. Screenshots of *SolarBird* app.** Screenshots of the *SolarBird* app (left panel), including the initial user homepage with a map and eclipse-related countdown (middle panel), and the observation portal with a 30-second countdown timer and 10 non-mutually exclusive click-boxes for users to report what they saw or heard (right panel).

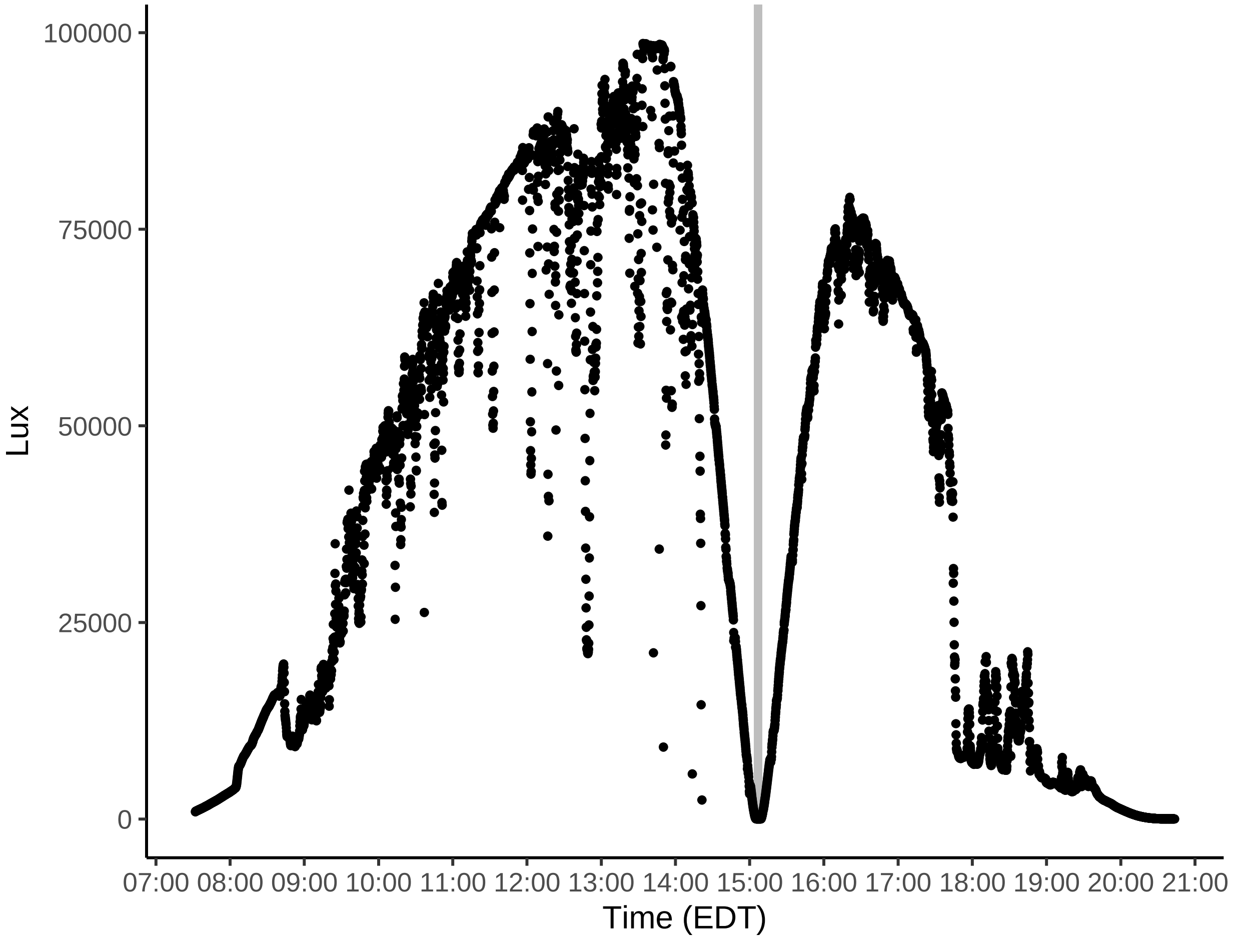


**Fig. S2. Illuminance on the day of totality.** Illuminance was recorded on the day of the eclipse, using a LICOR LI-180 spectrometer in central Bloomington, IN USA. Data are presented in units of lux (1 lux = 1 lumen $m^{-2}$). The spectrometer was placed in an open area with limited interference from surrounding foliage. The shaded bar marks the period of totality.

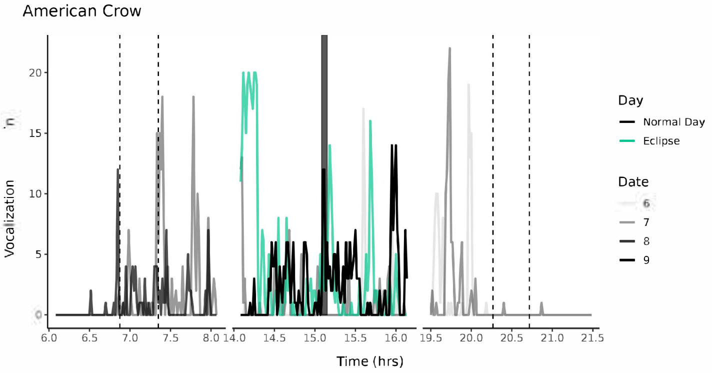
American Crow
Vocalization
Time (hrs)
Day
Normal Day
Eclipse
Date
6
7
8
9

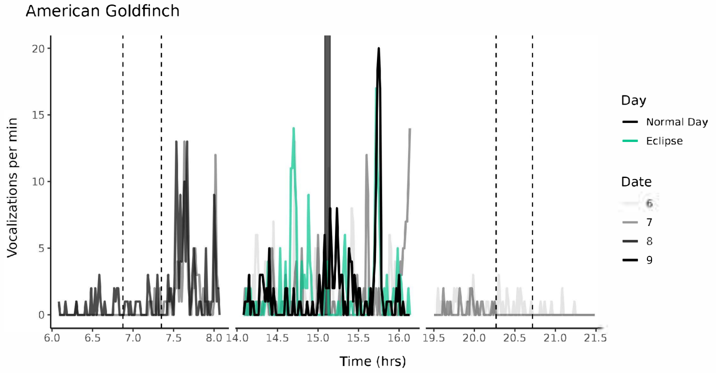
American Goldfinch
Vocalizations per min
Time (hrs)
Day
Normal Day
Eclipse
Date
6
7
8
9

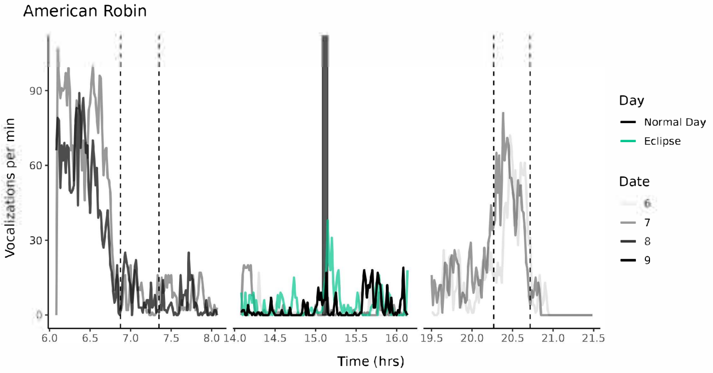
American Robin
Vocalizations per min
Time (hrs)
Day
Normal Day
Eclipse
Date
6
7
8
9

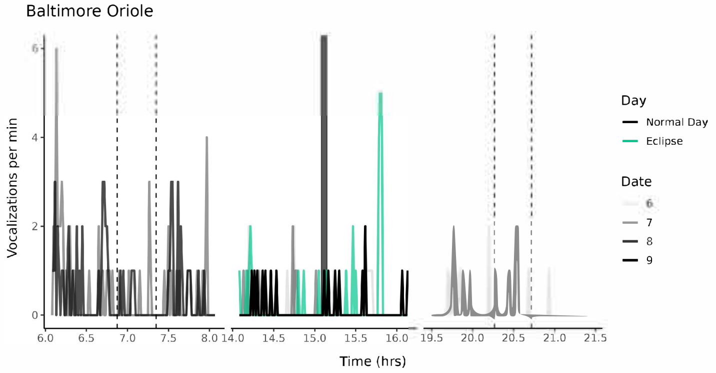
Baltimore Oriole
Vocalizations per min
Time (hrs)
Day
Normal Day
Eclipse
Date
6
7
8
9

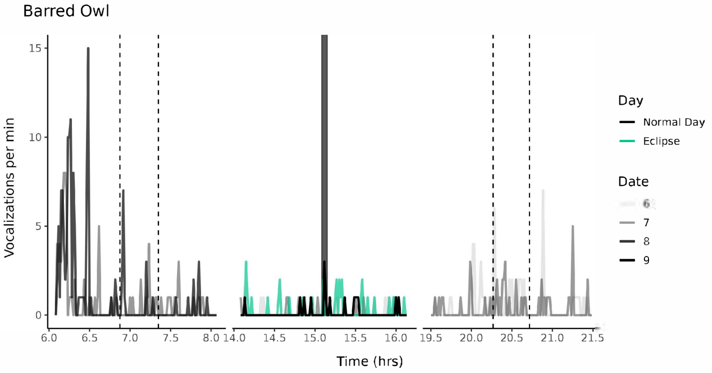
Barred Owl
Vocalizations per min
Time (hrs)
Day
Normal Day
Eclipse
Date
6
7
8
9

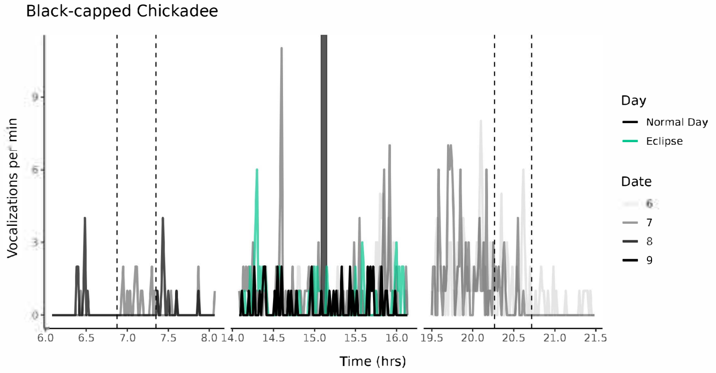
Black-capped Chickadee
Vocalizations per min
Time (hrs)
Day
Normal Day
Eclipse
Date
6
7
8
9

Blue Jay

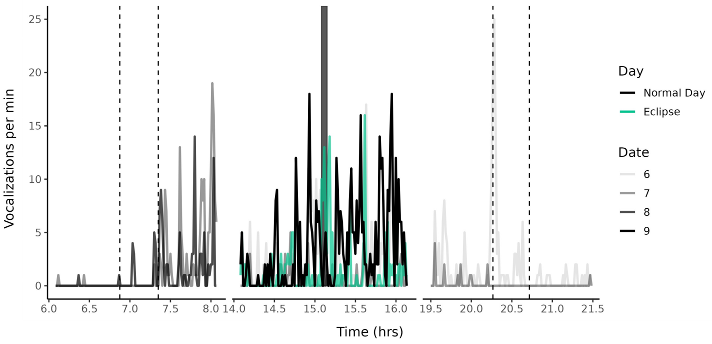


Brown Thrasher

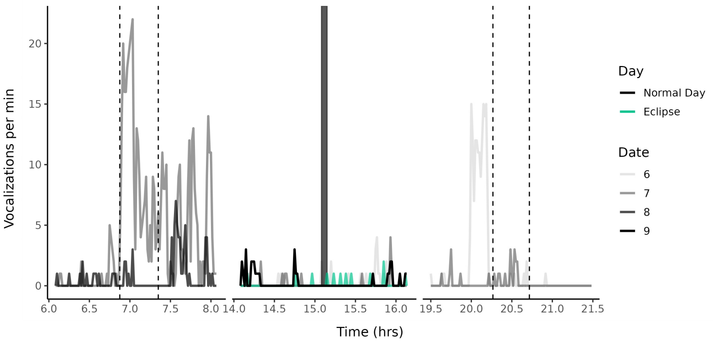


Brown-headed Cowbird

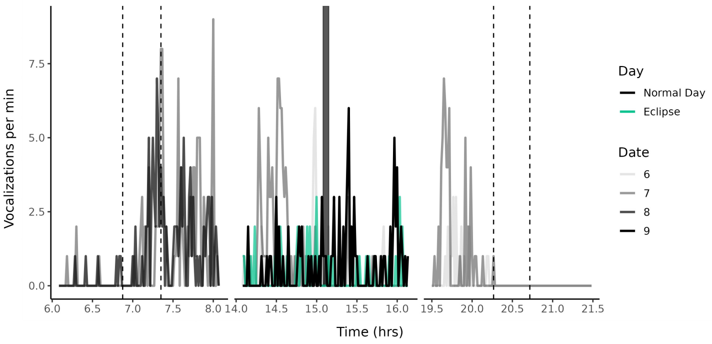


Canada Goose

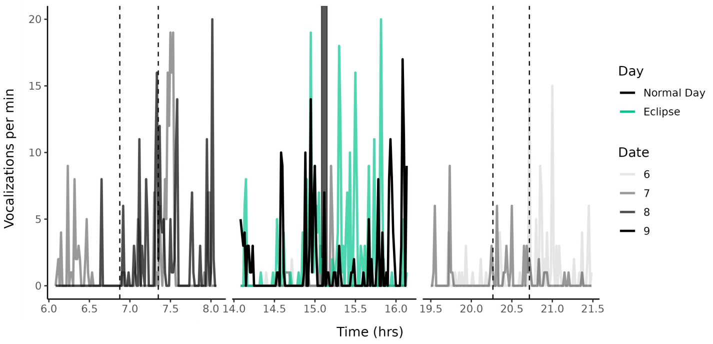


Carolina Chickadee

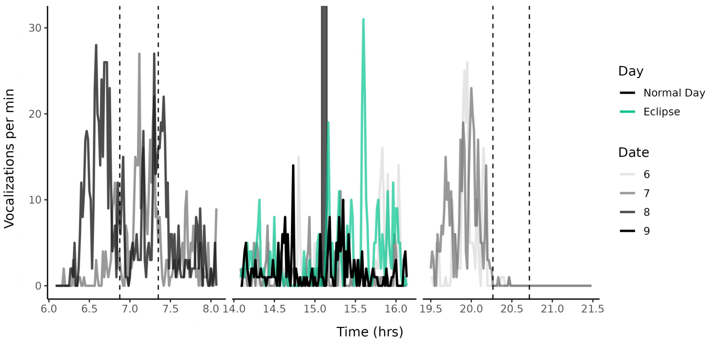


Carolina Wren

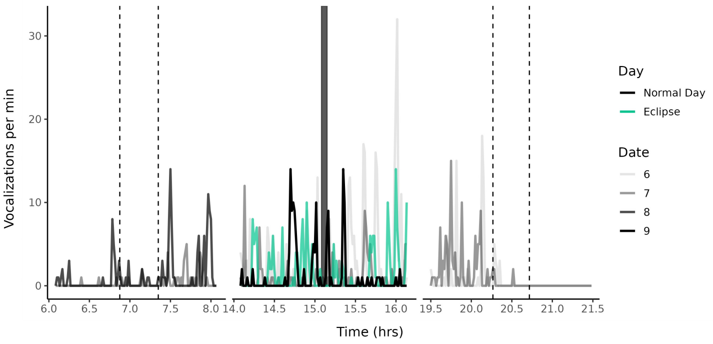

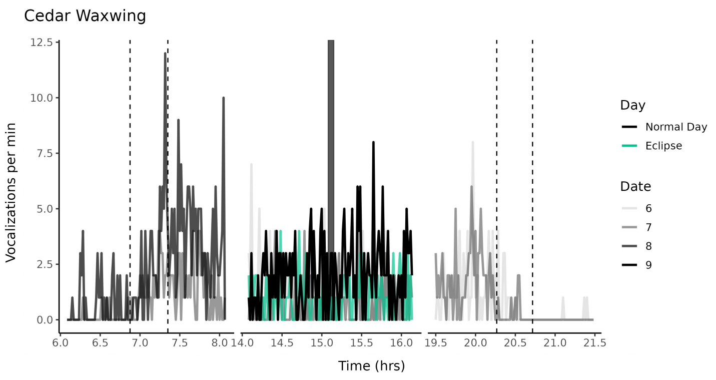
Cedar Waxwing
Vocalizations per min
Time (hrs)
Day
Normal Day
Eclipse
Date
6
7
8
9

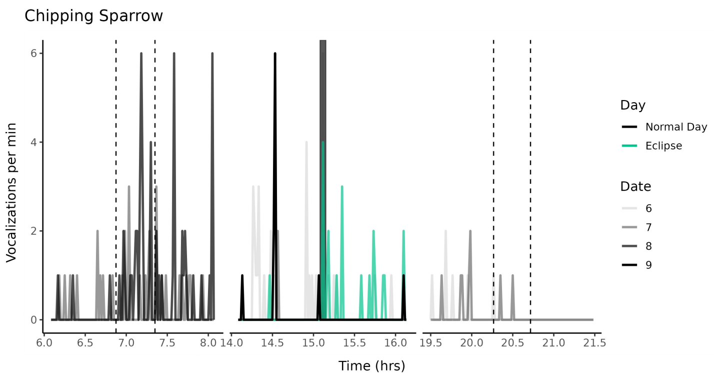
Chipping Sparrow
Vocalizations per min
Time (hrs)
Day
Normal Day
Eclipse
Date
6
7
8
9

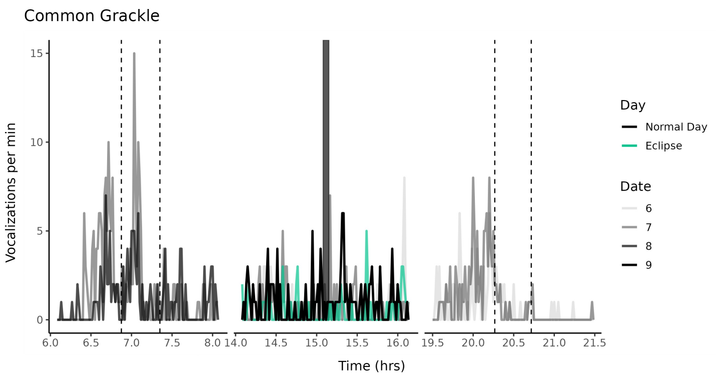
Common Grackle
Vocalizations per min
Time (hrs)
Day
Normal Day
Eclipse
Date
6
7
8
9

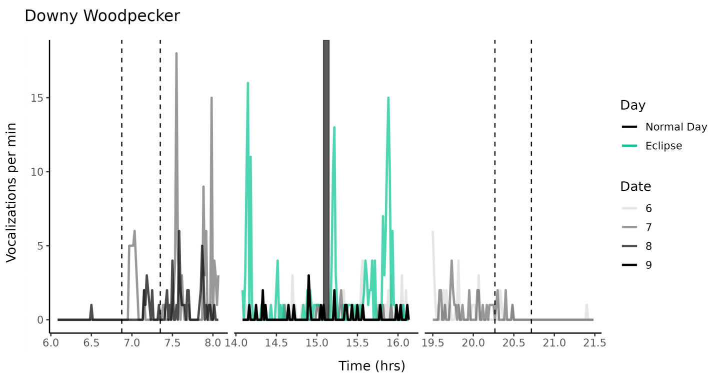
Downy Woodpecker
Vocalizations per min
Time (hrs)
Day
Normal Day
Eclipse
Date
6
7
8
9

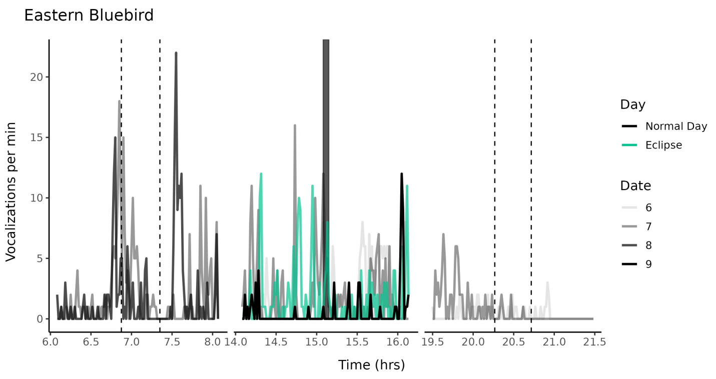
Eastern Bluebird
Vocalizations per min
Time (hrs)
Day
Normal Day
Eclipse
Date
6
7
8
9

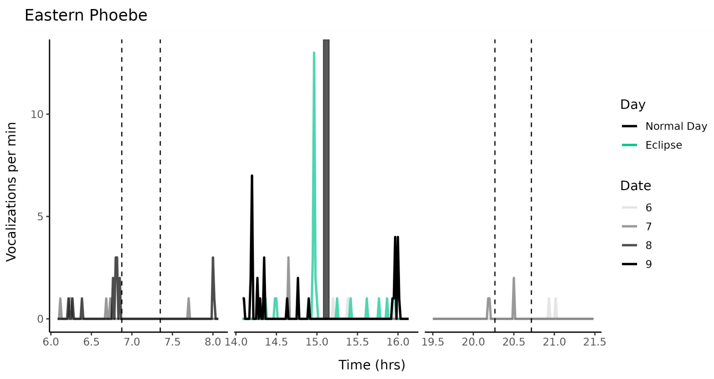
Eastern Phoebe
Vocalizations per min
Time (hrs)
Day
Normal Day
Eclipse
Date
6
7
8
9

Eastern Towhee

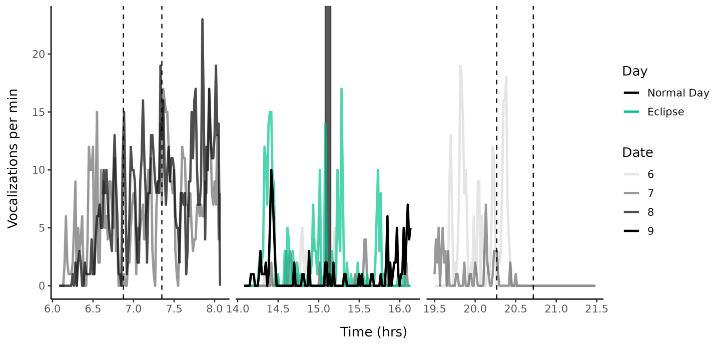


Eurasian Collared-Dove

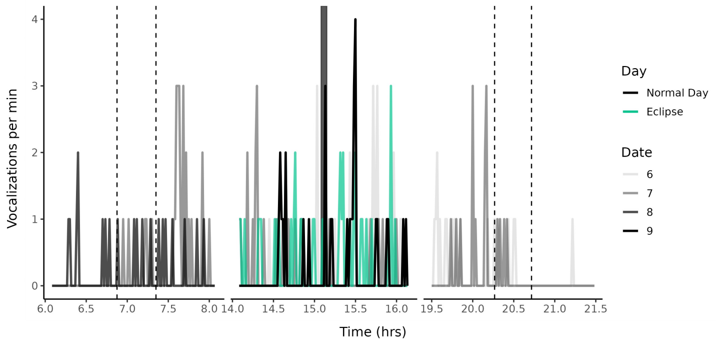


European Starling

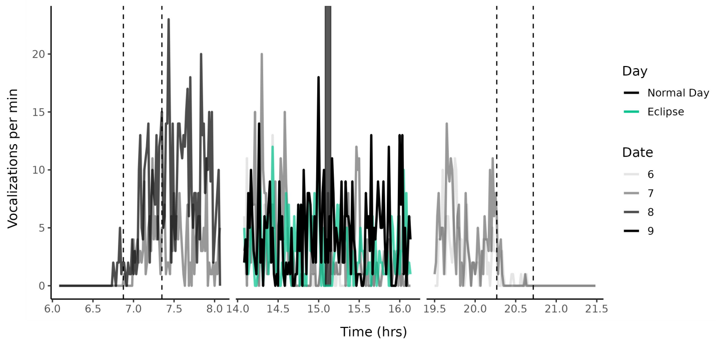


Field Sparrow

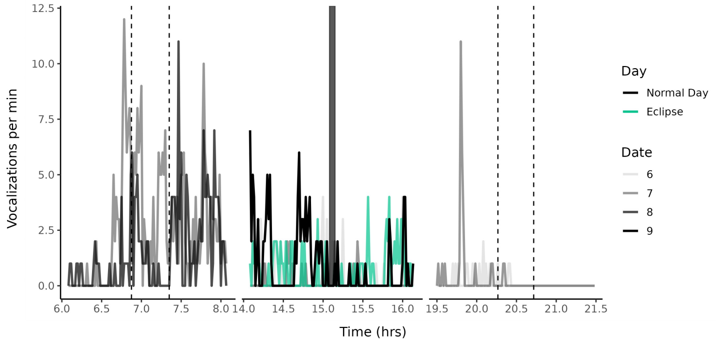


Fish Crow

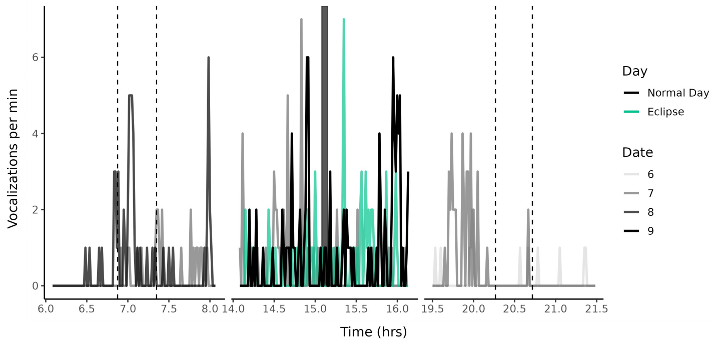


Golden-crowned Kinglet

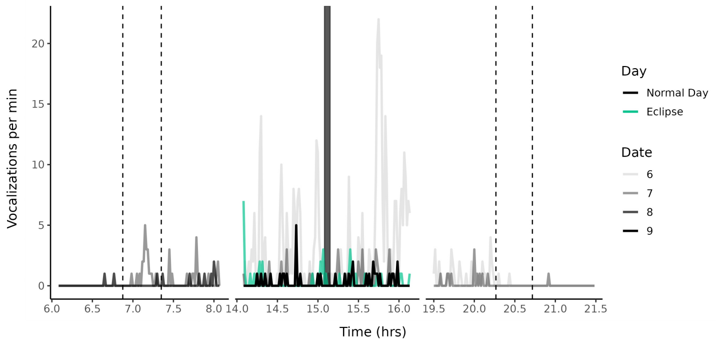

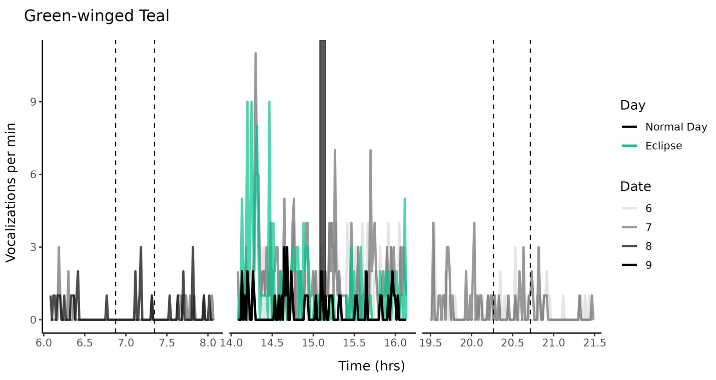
Green-winged Teal
Vocalizations per min
Time (hrs)
Day
Normal Day
Eclipse
Date
6
7
8
9

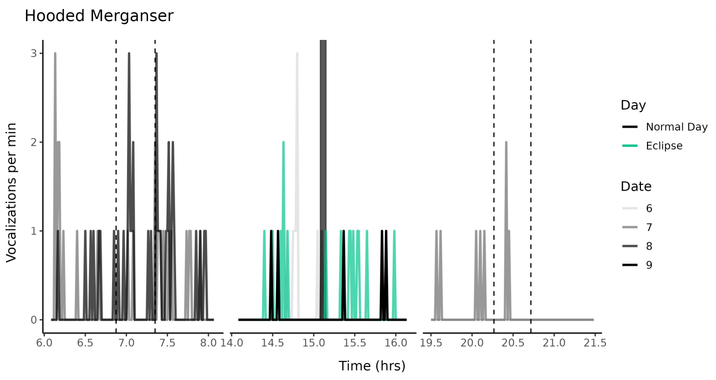
Hooded Merganser
Vocalizations per min
Time (hrs)
Day
Normal Day
Eclipse
Date
6
7
8
9

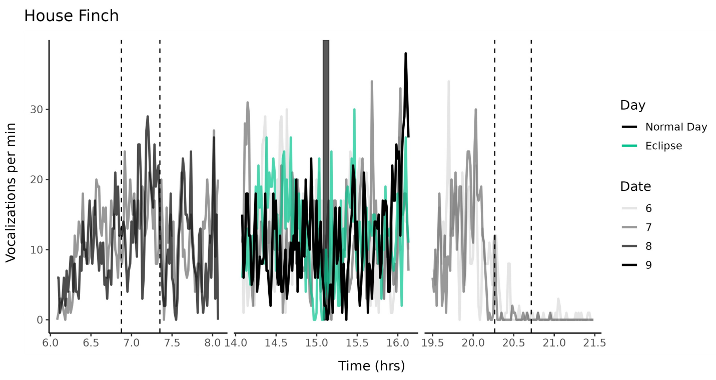
House Finch
Vocalizations per min
Time (hrs)
Day
Normal Day
Eclipse
Date
6
7
8
9

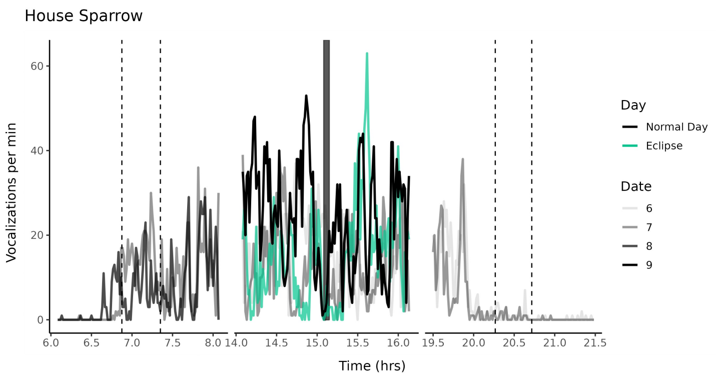
House Sparrow
Vocalizations per min
Time (hrs)
Day
Normal Day
Eclipse
Date
6
7
8
9

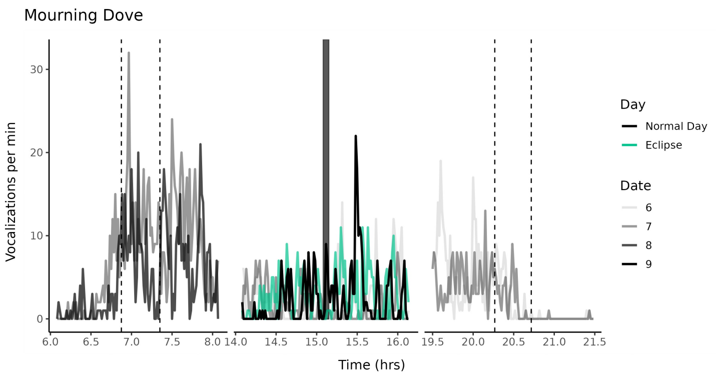
Mourning Dove
Vocalizations per min
Time (hrs)
Day
Normal Day
Eclipse
Date
6
7
8
9

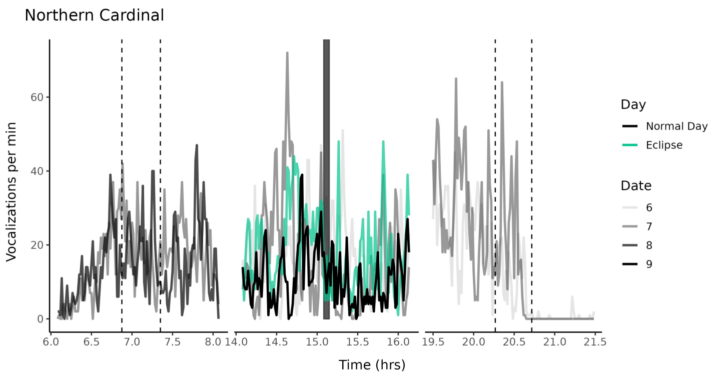
Northern Cardinal
Vocalizations per min
Time (hrs)
Day
Normal Day
Eclipse
Date
6
7
8
9

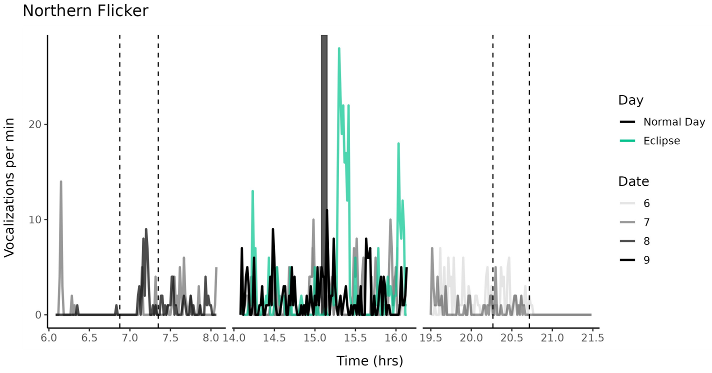
Northern Flicker
Vocalizations per min
Time (hrs)
Day
Normal Day
Eclipse
Date
6
7
8
9

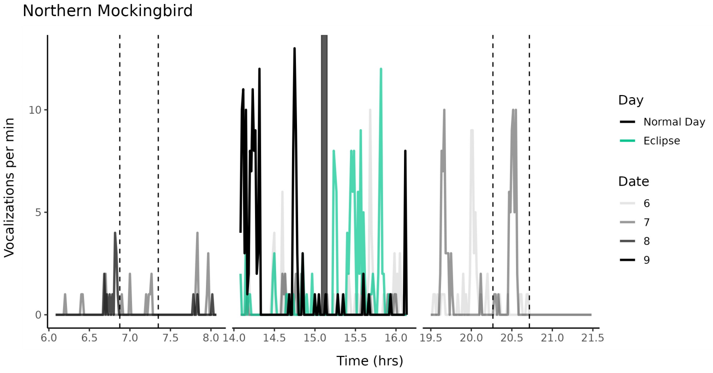
Northern Mockingbird
Vocalizations per min
Time (hrs)
Day
Normal Day
Eclipse
Date
6
7
8
9

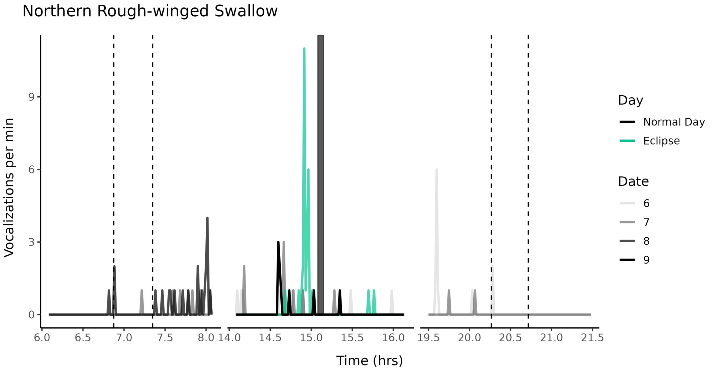
Northern Rough-winged Swallow
Vocalizations per min
Time (hrs)
Day
Normal Day
Eclipse
Date
6
7
8
9

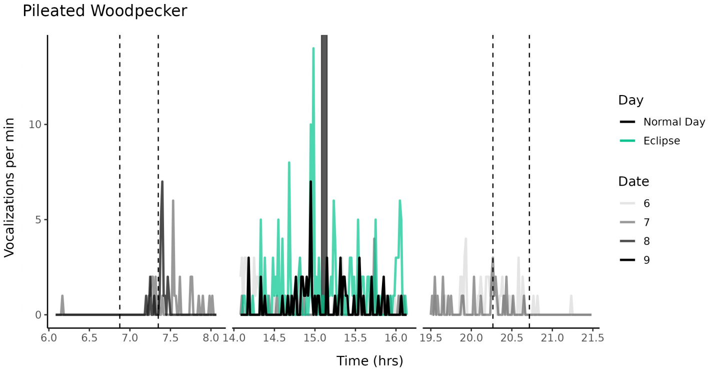
Pileated Woodpecker
Vocalizations per min
Time (hrs)
Day
Normal Day
Eclipse
Date
6
7
8
9

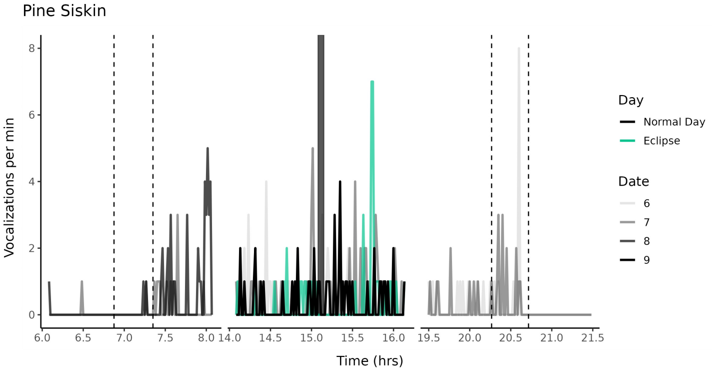
Pine Siskin
Vocalizations per min
Time (hrs)
Day
Normal Day
Eclipse
Date
6
7
8
9

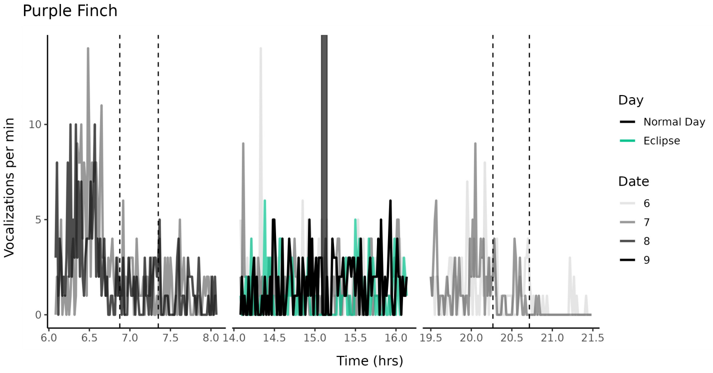
Purple Finch
Vocalizations per min
Time (hrs)
Day
Normal Day
Eclipse
Date
6
7
8
9

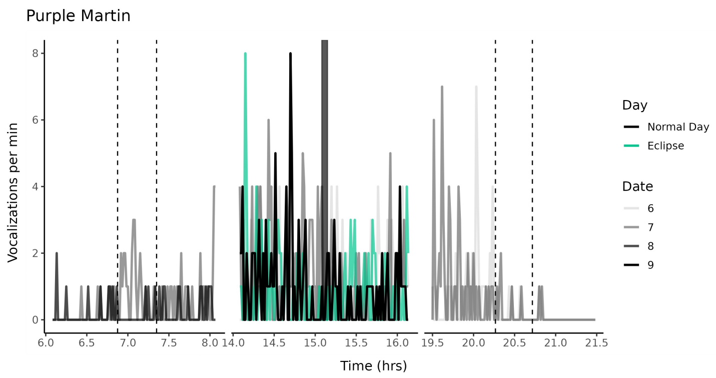
Purple Martin
Vocalizations per min
Time (hrs)
Day
Normal Day
Eclipse
Date
6
7
8
9

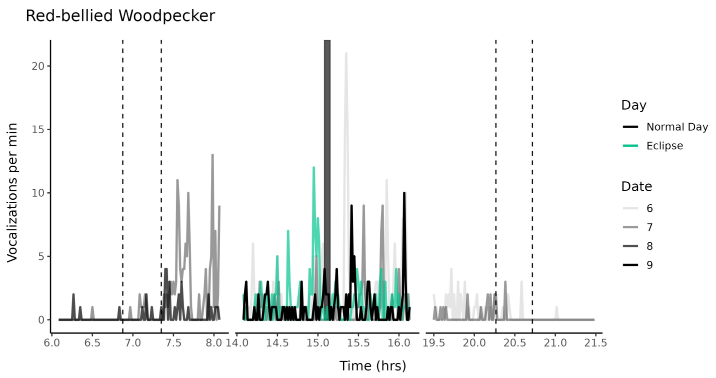
Red-bellied Woodpecker
Vocalizations per min
Time (hrs)
Day
Normal Day
Eclipse
Date
6
7
8
9

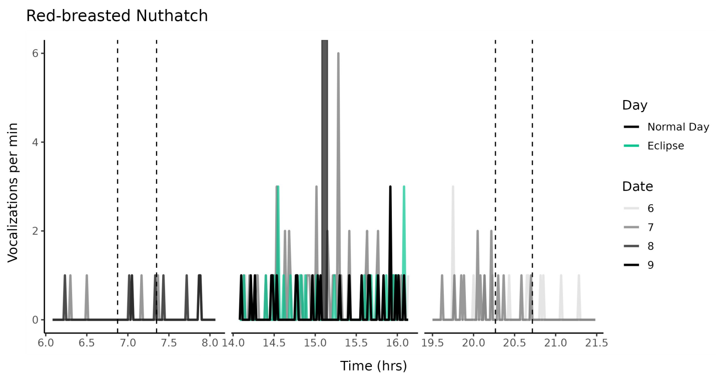
Red-breasted Nuthatch
Vocalizations per min
Time (hrs)
Day
Normal Day
Eclipse
Date
6
7
8
9

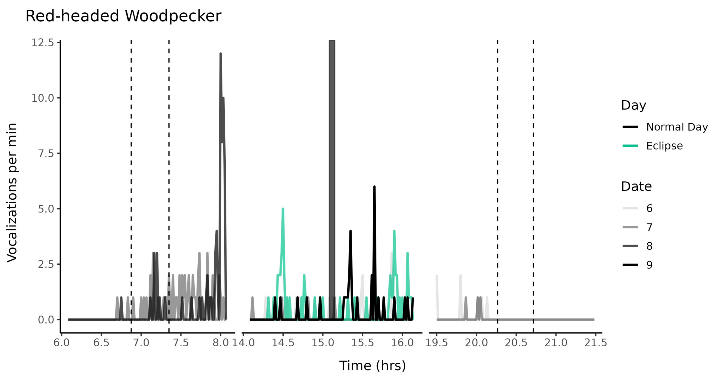
Red-headed Woodpecker
Vocalizations per min
Time (hrs)
Day
Normal Day
Eclipse
Date
6
7
8
9

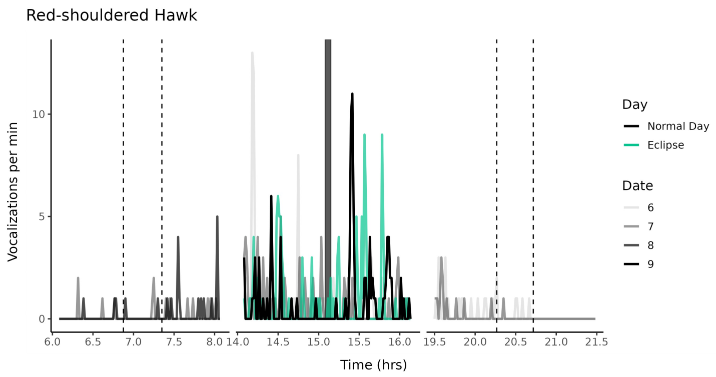
Red-shouldered Hawk
Vocalizations per min
Time (hrs)
Day
Normal Day
Eclipse
Date
6
7
8
9

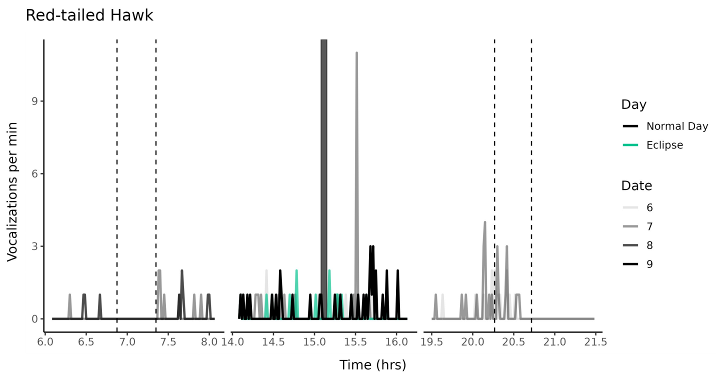
Red-tailed Hawk
Vocalizations per min
Time (hrs)
Day
Normal Day
Eclipse
Date
6
7
8
9

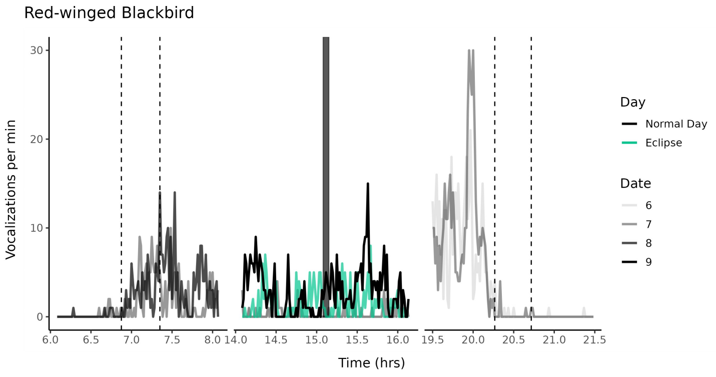
Red-winged Blackbird
Vocalizations per min
Time (hrs)
Day
Normal Day
Eclipse
Date
6
7
8
9

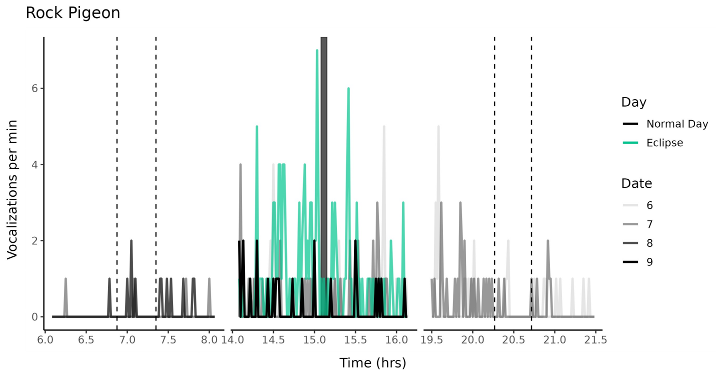
Rock Pigeon
Vocalizations per min
Time (hrs)
Day
Normal Day
Eclipse
Date
6
7
8
9

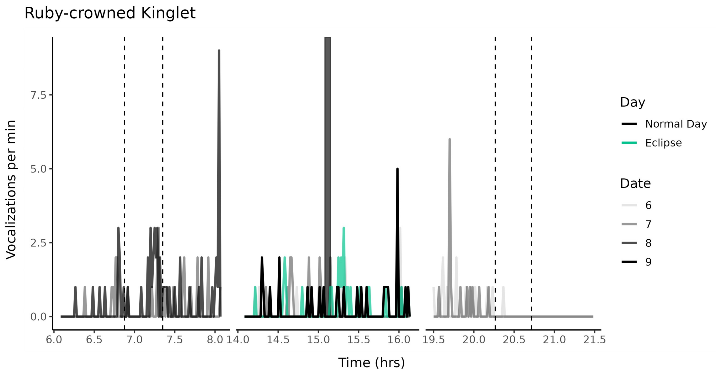
Ruby-crowned Kinglet
Vocalizations per min
Time (hrs)
Day
Normal Day
Eclipse
Date
6
7
8
9

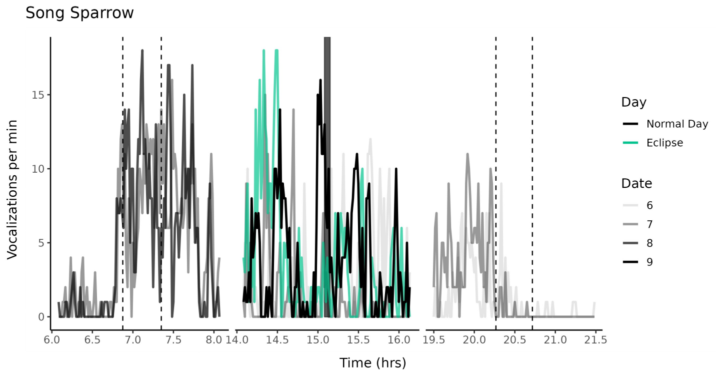
Song Sparrow
Vocalizations per min
Time (hrs)
Day
Normal Day
Eclipse
Date
6
7
8
9

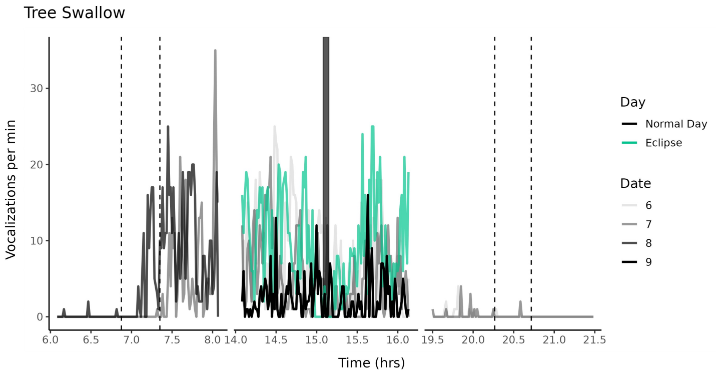
Tree Swallow
Vocalizations per min
Time (hrs)
Day
Normal Day
Eclipse
Date
6
7
8
9

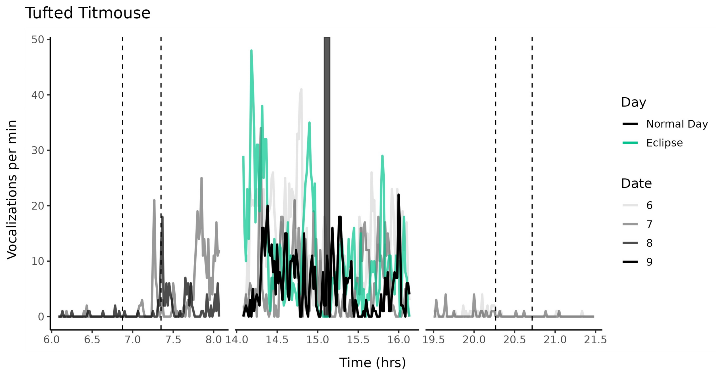
Tufted Titmouse
Vocalizations per min
Time (hrs)
Day
Normal Day
Eclipse
Date
6
7
8
9

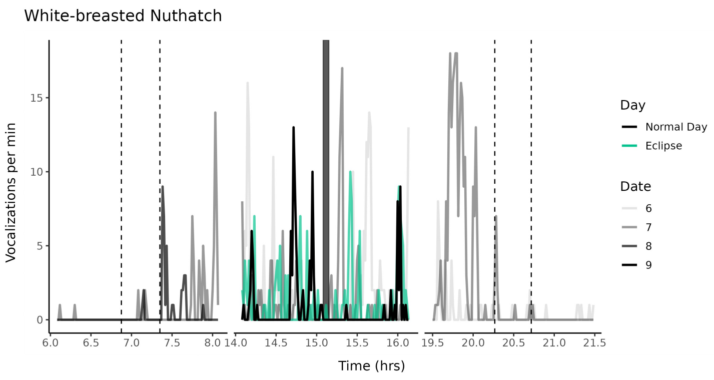
White-breasted Nuthatch
Vocalizations per min
Time (hrs)
Day
Normal Day
Eclipse
Date
6
7
8
9

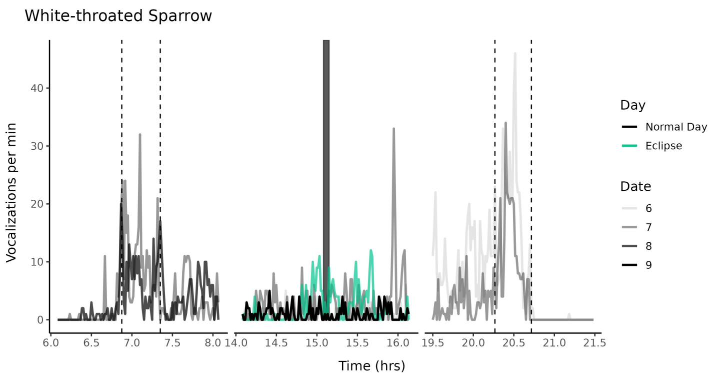
White-throated Sparrow
Vocalizations per min
Time (hrs)
Day
Normal Day
Eclipse
Date
6
7
8
9

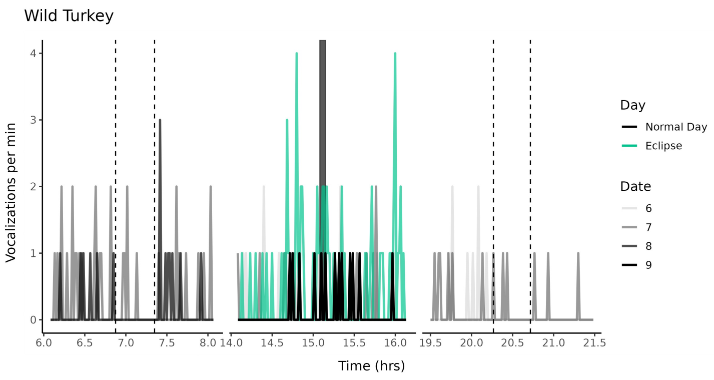
Wild Turkey
Vocalizations per min
Time (hrs)
Day
Normal Day
Eclipse
Date
6
7
8
9

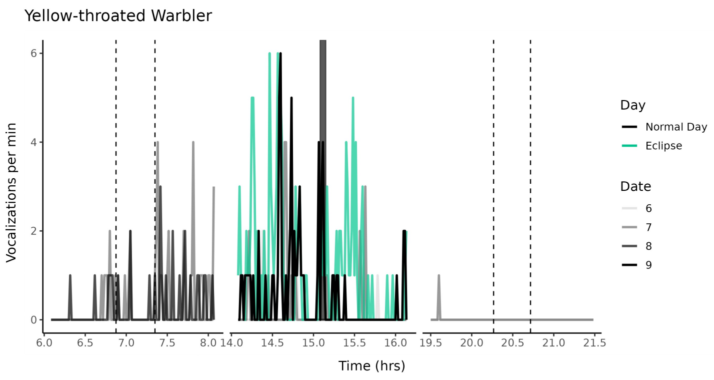
Yellow-throated Warbler
Vocalizations per min
Time (hrs)
Day
Normal Day
Eclipse
Date
6
7
8
9

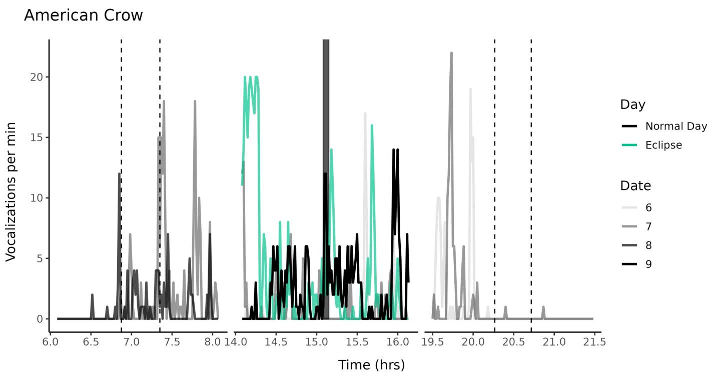
American Crow
Vocalizations per min
Time (hrs)
Day
Normal Day
Eclipse
Date
6
7
8
9

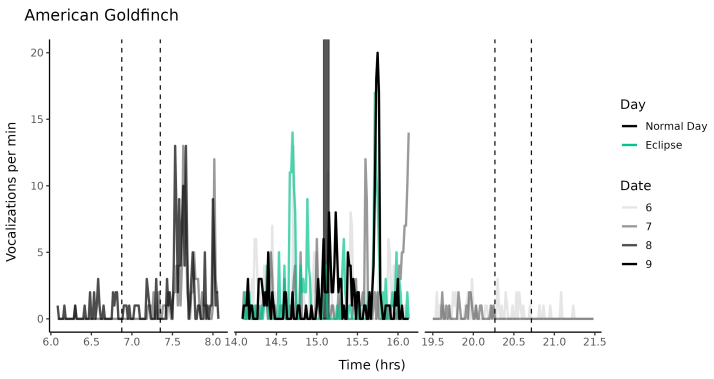
American Goldfinch
Vocalizations per min
Time (hrs)
Day
Normal Day
Eclipse
Date
6
7
8
9

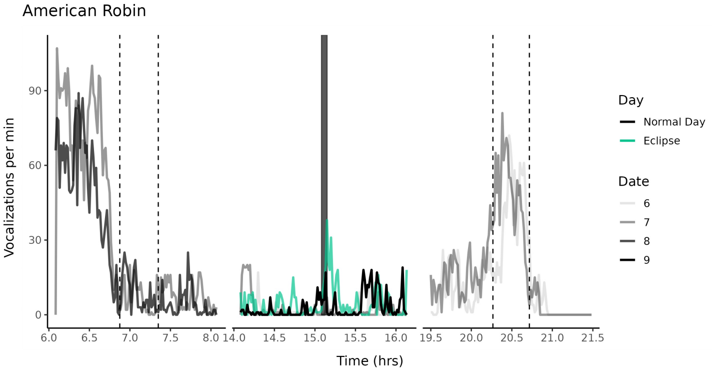
American Robin
Vocalizations per min
Time (hrs)
Day
Normal Day
Eclipse
Date
6
7
8
9

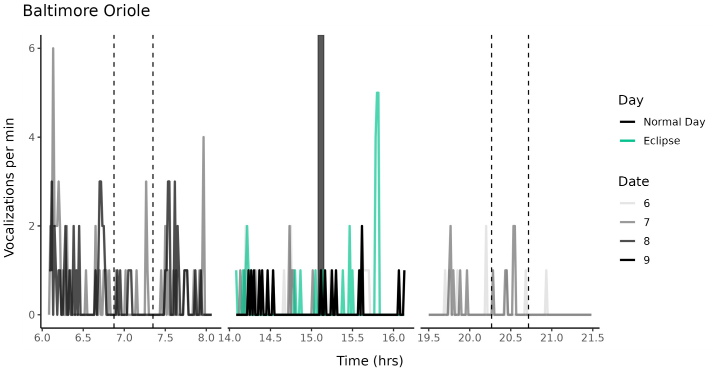
Baltimore Oriole
Vocalizations per min
Time (hrs)
Day
Normal Day
Eclipse
Date
6
7
8
9

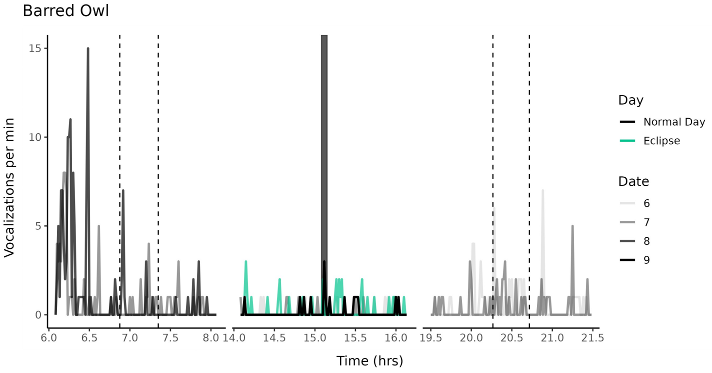
Barred Owl
Vocalizations per min
Time (hrs)
Day
Normal Day
Eclipse
Date
6
7
8
9

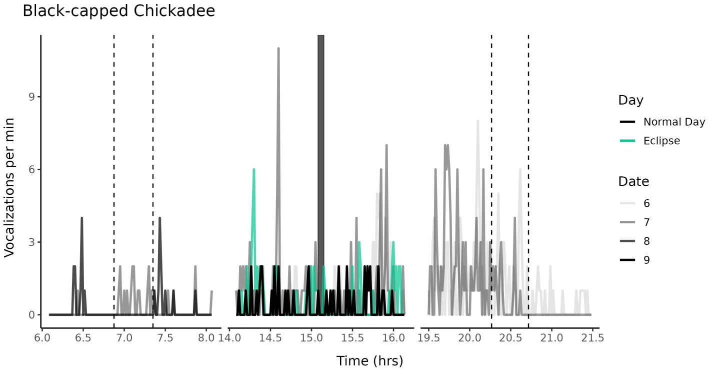
Black-capped Chickadee
Vocalizations per min
Time (hrs)
Day
Normal Day
Eclipse
Date
6
7
8
9

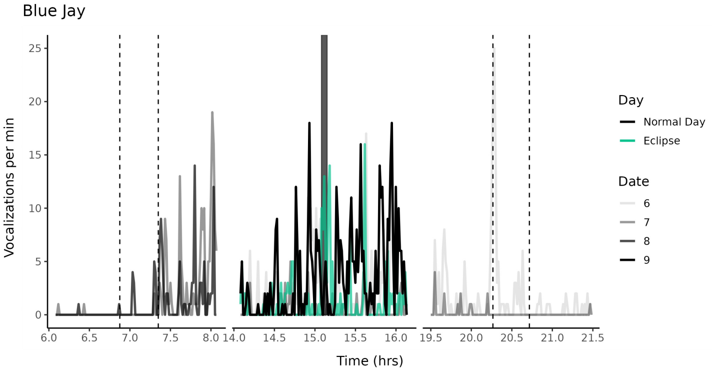
Blue Jay
Vocalizations per min
Time (hrs)
Day
Normal Day
Eclipse
Date
6
7
8
9

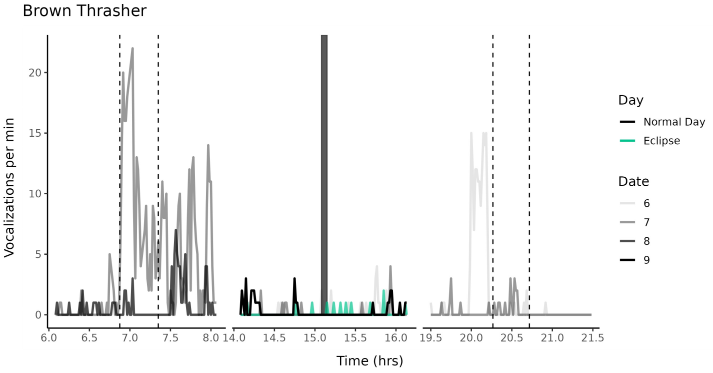
Brown Thrasher
Vocalizations per min
Time (hrs)
Day
Normal Day
Eclipse
Date
6
7
8
9

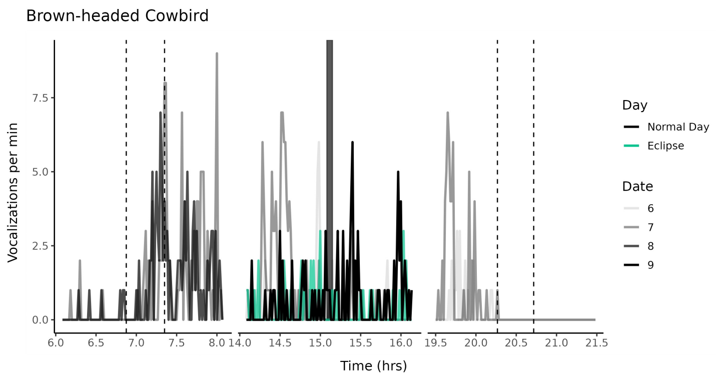
Brown-headed Cowbird
Vocalizations per min
Time (hrs)
Day
Normal Day
Eclipse
Date
6
7
8
9

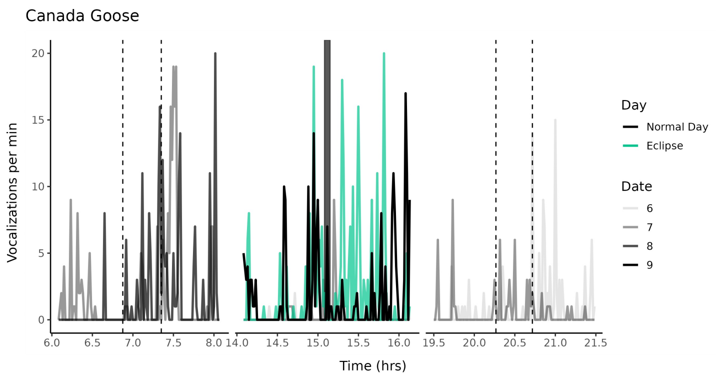
Canada Goose
Vocalizations per min
Time (hrs)
Day
Normal Day
Eclipse
Date
6
7
8
9

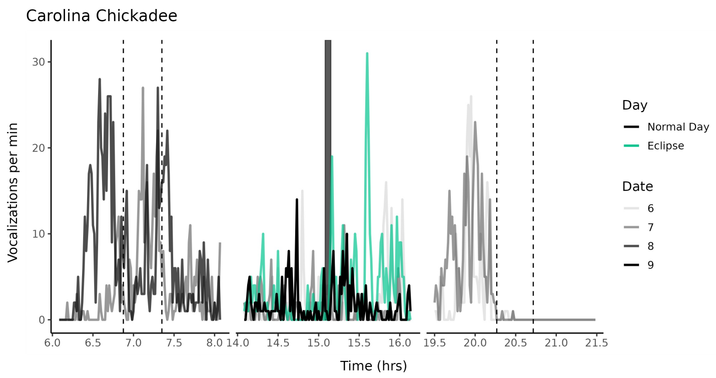
Carolina Chickadee
Vocalizations per min
Time (hrs)
Day
Normal Day
Eclipse
Date
6
7
8
9

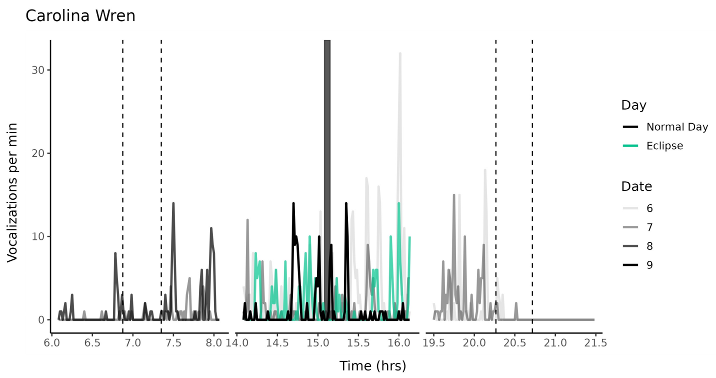
Carolina Wren
Vocalizations per min
Time (hrs)
Day
Normal Day
Eclipse
Date
6
7
8
9

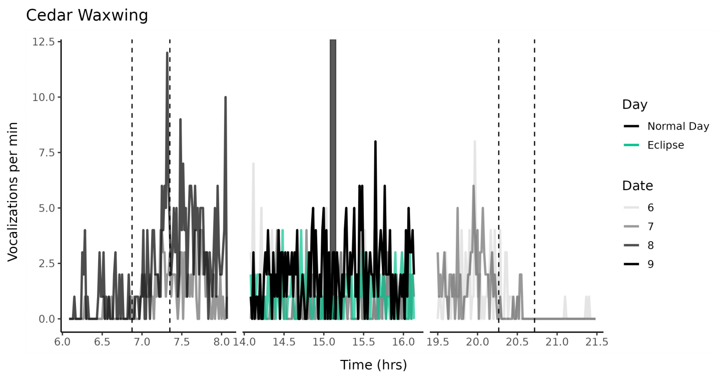
Cedar Waxwing
Vocalizations per min
Time (hrs)
Day
Normal Day
Eclipse
Date
6
7
8
9

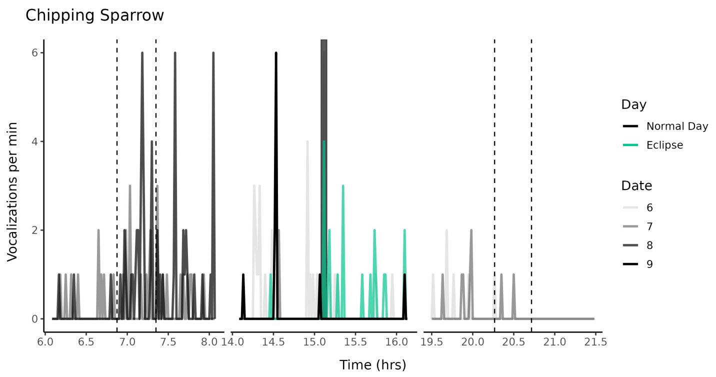
Chipping Sparrow
Vocalizations per min
Time (hrs)
Day
Normal Day
Eclipse
Date
6
7
8
9

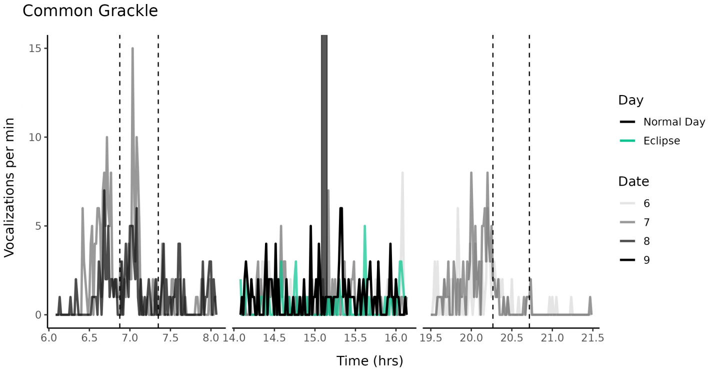
Common Grackle
Vocalizations per min
Time (hrs)
Day
Normal Day
Eclipse
Date
6
7
8
9

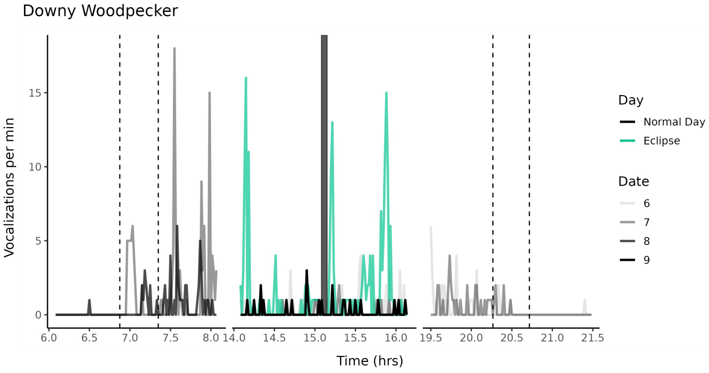
Downy Woodpecker
Vocalizations per min
Time (hrs)
Day
Normal Day
Eclipse
Date
6
7
8
9

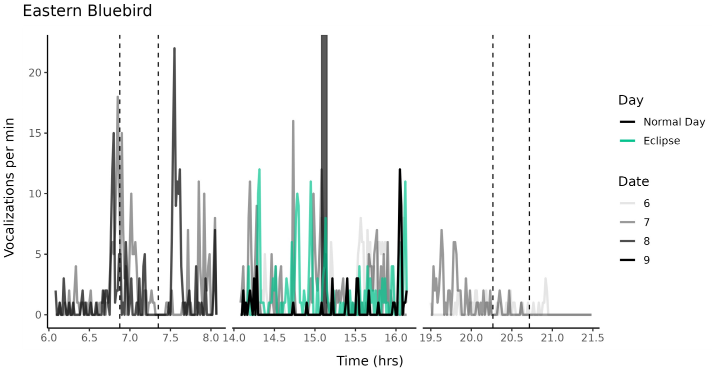
Eastern Bluebird
Vocalizations per min
Time (hrs)
Day
Normal Day
Eclipse
Date
6
7
8
9

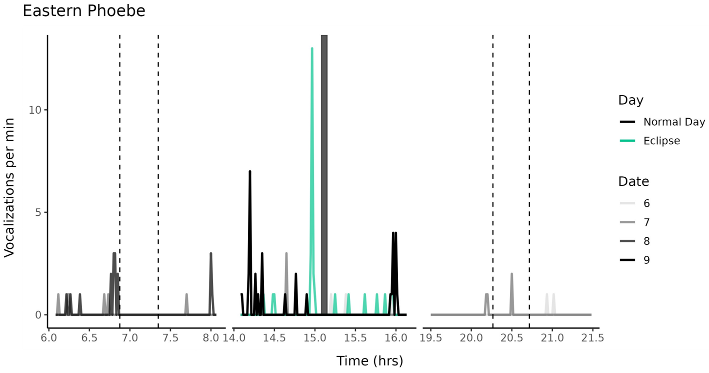
Eastern Phoebe
Vocalizations per min
Time (hrs)
Day
Normal Day
Eclipse
Date
6
7
8
9

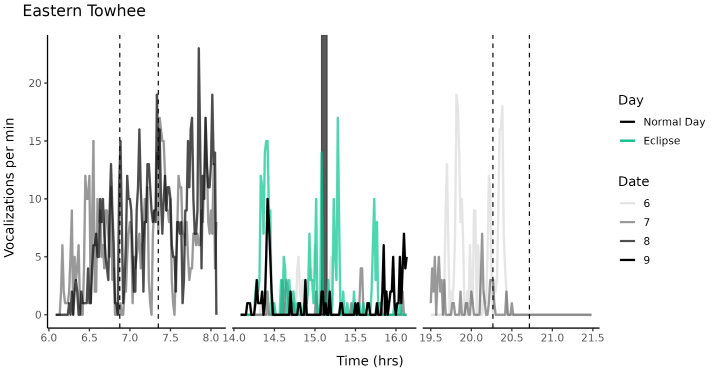
Eastern Towhee
Vocalizations per min
Time (hrs)
Day
Normal Day
Eclipse
Date
6
7
8
9

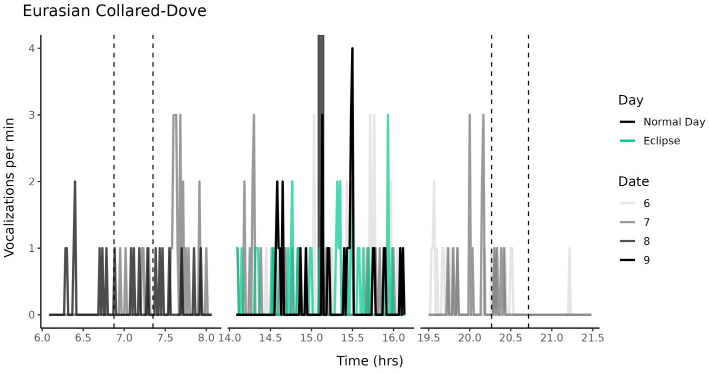
Eurasian Collared-Dove
Vocalizations per min
Time (hrs)
Day
Normal Day
Eclipse
Date
6
7
8
9

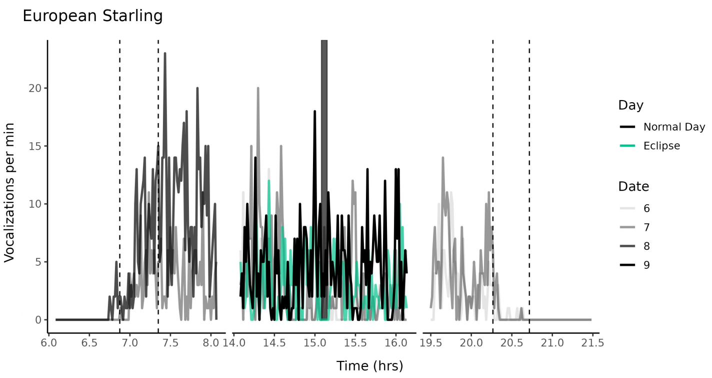
European Starling
Vocalizations per min
Time (hrs)
Day
Normal Day
Eclipse
Date
6
7
8
9

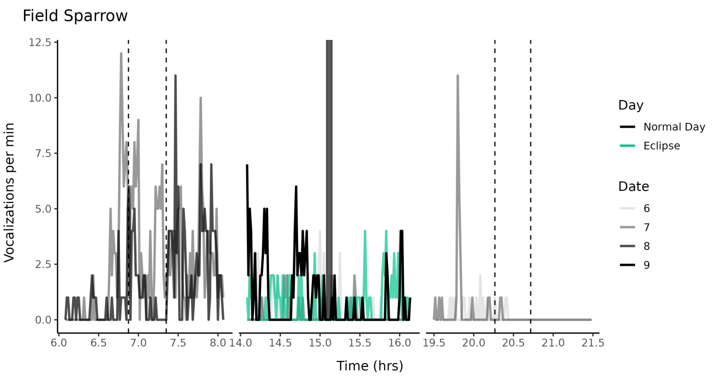
Field Sparrow
Vocalizations per min
Time (hrs)
Day
Normal Day
Eclipse
Date
6
7
8
9

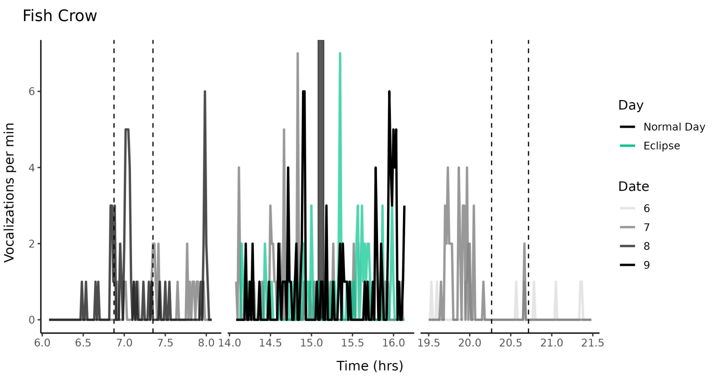
Fish Crow
Vocalizations per min
Time (hrs)
Day
Normal Day
Eclipse
Date
6
7
8
9

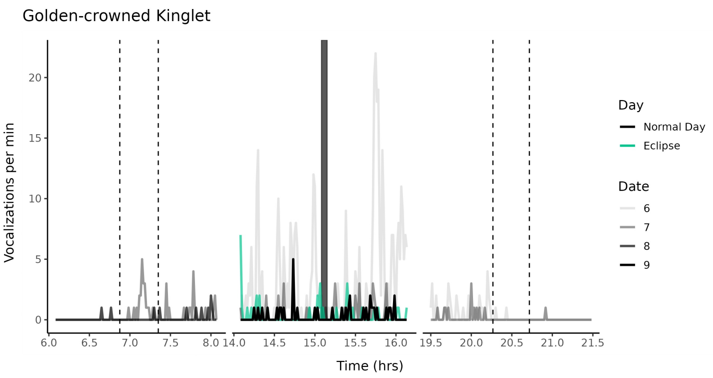
Golden-crowned Kinglet
Vocalizations per min
Time (hrs)
Day
Normal Day
Eclipse
Date
6
7
8
9

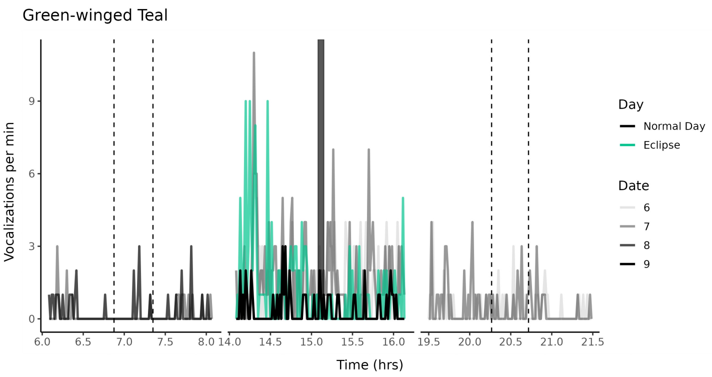
Green-winged Teal
Vocalizations per min
Time (hrs)
Day
Normal Day
Eclipse
Date
6
7
8
9

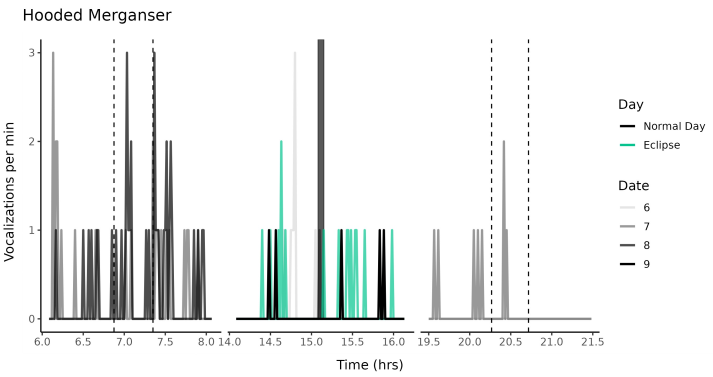
Hooded Merganser
Vocalizations per min
Time (hrs)
Day
Normal Day
Eclipse
Date
6
7
8
9

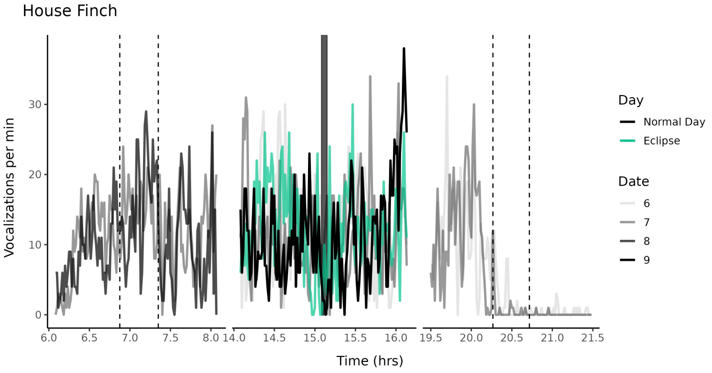
House Finch
Vocalizations per min
Time (hrs)
Day
Normal Day
Eclipse
Date
6
7
8
9

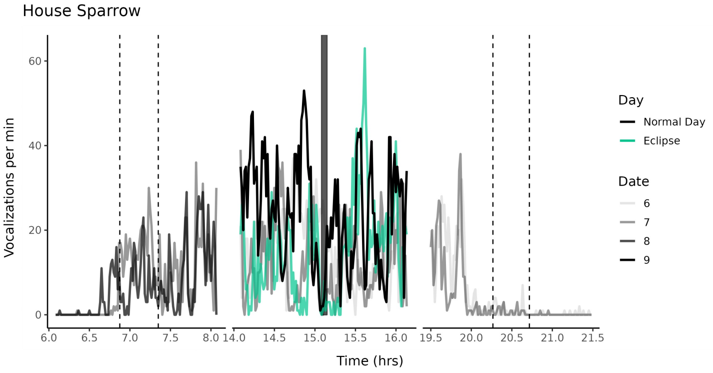
House Sparrow
Vocalizations per min
Time (hrs)
Day
Normal Day
Eclipse
Date
6
7
8
9

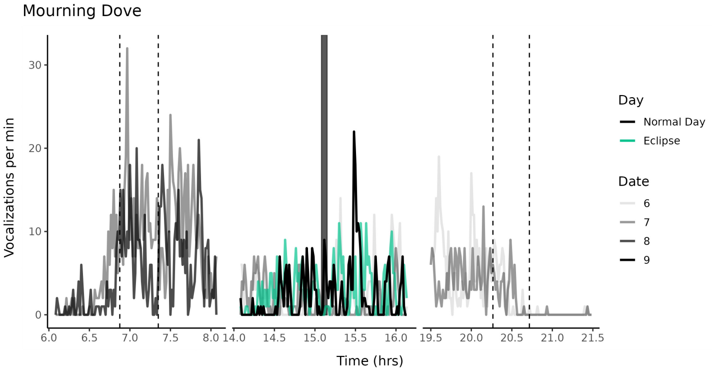
Mourning Dove
Vocalizations per min
Time (hrs)
Day
Normal Day
Eclipse
Date
6
7
8
9

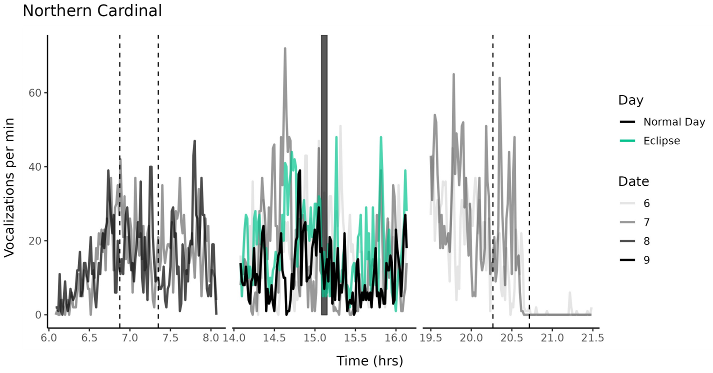
Northern Cardinal
Vocalizations per min
Time (hrs)
Day
Normal Day
Eclipse
Date
6
7
8
9

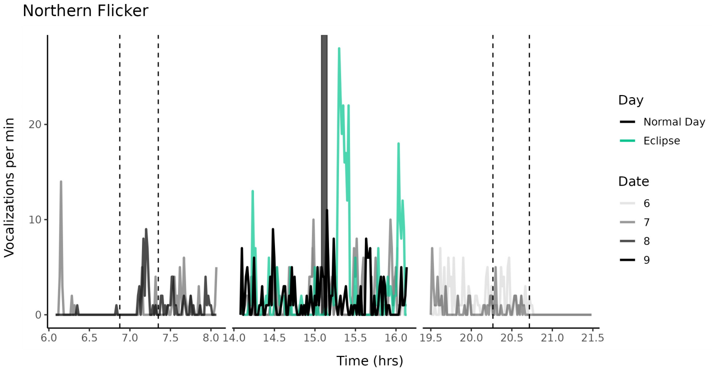
Northern Flicker
Vocalizations per min
Time (hrs)
Day
Normal Day
Eclipse
Date
6
7
8
9

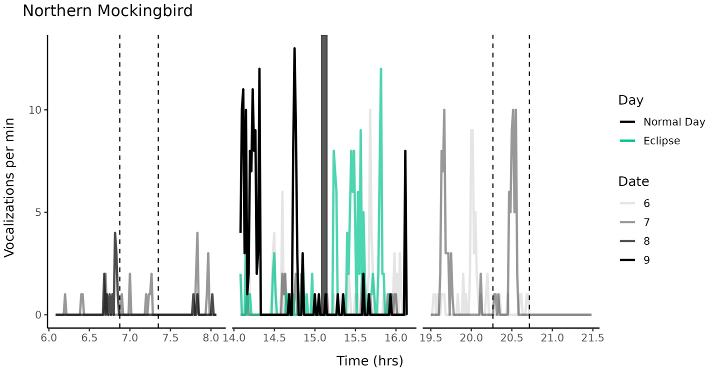
Northern Mockingbird
Vocalizations per min
Time (hrs)
Day
Normal Day
Eclipse
Date
6
7
8
9

Northern Rough-winged Swallow

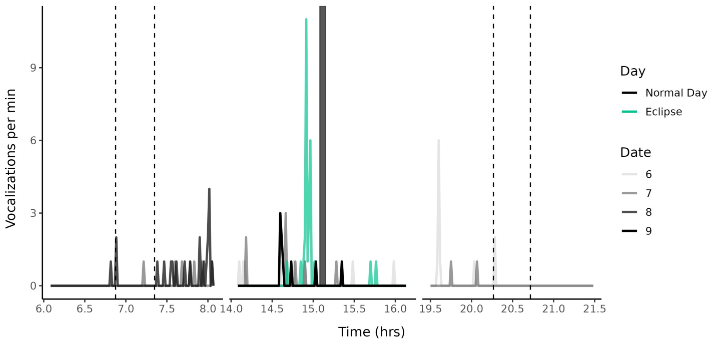


Pileated Woodpecker

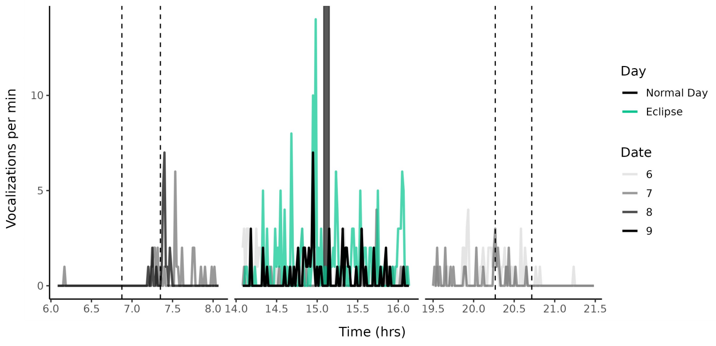


Pine Siskin

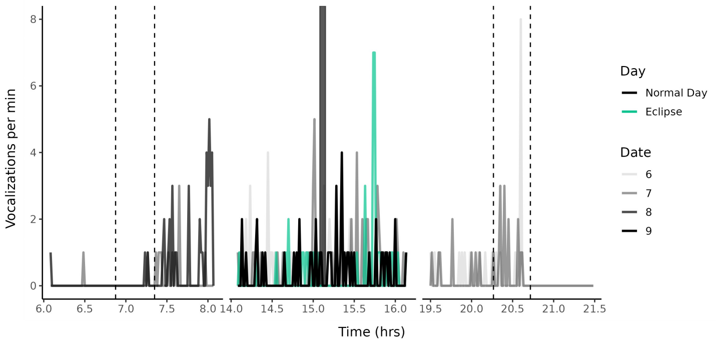


Purple Finch

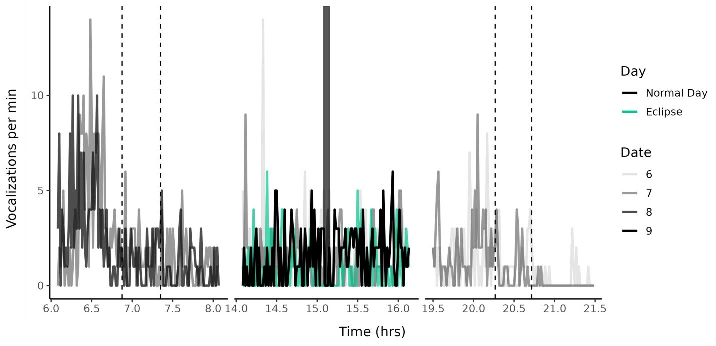


Purple Martin

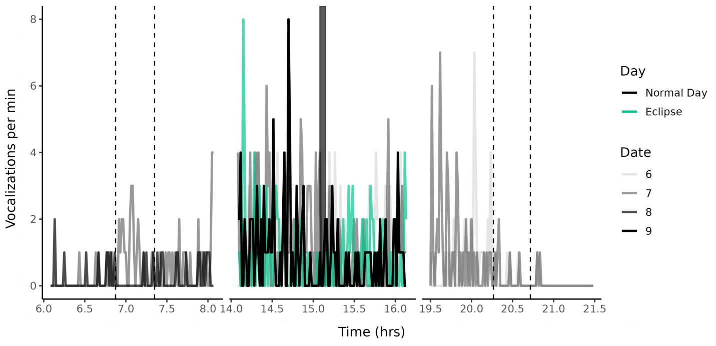


Red-bellied Woodpecker

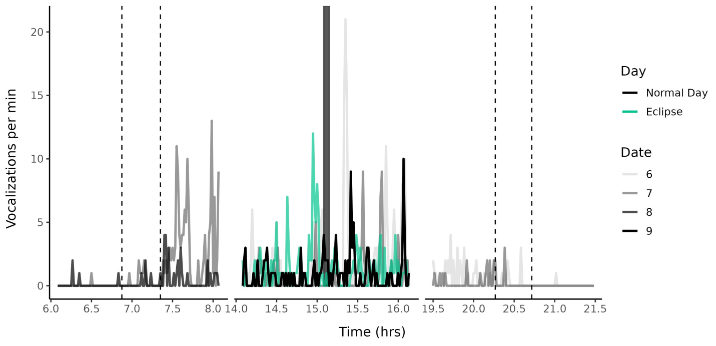

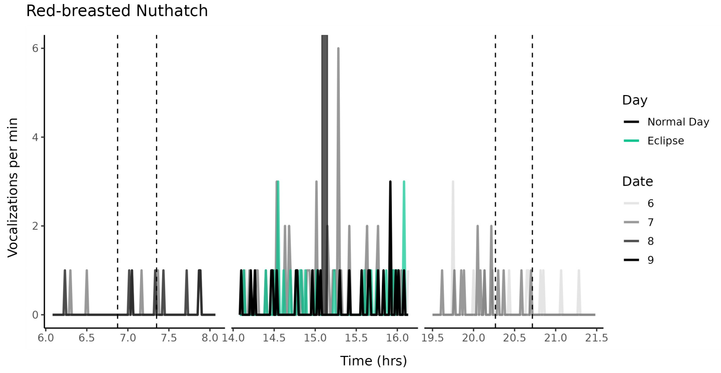
Red-breasted Nuthatch
Vocalizations per min
Time (hrs)
Day
Normal Day
Eclipse
Date
6
7
8
9

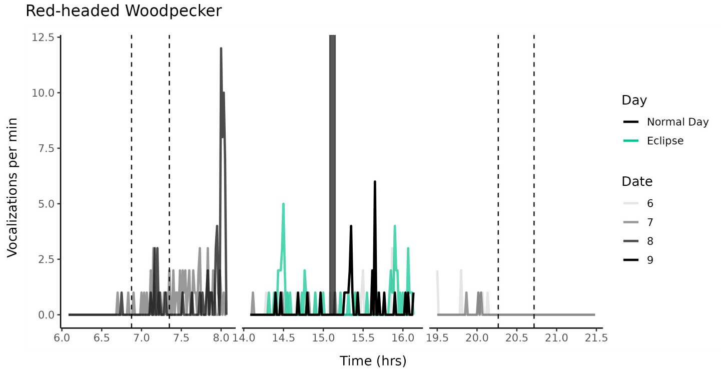
Red-headed Woodpecker
Vocalizations per min
Time (hrs)
Day
Normal Day
Eclipse
Date
6
7
8
9

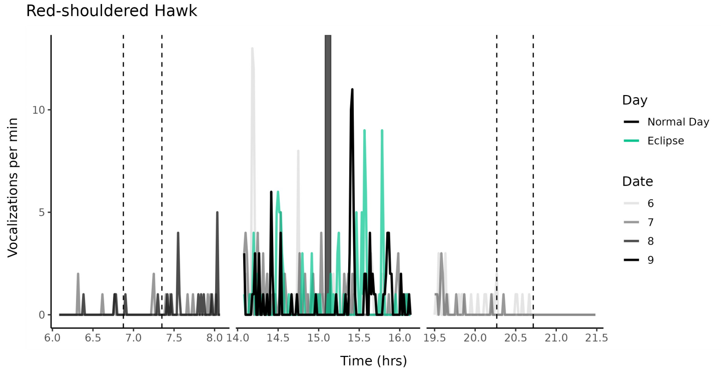
Red-shouldered Hawk
Vocalizations per min
Time (hrs)
Day
Normal Day
Eclipse
Date
6
7
8
9

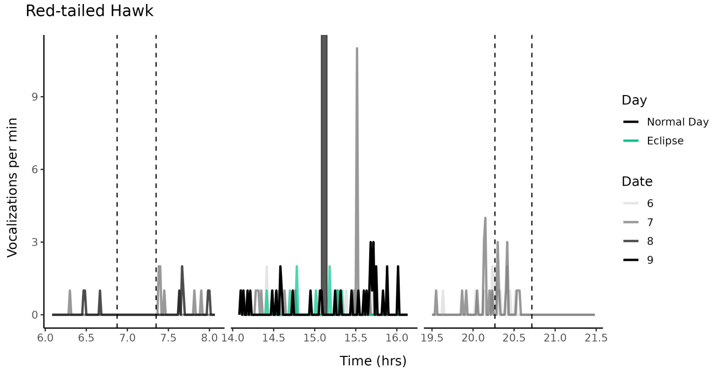
Red-tailed Hawk
Vocalizations per min
Time (hrs)
Day
Normal Day
Eclipse
Date
6
7
8
9

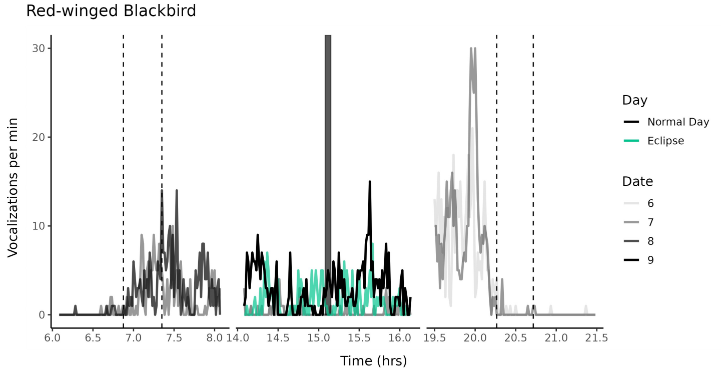
Red-winged Blackbird
Vocalizations per min
Time (hrs)
Day
Normal Day
Eclipse
Date
6
7
8
9

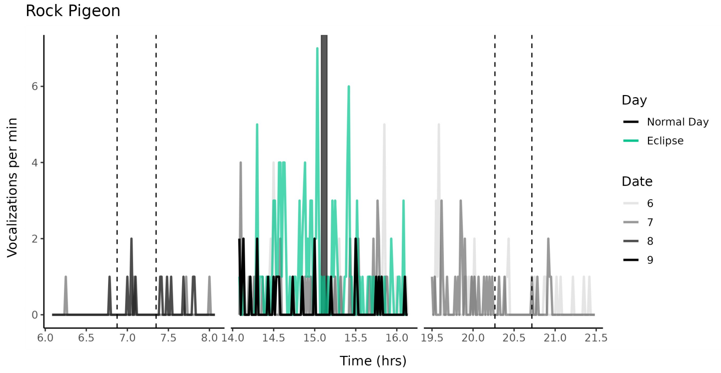
Rock Pigeon
Vocalizations per min
Time (hrs)
Day
Normal Day
Eclipse
Date
6
7
8
9

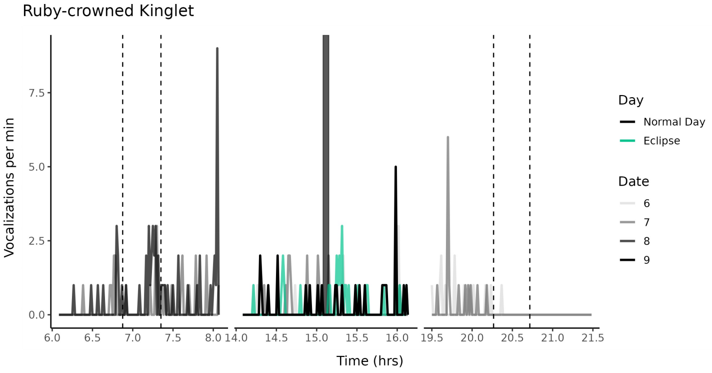
Ruby-crowned Kinglet
Vocalizations per min
Time (hrs)
Day
Normal Day
Eclipse
Date
6
7
8
9

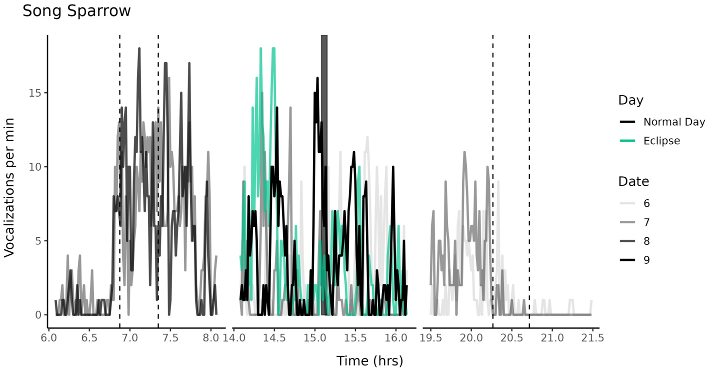
Song Sparrow
Vocalizations per min
Time (hrs)
Day
Normal Day
Eclipse
Date
6
7
8
9

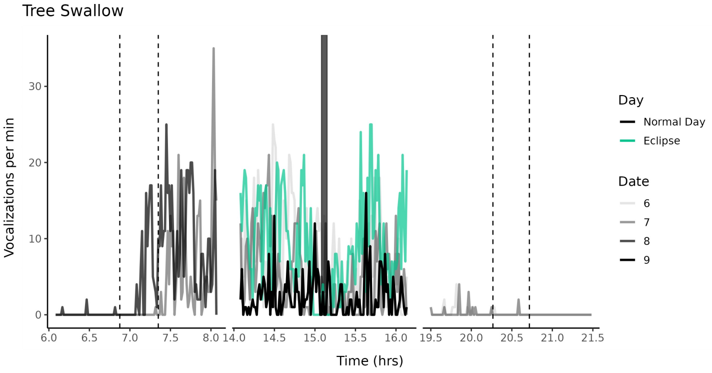
Tree Swallow
Vocalizations per min
Time (hrs)
Day
Normal Day
Eclipse
Date
6
7
8
9

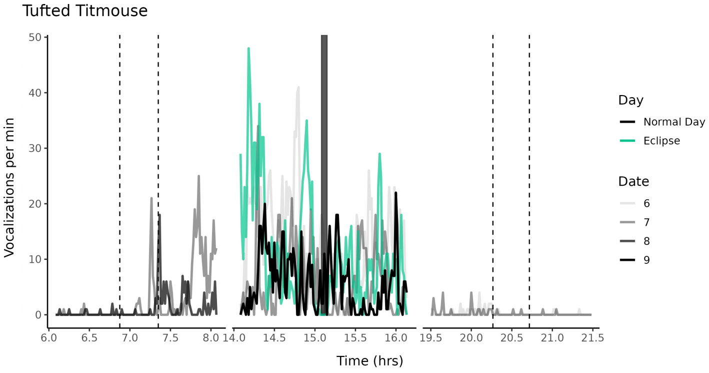
Tufted Titmouse
Vocalizations per min
Time (hrs)
Day
Normal Day
Eclipse
Date
6
7
8
9

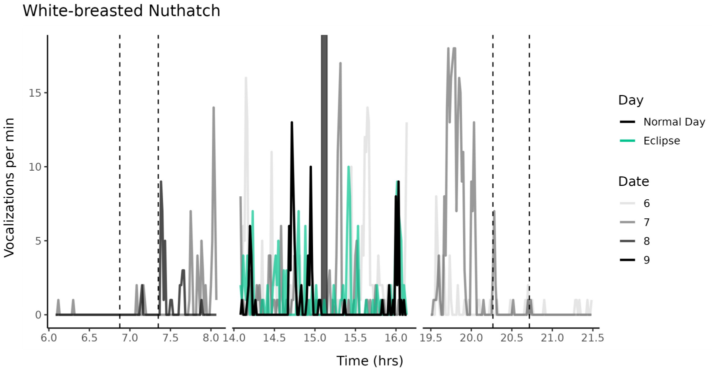
White-breasted Nuthatch
Vocalizations per min
Time (hrs)
Day
Normal Day
Eclipse
Date
6
7
8
9

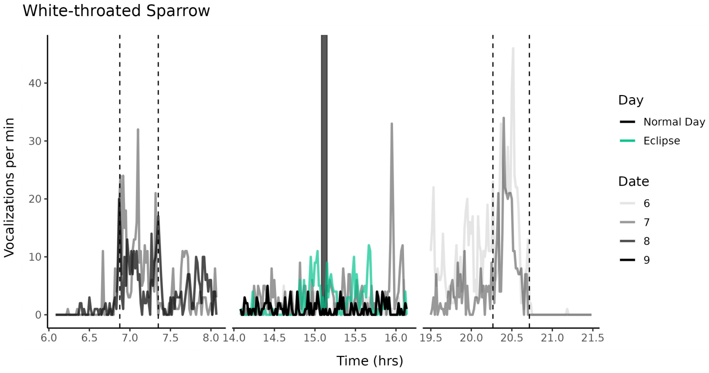
White-throated Sparrow
Vocalizations per min
Time (hrs)
Day
Normal Day
Eclipse
Date
6
7
8
9

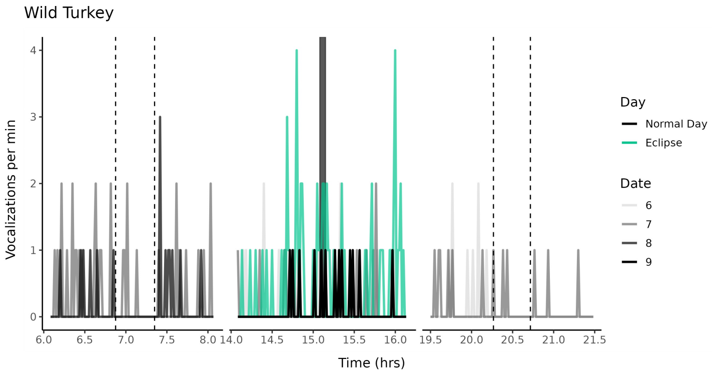
Wild Turkey
Vocalizations per min
Time (hrs)
Day
Normal Day
Eclipse
Date
6
7
8
9

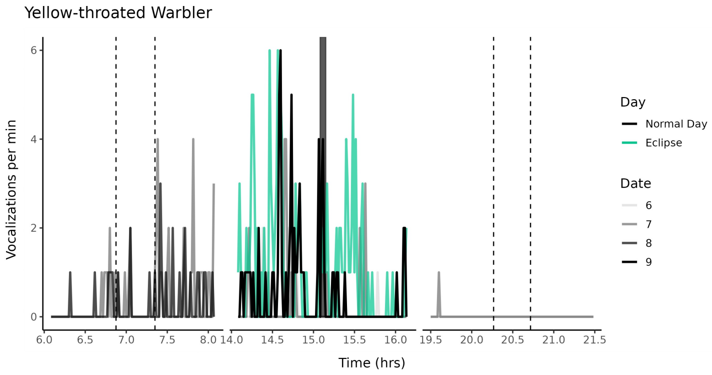
Yellow-throated Warbler
Vocalizations per min
Time (hrs)
Day
Normal Day
Eclipse
Date
6
7
8
9

**Fig. S3. Vocalizations detected, by species across focal days.** For each species, we show the rate of vocalization using a 1-min moving sum across the three core periods of analysis, including the two hours surrounding sunrise, sunset, and the eclipse. Dashed lines denote major natural shifts in light, from left to right: civil twilight in the morning, sunrise, sunset, and civil twilight in the evening. The shaded bar at 15:04:55 marks the period of totality.

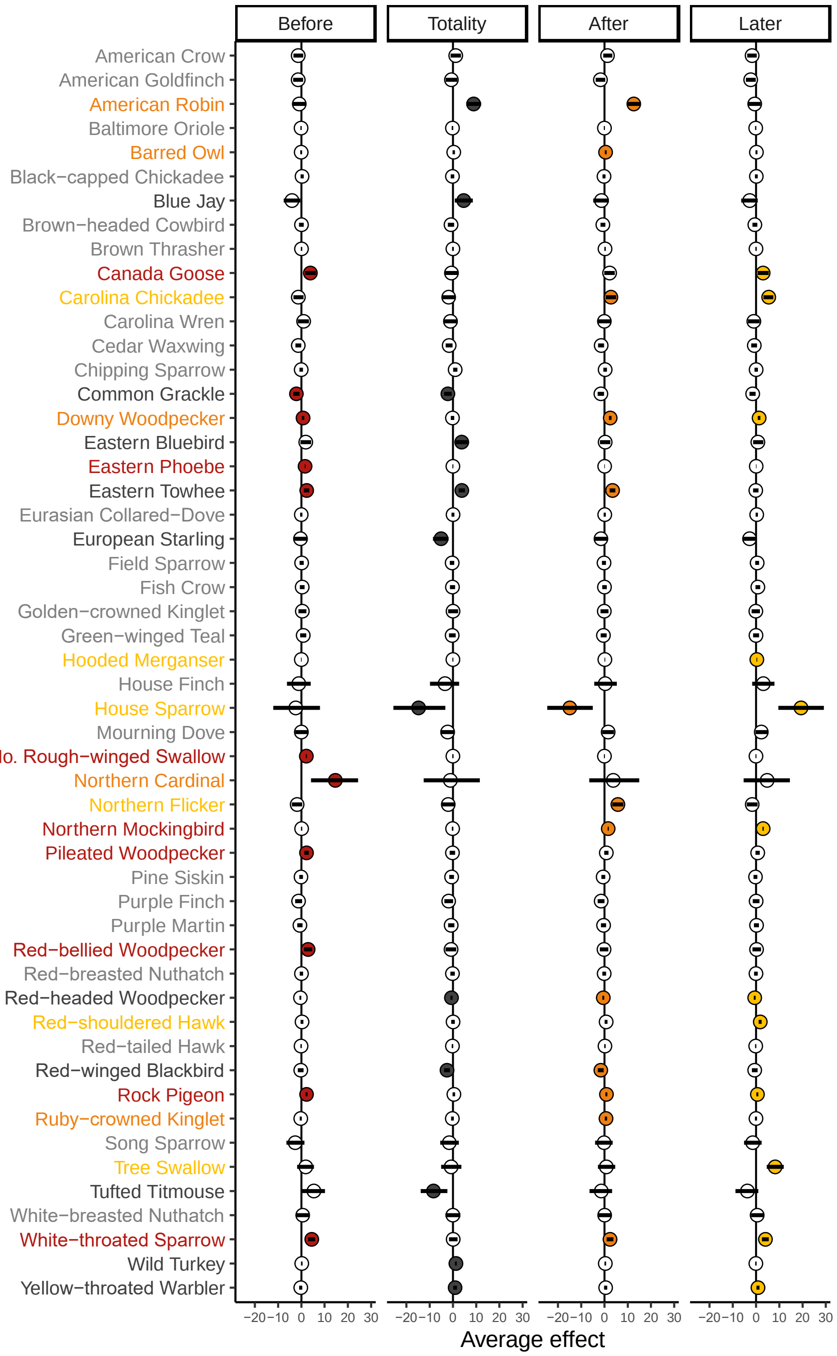


**Fig. S4. Forest plot for all 52 species.** Species are sorted alphabetically by common name. Each point represents an effect size (scaled per minute) with 95% CI. Species names and points are colored if significant during one of four time periods: 12 min just before totality ("Before", red), 4 min during totality ("Totality", black), 12 min just after totality ("After", orange), and 12 min slightly later ("Later", yellow). Light gray denotes species that were not significantly affected in any of these sampling periods.

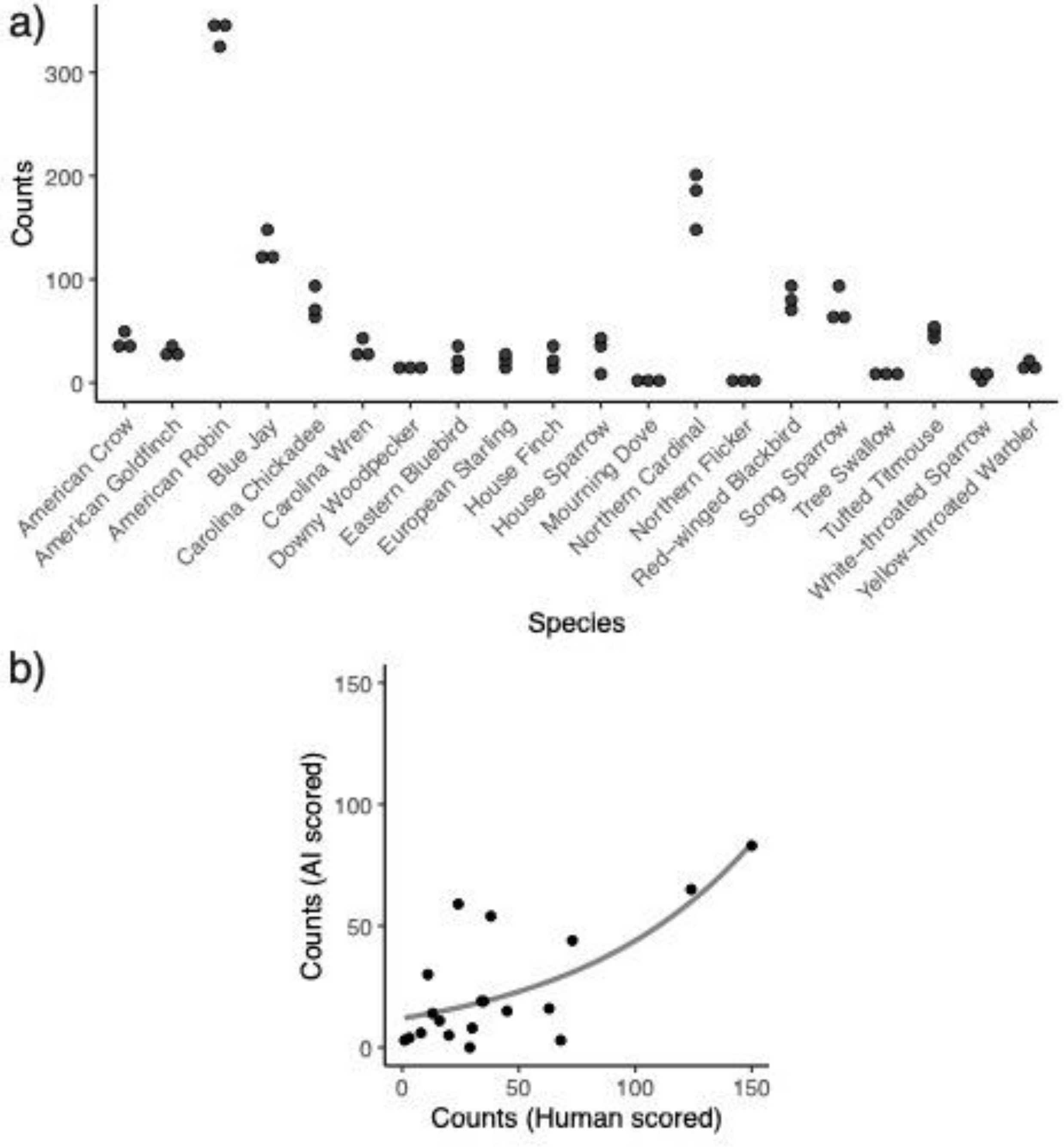


**Fig. S5. Validations by expert birder.** Human-scored vocalization counts from an 8-minute period during the eclipse (4 minutes of totality, 4 minutes immediately after) were (a) highly repeatable across a focal subset of species, and (b) significantly correlated with the AI-derived counts from the same period (GLM: $t(18) = 3.588$, $p = 0.002$, $d^2 = 0.419$). American Robin was removed from (b) and the analysis as an outlier (see Materials and Methods for details). Data were derived from 13 of 14 autonomous recording units. One recorder was not scored by the human due to potential privacy issues from a nearby human conversation.

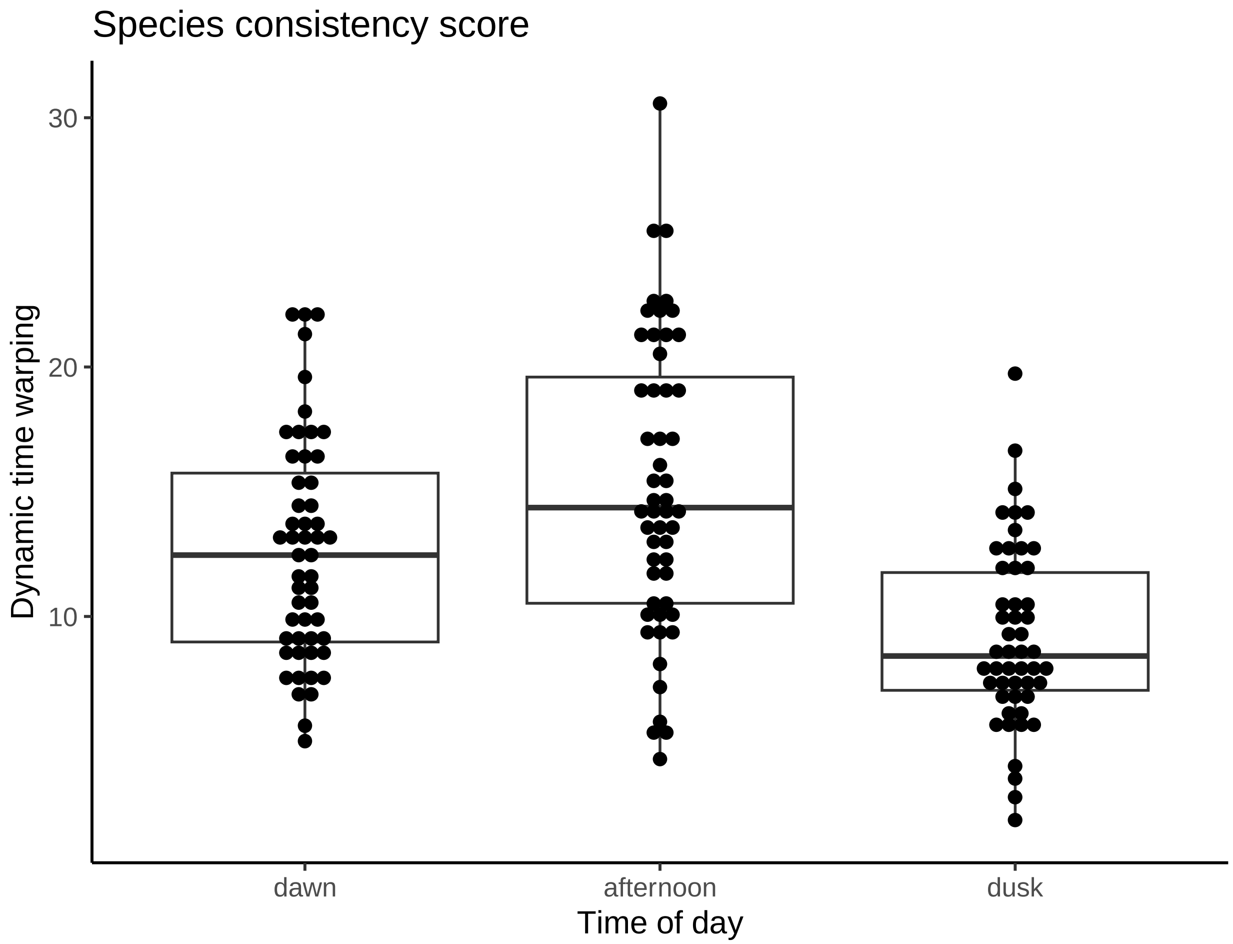


**Fig. S6. Consistency in vocalization surrounding dawn, dusk, and afternoon.** Consistency scores per species as measured by dynamic time warping score for the three time-windows across the day (i.e., dawn, afternoon, dusk, see Materials & Methods). A *lower* score indicates vocalization temporal dynamics that are more similar across days. Afternoon scores represent average across pairwise comparisons of April 6, 7, and 9. The central line of the boxplots identifies the median, the box encompasses the interquartile range, and the whiskers show the range of data within 1.5 times the interquartile range.

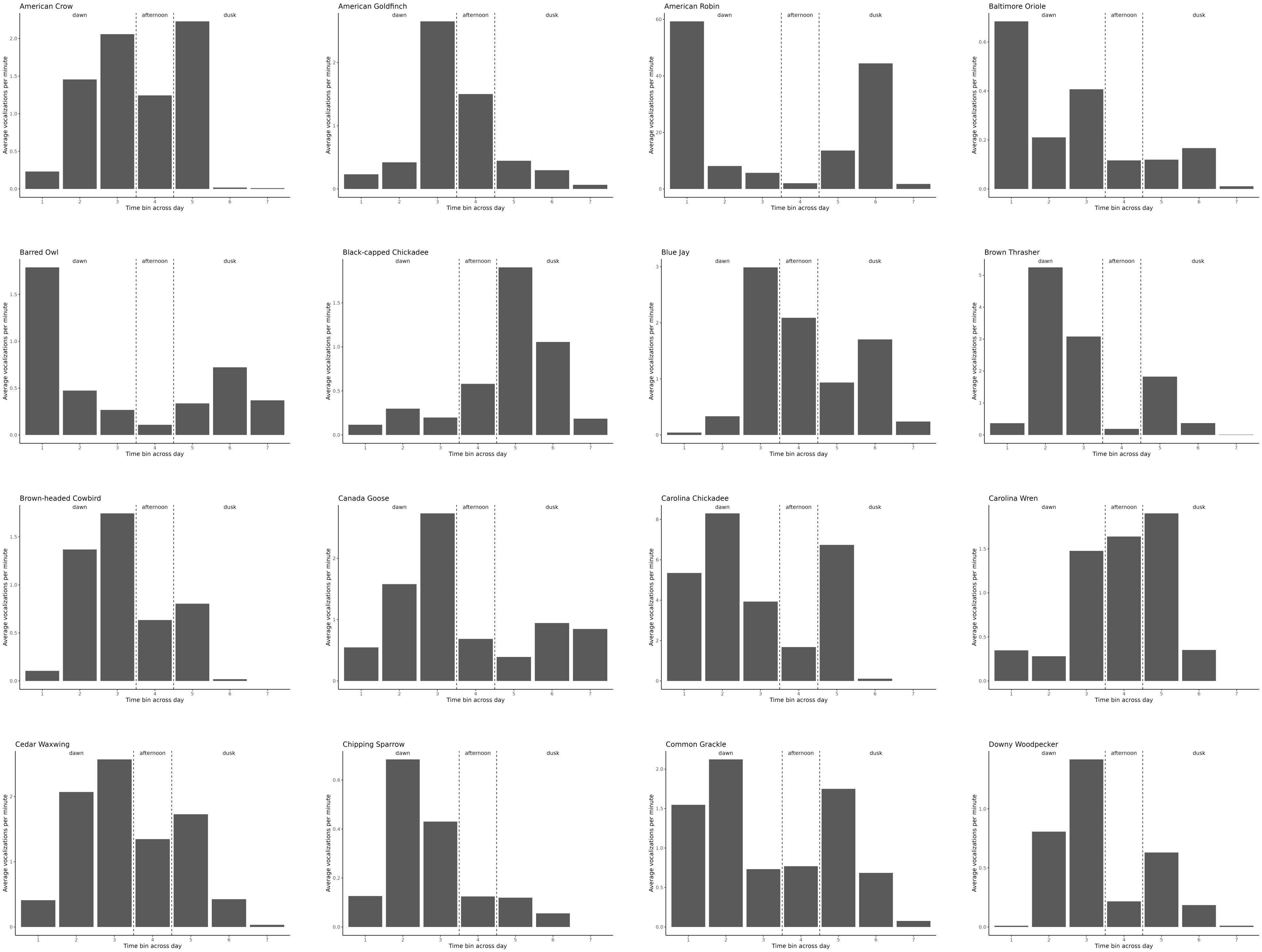
American Crow
American Goldfinch
American Robin
Baltimore Oriole
Barred Owl
Black-capped Chickadee
Blue Jay
Brown Thrasher
Brown-headed Cowbird
Canada Goose
Carolina Chickadee
Carolina Wren
Cedar Waxwing
Chipping Sparrow
Common Grackle
Downy Woodpecker
dawn
afternoon
dusk
Average vocalizations per minute
Time bin across day

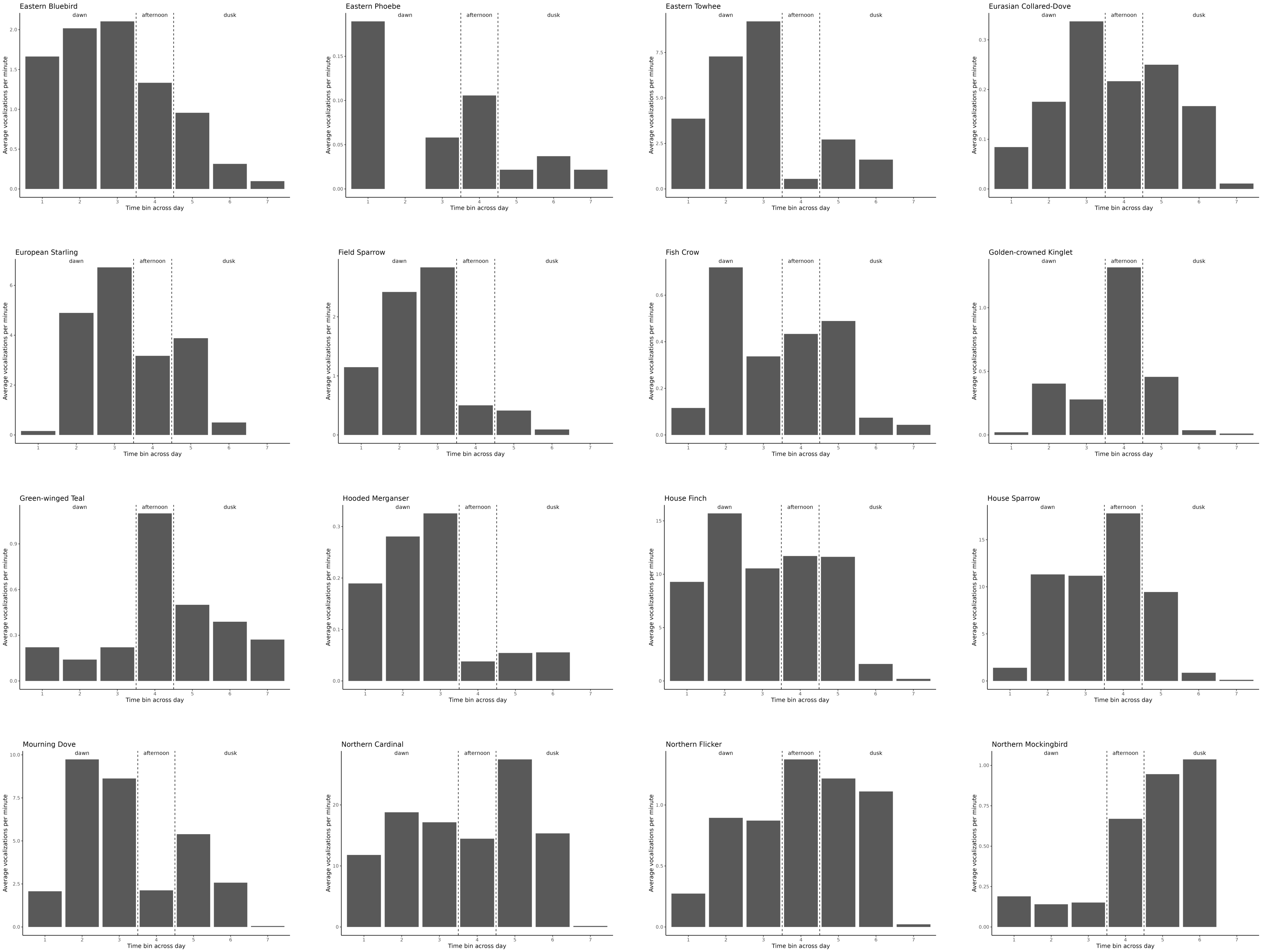
Eastern Bluebird
Eastern Phoebe
Eastern Towhee
Eurasian Collared-Dove
European Starling
Field Sparrow
Fish Crow
Golden-crowned Kinglet
Green-winged Teal
Hooded Merganser
House Finch
House Sparrow
Mourning Dove
Northern Cardinal
Northern Flicker
Northern Mockingbird
dawn
afternoon
dusk
Average vocalizations per minute
Time bin across day

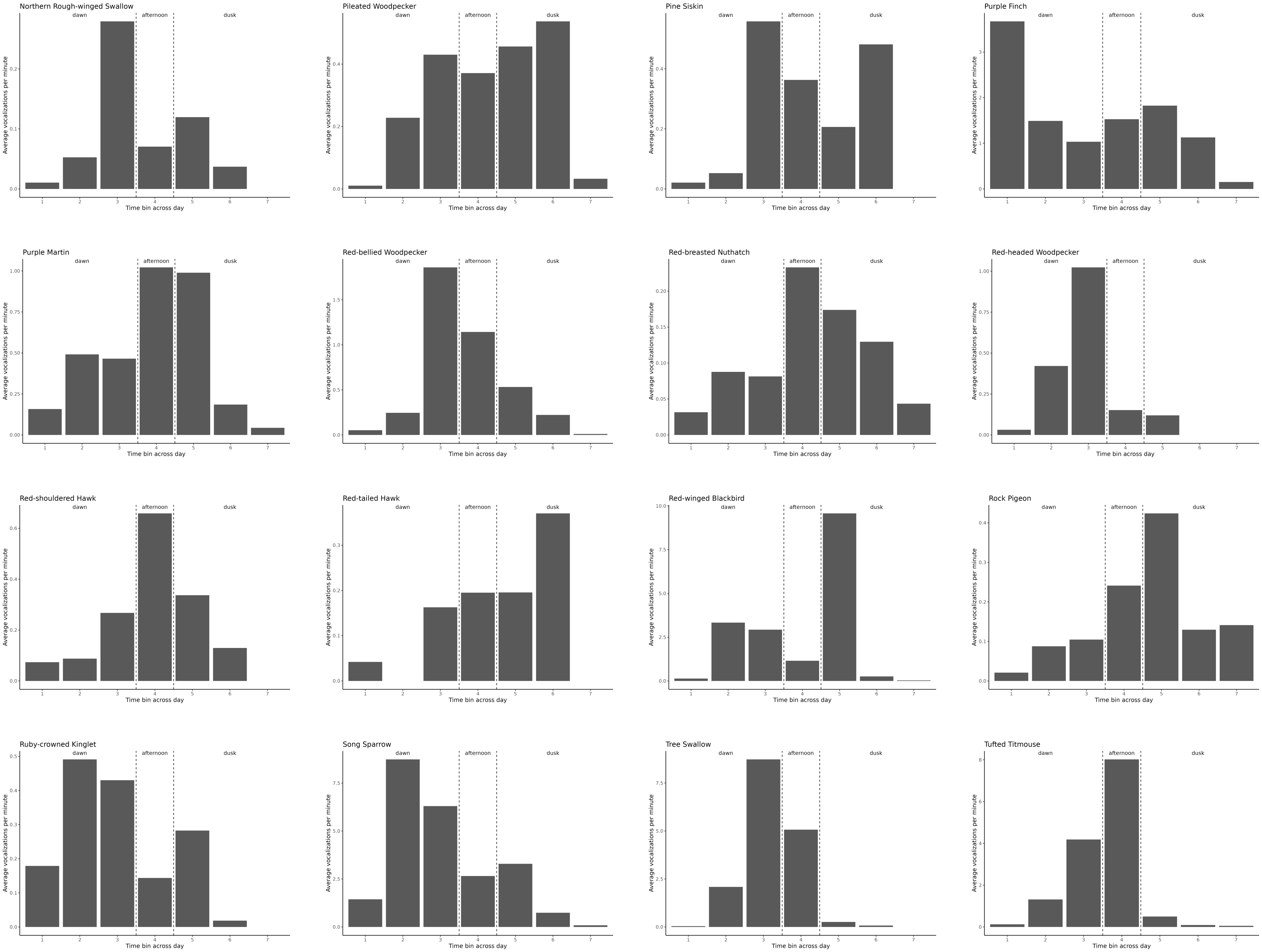
Northern Rough-winged Swallow
Pileated Woodpecker
Pine Siskin
Purple Finch
Purple Martin
Red-bellied Woodpecker
Red-breasted Nuthatch
Red-headed Woodpecker
Red-shouldered Hawk
Red-tailed Hawk
Red-winged Blackbird
Rock Pigeon
Ruby-crowned Kinglet
Song Sparrow
Tree Swallow
Tufted Titmouse
dawn
afternoon
dusk
Average vocalizations per minute
Time bin across day

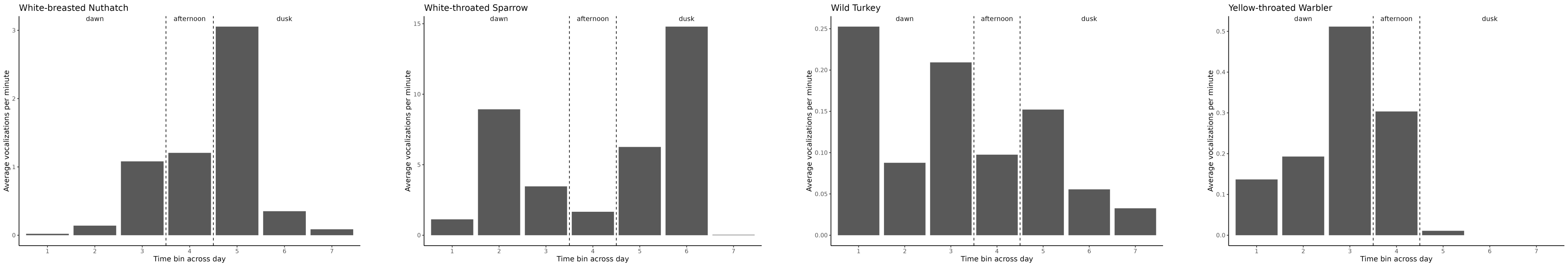
White-breasted Nuthatch
White-throated Sparrow
Wild Turkey
Yellow-throated Warbler
dawn
afternoon
dusk
Average vocalizations per minute
Time bin across day

**Fig. S7. Per minute vocalization rate across seven time periods in normal day.** Average vocalizations per minute across normal days for each species across time periods in relation to sunrise and sunset. Time periods were categorized as dawn (1,2,3); afternoon (4); and dusk (5,6,7). See Materials and Methods for specific time windows that encompass these seven periods.

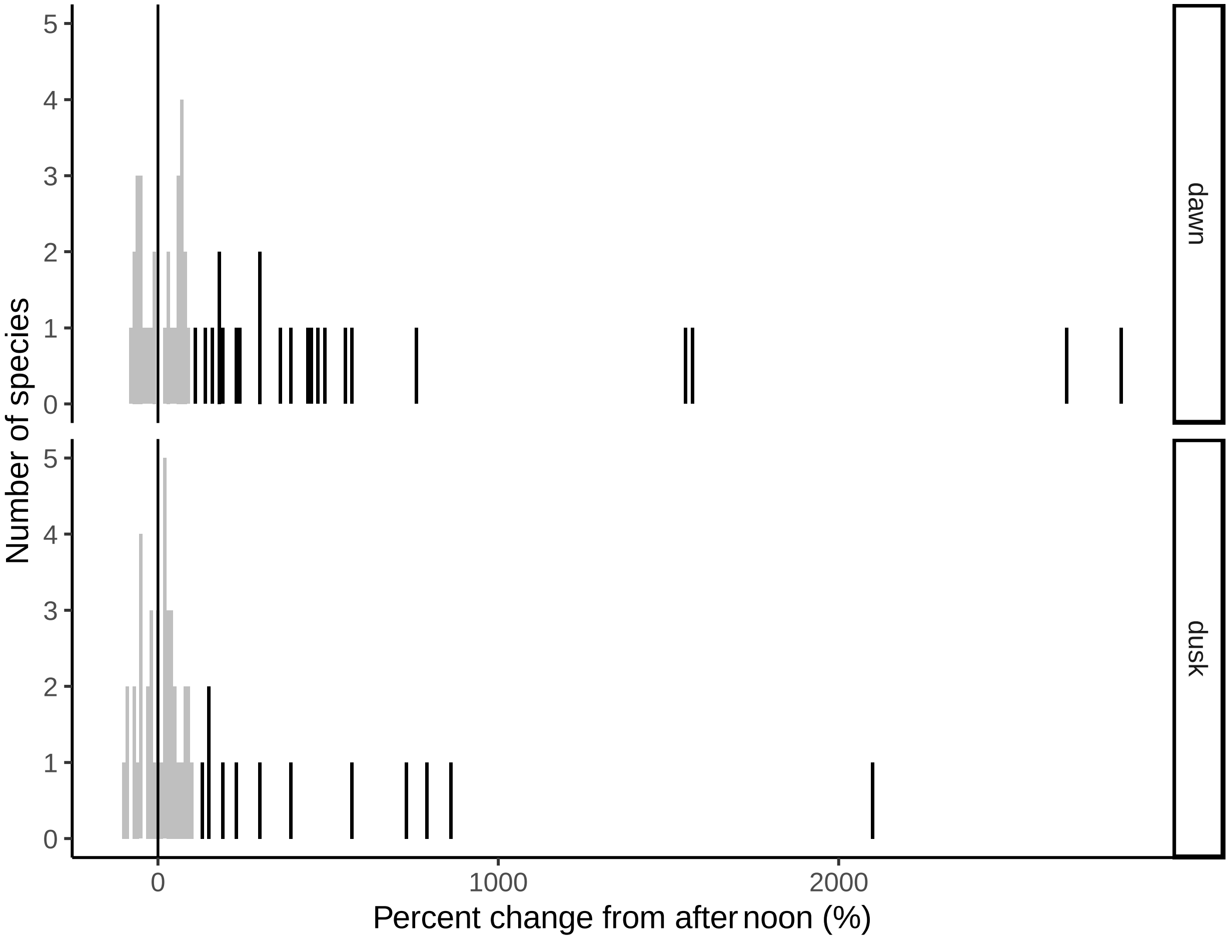


**Fig. S8. Dawn and dusk chorus threshold.** Species with a percent change greater than 100% in average number of vocalizations per minute across normal days for one time-period in dawn (top) or dusk (bottom) are considered to have a dawn or dusk chorus, respectively, and are colored black.

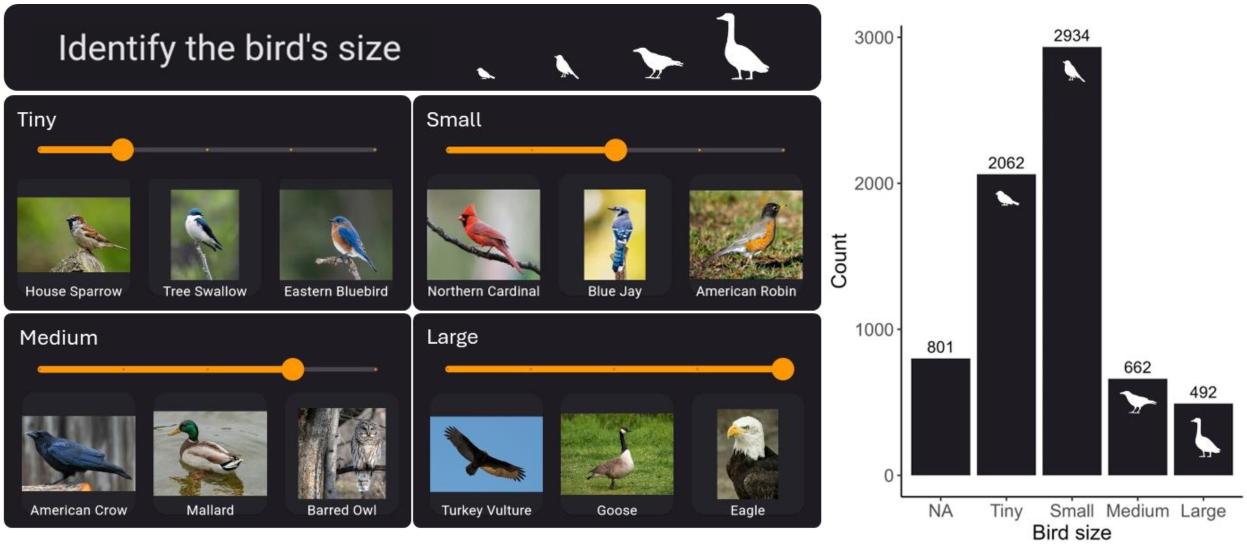


**Fig. S9. *SolarBird* app entries on bird size.** App participants could choose from four options to identify the size of the bird they were observing. Most users reported observing a small bird, akin to a northern cardinal, blue jay, or American robin.

| | Vocalizing | Flying | Stationary | Other |
|---|---|---|---|---|
| Before vs After | 0.002 | 0.003 | 0.093 | 0.987 |
| Totality vs After | 0.030 | < 0.001 | < 0.001 | < 0.001 |
| Totality vs Before | < 0.001 | < 0.001 | < 0.001 | < 0.001 |

**Table S1. Exact p-values from *SolarBird* behavioral observations.** App data showed that behaviors varied in their frequency, before ("Before), during ("Totality"), and after ("After") totality (binomial general linear model with Tukey post-hoc analysis and FDR correction).

| Keystroke | Keystroke meaning | Retained for AI validation? |
|---|---|---|
| 0 | downy woodpecker | X |
| 1 | red shouldered hawk | |
| 2 | red tailed hawk | |
| 3 | sharp shinned hawk | |
| 4 | coopers hawk | |
| 8 | pileated woodpecker | |
| 9 | northern flicker | X |
| a | american crow | X |
| b | brown-headed cowbird | |
| c | northern cardinal | X |
| d | mourning dove | X |
| e | european starling | X |
| f | house finch | X |
| g | american goldfinch | X |
| h | house sparrow | X |
| j | blue jay | X |
| k | carolina chickadee | X |
| l | eastern bluebird | X |
| m | yellow-throated warbler | X |
| n | nuthatch, white breasted | |
| q | tree swallow | X |
| r | american robin | X |
| s | song sparrow | X |
| t | tufted titmouse | X |
| u | cedar waxwing | |
| v | red-winged blackbird | X |
| w | carolina wren | X |
| z | white-throated sparrow | X |
| ` | owl (any species) | |
| 5 | other raptor | |
| 7 | other woodpecker | |
| x | other sparrow (junco, towhee, field, chipping, etc.) | |
| y | shorebirds (geese, ducks, etc) | |
| / | OTHER bird | |
| [ | NATURAL NOISE (e.g. wind, rain) | |
| o | HUMAN VOICE | |
| p | OTHER ANTHROPOGENIC NOISE (e.g. mower, traffic) | |
| ] | CANT SCORE (other interference or distortion) | |
| i | UNKNOWN (hear a bird, but can't tell what it is) | |

**Table S2. JWatcher focal species and keystrokes.** An expert birder scored excerpts of audio files by ear for 36 possible species or species groups, recording vocalizations with unique keystrokes. Twenty of those species were observed by the human and used to validate vocalization counts by the AI.
